# CLASH (Chromatin Loop Across-sample Score Harmonizer) quantifies the relative contributions of genetic variation, methylation, and CTCF occupancy on chromatin loop strength across individuals

**DOI:** 10.64898/2026.06.01.729143

**Authors:** Valmik Ranparia, The Human Genome Structural Variation Consortium, Geoffrey Fudenberg, Mark JP Chaisson

**Affiliations:** Department of Quantitative and Computational Biology, University of Southern California, CA, USA; Norris Comprehensive Cancer Center, University of Southern California, CA, USA

## Abstract

Three-dimensional genome organization constrains the regulatory interactions that govern vital cellular processes. Chromatin loops are key features of genome folding, yet it is unclear how genetic and epigenetic variation influences differential loop formation across individuals. Loops primarily form between two CTCF binding proteins, which recognize a specific motif at loop anchors. CTCF binding site motifs are frequently altered by base substitutions, structural variation, and 5-methylcytosine (m⁵C) CpG methylation, yet no study has comprehensively profiled this variation across diverse individuals. Moreover, existing approaches relying on binary loop calls fail to capture subtle changes in genetic and epigenetic features, as well as CTCF occupancy, that drive variation in loop strength. Here, we combined high-resolution Hi-C, Fiber-seq, near telomere-to-telomere phased assemblies, and m⁵C methylation maps across five lymphoblastoid cell lines to quantify how genetic and epigenetic variation shape genome folding. We used DiffHiC to identify 367 differential pixels and found that sequence variation, chromatin accessibility, and m⁵C CpG methylation are each significantly associated with differential chromatin contacts. Next, we developed CLASH (Chromatin Loop Across-sample Score Harmonizer) to harmonize loop calls across samples and enable robust comparisons of loop strengths across individuals. CLASH substantially improved loop calls and loop score calibration with respect to the classification boundary over existing methods and confirmed a significant relationship between CTCF occupancy and loop strength. We then characterized independent contributions of sequence and epigenetic variation to differential loop formation, demonstrating that 57% of sequence variation- and 40% of methylation-associated effects on loop formation acted through CTCF occupancy. Together, we present a multimodal dataset and computational approach to facilitate the study of 3D genome structure across human populations.

## Introduction

In eukaryotic cells, the three-dimensional spatial organization of the genome regulates vital cellular processes such as transcription and replication. Altered genome folding has been implicated in conditions ranging from polydactyly (Paliou et al. 2019), Cornelia de Lange Syndrome (Panarotto et al. 2022), and some cancers (Yoon et al. 2024; Glushakow-Smith and Tothova 2025). Three primary genome folding features are visible in interphase Hi-C maps: A/B compartments, topologically associating domains (TADs), and chromatin loops (McCord et al. 2020). Chromatin loop topology differs across cell types (Bond et al. 2023; Grubert et al. 2020; Burren et al. 2017), developmental stages (Li et al. 2019; Siersbæk et al. 2017), and between individuals sharing the same cell state (Greenwald et al. 2019; Li et al. 2024).

Loop formation and chromatin architecture reflect dynamic interactions of DNA binding proteins including CTCF and the cohesin complex (van Ruiten and Rowland 2021), modulated by methylation, chromatin accessibility, and genetic variation. Early studies reported effects of methylation (Wang et al. 2012) and sequence variation (Maurano et al. 2012) on CTCF binding, followed by studies characterizing differential 3D genome structure across cell (epigenetic) states (Grubert et al. 2020; Monteagudo-Sánchez et al. 2024; Pękowska et al. 2018). More recently, interaction quantitative trait loci (iQTLs) identified using Hi-C sequencing from chromatin immunoprecipitation (HiChIP; (Bhattacharyya and Ay 2024) were associated with expression quantitative trait loci (eQTLs) – though HiChIP signal can also reflect changes in the target ChIP protein positioning. Additionally, the effect of structural variation (SVs; insertions, deletions, and rearrangements ≥ 50 bases) on chromatin structure was characterized across high-quality reference genomes assembled by the Human Genome Structural Variation Consortium (HGSVC) (Li et al. 2024). Despite these insights, quantifying the independent contributions of genetic and epigenetic variation to altered folding has remained out of reach. A more detailed understanding of how chromatin architecture and loop formation differs between individuals is therefore crucial for understanding how variation influences traits and disease.

Addressing this gap requires new datasets and new computational approaches. Advancements in sequencing technologies have enabled multi-modal, comprehensive analysis of genetic and epigenetic variation via telomere-to-telomere (T2T) genomes assembled using long read sequencing (LRS) (Logsdon et al. 2025), phased methylation measured directly from LRS data (Simpson et al. 2017), and both CTCF and nucleosome occupancy derived from Fiber-seq profiles (Stergachis et al. 2020). Loop calling in conformation capture data presents an additional computational challenge: a study of 22 existing methods showed inconsistent loop calling (Chowdhury et al. 2024). Furthermore, although previous studies have indicated that subtle differences in loop strength are associated with changes in molecular phenotypes (Greenwald et al. 2019), existing methods focus primarily on classifying whether a loop is present or not. Consequently, loop comparisons between individuals have remained largely unexplored. The creation of a method that quantifies loop strength and that can harmonize calls within and between individuals would enable deeper insight into the relationship between genetic and epigenetic variation and genome structure.

Here, we present a multi-modal, haplotype-resolved map of genetic and epigenetic features for five diverse lymphoblastoid cell lines with near telomere-to-telomere (T2T) genomes. We use this resource to quantify the impact of genetic and epigenetic variation on three-dimensional genome structure, either through modulation of CTCF occupancy or other indirect factors. Analysis of loops on this integrated resource revealed a need to both harmonize loop calls between individuals and quantify fine-grained differences between them. To address this, we developed CLASH (Chromatin Loop Across-sample Score Harmonizer), a method that assigns harmonized loop-strength scores to candidate loops and enables robust inter-individual comparisons of loop strength. Using CLASH-scored chromatin loops from high-resolution Hi-C contact maps together with phased sequence variation, m^5^C methylation profiles, and single-molecule CTCF-occupancy measurements from Fiber-seq, we determined the independent contributions of CTCF occupancy, genetic variation, and epigenetic variation to differential loop formation.

## Results

### Construction of an integrated resource spanning genetic, epigenetic and genome folding variation

We generated an integrated resource of Hi-C and Fiber-seq to complement existing near-telomere-to-telomere (T2T) assembly datasets (Logsdon et al. 2025) and m⁵C CpG methylation calls from the HGSVC for five lymphoblastoid cell lines: GM19317, GM19347, HG01457, HG02666, and HG03248 (Logsdon et al. 2025)(Fig. 1a). The five genomes contained an average of 4.0M SNVs, 833k indels (< 50 bases), and 23.4k SVs (≥ 50 bases) per haplotype detected by the Phased Assembly Caller (Ebert et al. 2021). Hi-C samples yielded an average of ∼163.5x genomic coverage with equalized proportions of read-pair orientations at distances between 1–2 kb, supporting analyses at 2 kb resolution (Table S1). Fiber-seq data (Stergachis et al. 2020) were generated using PacBio HiFi sequencing with a mean coverage of 33.5× per genome (Table S2), enabling haplotype-resolved annotation of chromatin accessibility. The full computational pipeline we used is summarized in Fig. 1b. All samples exhibited high sign concordance for both Hi-C A/B compartment profiles (∼95%; Supplementary Figures 1, 2) and Fiber-seq mean-centered accessibility profiles (∼83%; Supplementary Figure 1), confirming that all samples are in the same cell state and indicating that compartment-level differences do not confound subsequent analyses of contacts and loop formation.

**Figure 1.**
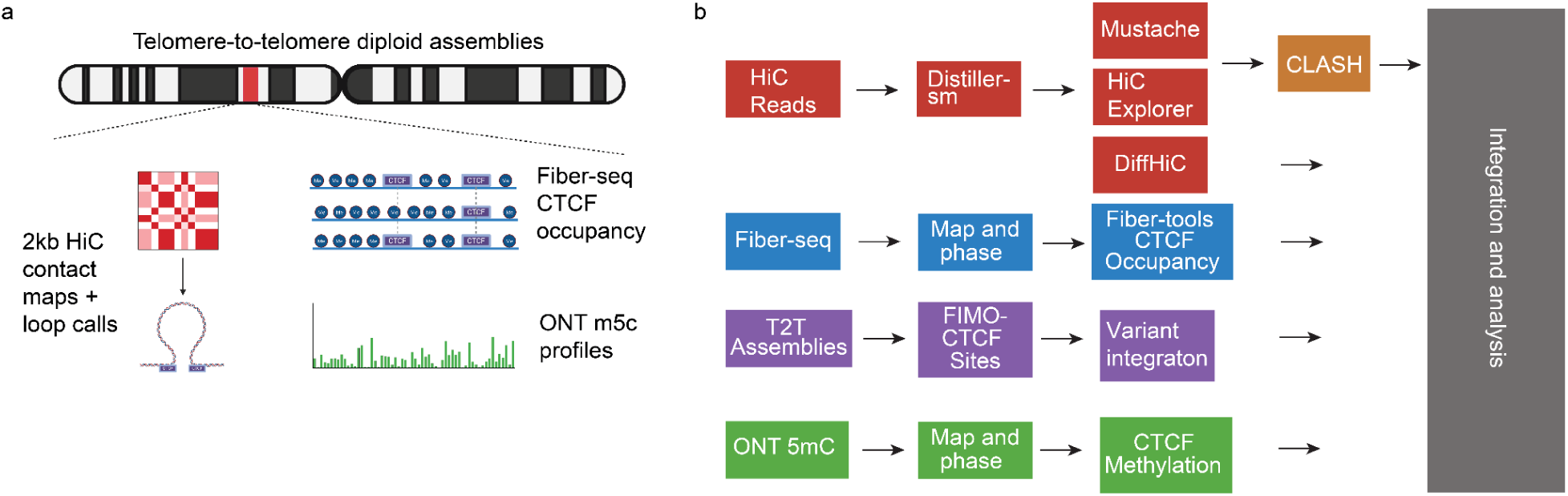
Multi-omic data generation and processing across five lymphoblastoid samples. **a,** Overview of all datasets integrated in this study; Hi-C contact maps, Fiber-seq m⁶A–based chromatin accessibility, and CTCF occupancy datasets for GM19317, GM19347, HG01457, HG02666, and HG03248 were generated, while ONT-derived m⁵C CpG methylation and haplotype-resolved variant calls for those samples were obtained from previously published datasets. All non-Hi-C data modalities are both phased and haplotype-aware. Hi-C contact matrices were aligned to the GRCh38 reference genome, and Fiber-seq, m⁵C methylation, and variant data were aligned or lifted over to GRCh38 to enable joint analysis. **b,** Schematic overview of the computational workflows used in this study, including Hi-C processing and loop calling (distiller-sm, diffHiC, Mustache, and HiCExplorer), loop scoring using the CLASH method introduced here, Fiber-seq alignment and CTCF occupancy calling (WhatsHap and fibertools), m⁵C CpG methylation phasing (WhatsHap), structural variant integration (pav2 pipeline), and CTCF motif identification (FIMO).

### Differential contacts are associated with both sequence and epigenetic variation

We first measured how genetic and epigenetic variation influence Hi-C maps irrespective of specific contact patterns or domain structures. Each sample was compared against a set of the remaining samples using DiffHiC (Lun and Smyth 2015) at 5 kb resolution to identify 367 differential Hi-C pixels (defined as the interaction-pair between two bins) after stringent filtering (Methods; Supplementary Figures 3-7). These formed 123 clusters, which we defined as groups of pixels within 10 kb of each other. We then assessed whether the samples with a differential Hi-C pixel matched the samples with the respective class of variation, comparing against a null match rate of 20% under independence. We considered four mechanistic sources of variation that could generate differential interaction-pairs: (i) DNA sequence changes between interaction-pair bins, (ii) DNA sequence changes within interaction-pair bins, (iii) changes in chromatin accessibility, and (iv) CpG methylation changes. Representative examples of differential interactions associated with each mechanism are shown in annotated Hi-C heatmaps (Figs. 2b-e). For the 240 of the 367 differential interactions where at least one source of variation was present, we observed a match rate of 63.3% between the sample with a differential interaction and the sample with a mechanistic source of variation (Fig. 2f).

**Figure 2.**
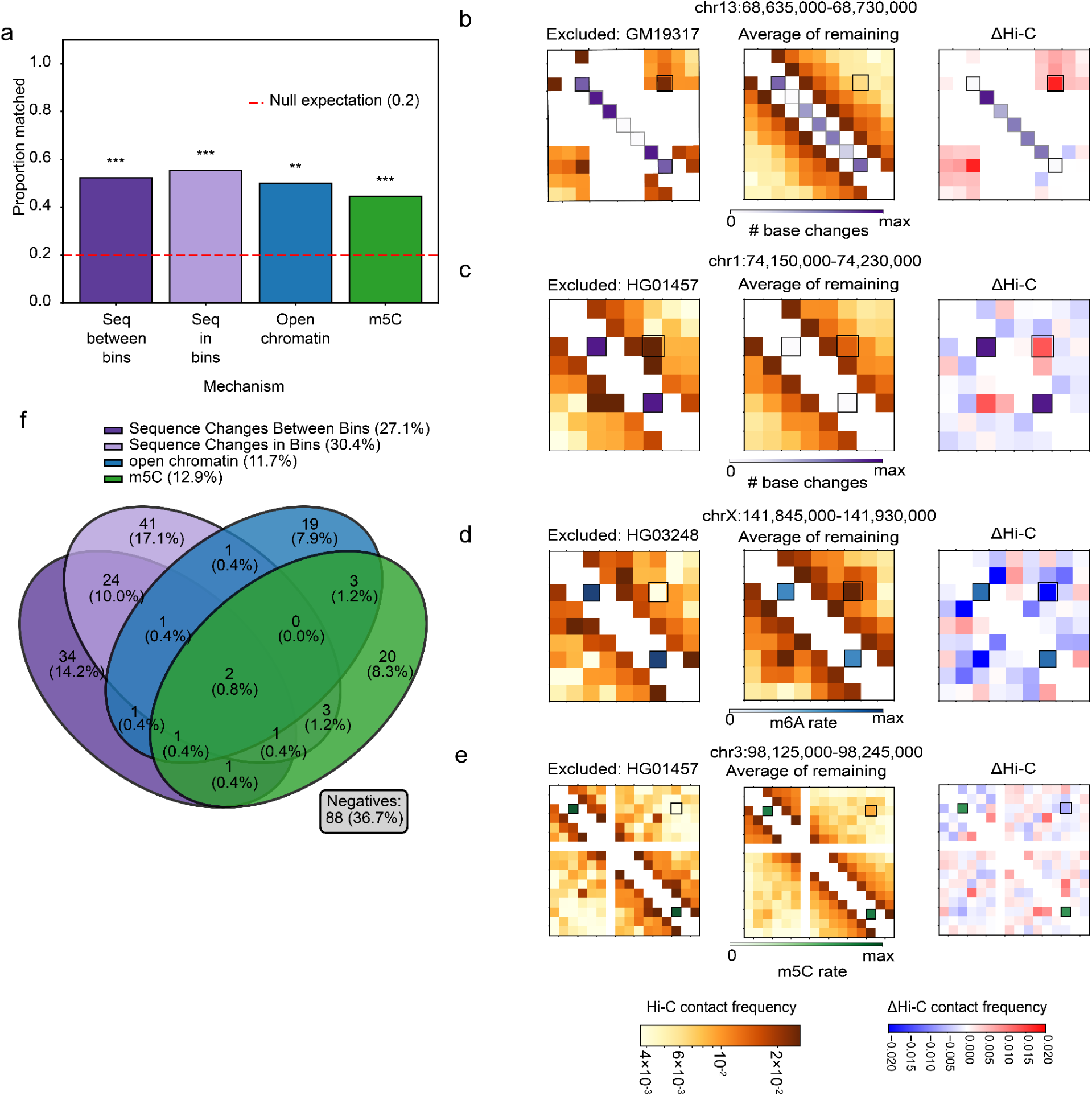
Mechanisms affecting diffHiC-identified differential chromatin interactions. **a,** Match rates for four mechanisms – sequence changes between interaction bins (dark purple), sequence changes within interaction bins (light purple), m⁶A methylation (blue), and m⁵C methylation (green) – after accounting for the influence of the other three mechanisms. All mechanisms remain significantly associated with differential chromatin contacts (between interaction bins: 52.3%, n = 65, p = 7.39 × 10^-9^; in interaction bins: 55.4%, n = 74, p = 1.97 × 10^-11^; m⁶A: 44.2%, n = 43, p = 3.47 × 10^-4^; m⁵C: 44.4%, n = 45, p = 2.20 × 10^-4^). Significance levels are denoted as p < 0.05 (*), p < 0.01 (**), and p < 0.001 (***), and were calculated using the binomial test compared against the null match rate of 0.2. **b,** Example of a differential chromatin interaction driven by sequence changes between interaction bins. Left: Hi-C contact map for the excluded sample (GM19317) at chr13:68,635,000–68,730,000. Middle: Average Hi-C contact map of the remaining four samples. Right: ΔHi-C map (GM19317 − mean of others), highlighting the interaction that differs most strongly in GM19317. The purple diagonal bins mark the total number of base changes between the two interacting bins of the differential interaction, and the purple rectangle centers around the focal contact whose strength deviates in GM19317. Compared to the other samples, GM19317 carries a greater number of sequence changes between interaction bins, corresponding to an increase in contact strength (red shift). **c,** Example of a differential chromatin interaction at chr1:74,150,000-74,320,000 driven by sequence changes in interaction bins. Shown in the same layout as panel b. Interaction bins are annotated in purple for the number of bases changed within the bins. The highlighted interaction shows increased contact strength in HG01457 (red shift). **d,** Example of a differential chromatin interaction at chrX:141,845,000-141,930,000 aligning with differential accessibility in interaction bin anchors. Shown in the same layout as panel b. Interaction bins are annotated in blue for the chromatin accessibility rate in each bin. The highlighted interaction shows decreased contact strength in HG03248 (blue shift). **e,** Example of a differential chromatin interaction at chr3:98,125,000-98,245,000 driven by m⁵C methylation rate in interaction bins. Shown in the same layout as panel b. Interaction bins are annotated in green for the m⁵C methylation rate in each bin. The highlighted interaction shows decreased contact strength in HG01457 (blue shift). **f,** Summary of the 240 differential bins exhibiting at least one differential mechanism. In total, 63.3% of these bins can be explained by one or more of the four mechanisms shown in panel a.

We next examined the match rate for each source of variation separately. When one sample had DNA sequence changes between interaction-pair bins, 60.7% matched that sample (n = 107, Supplementary Figure 8). When one sample had sequence changes within interaction-pair bins, 61.9% matched (n = 118; Supplementary Figure 9). Together, these results demonstrate that both large-scale structural rearrangements and smaller sequence changes in interaction-pair bins are strongly associated with altered chromatin contact strength across individuals, consistent with previous findings (Gorkin et al. 2019; Li et al. 2024).

As there were differential pixels without DNA sequence changes, we then measured the impact of epigenetic variation in the same samples. To reduce confounding between general chromatin accessibility and CTCF occupancy, we examined differential loci lacking a nearby CTCF site and found that decreased contacts were associated with increased chromatin accessibility (match rate = 51.4%, n = 35; Supplementary Figure 10). Additionally, differentially decreased contacts were associated with increased CpG methylation (match rate = 41.9%, n = 74; Supplementary Figure 11), consistent with previous findings (Monteagudo-Sánchez et al. 2024). The reciprocal relationship does not significantly differ from the null for either source of epigenetic variation, suggesting that while chromatin accessibility and CpG methylation may be linked to reduced chromatin interactions, their absence is not sufficient to promote them.

After accounting for each of the four mechanisms as covariates, all remained enriched among differential interactions (Fig. 2a-e). The adjusted match rates were 52.3% for bases changed between interaction-pair bins (n = 65, p = 7.39 × 10^-9^, binomial test), 55.4% for bases changed within interaction-pair bins (n = 74, p = 1.97 × 10^-11^, binomial test), 50.0% for chromatin accessibility (n =20, p = 2.59 × 10^-3^, binomial test), and 44.4% for m⁵C methylation (n = 45, p = 2.20 × 10^-4^, binomial test). When a sample had at least one deviant mechanism for a given pixel, there was a corresponding change in that sample’s pixel contact 63.3% of the time (Fig. 2f). Together, these results demonstrate that genetic variation, differences in chromatin accessibility, and CpG methylation collectively contribute to inter-individual variation in chromatin contact frequency.

Most bins with a differential interaction-pair had only one differential pixel, indicating isolated differences across genomes (Supplementary Figure 12). Among these differential pixels, 39% were contained in homozygous deletions of at least one anchor bin and 15% were within 10 kb of chromatin loops), both significantly higher than expected by chance (p < 0.001 for both, permutation test; Supplementary Figure 13). Similarly, we found that the majority of clusters of differential interaction-bins were either overlapping chromatin loops or deletions, and that differential interaction-bins not clustered were not associated with these features (Supplementary Figure 14).

### Multimodal data confirms the association of genetic and epigenetic alterations and CTCF binding

While deletions are directly interpretable for changes in contacts, the differential contact-bins at chromatin loops where there are no deletions may be caused by more complex variation. The influence of genetic and epigenetic variation on chromatin structure including loops is known to be in part mediated by CTCF binding (Rowley and Corces 2018). To measure these relationships using our integrated resource, we calculated sample-specific motif strength (PWM score) with occupancy from Fiber-seq across 262,463 CTCF sites identified by FIMO annotation of the 10 haplotype assemblies. We found a positive association between PWM score and occupancy (Pearson’s r = 0.38, p < 2.2 × 10^-308^; Fig. 3a), confirmed by a negative association of the number of bases differing from the consensus motif (Pearson’s r = –0.21, p < 2.2 × 10^-308^; Supplementary Figure 15). We next measured how epigenetic variation is associated with differential CTCF binding. Among the ∼33% of CTCF sites containing a CG dinucleotide, the CpG methylation within each motif was negatively correlated with occupancy (Pearson’s r = -0.41, n = 90,673, p < 2.2 × 10^-308^; Fig. 3b). This trend was also supported by comparing occupancy across hypo-, hyper-, and mixed-methylation states (Pearson’s r = -0.64, hypomethylated n = 58,094 sites, hypermethylated n = 140 sites, mixed n = 14 sites, p = 2.2 × 10^-37^; Supplementary Figure 16).

**Figure 3.**
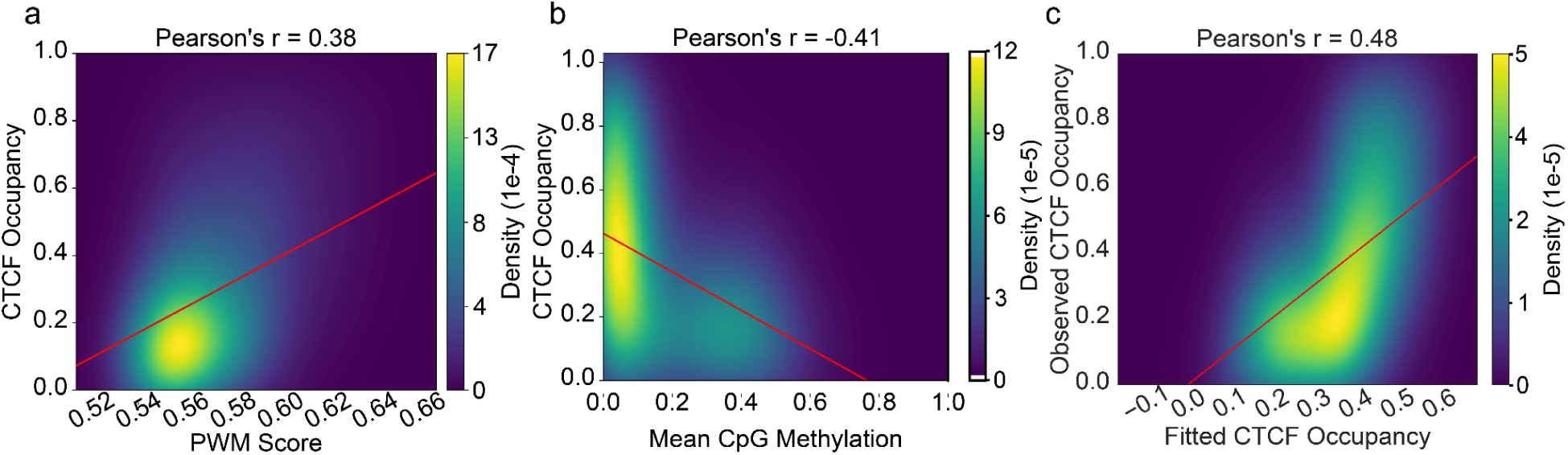
Quantification of genetic and epigenetic mechanisms influencing CTCF occupancy. **a,** Correlation between CTCF motif strength, through position weight matrix (PWM) scores, and Fiber-seq–derived CTCF occupancy (Pearson’s r = 0.38, n = 262,465 sites, p < 2.2 × 10^-308^). PWM scores were computed as the motif-averaged probability per nucleotide for each CTCF motif instance in each haplotype. Density denotes the local proportion of observations estimated by two-dimensional gaussian kernel density estimation evaluated on a 200 x 200 grid, which is the default mesh size used in all correlation visualizations unless otherwise noted. **b,** Correlation between m⁵C CpG methylation and CTCF occupancy (Pearson’s r = −0.41, n = 90,599 sites containing at least one CpG in any position in either haplotype, p < 2.2 × 10^-308^). For each CpG-containing motif methylation was calculated as the mean m⁵C methylation percentage across all CpG sites within that motif. **c,** Linear model estimating the combined effects of PWM score and m⁵C methylation values on CTCF occupancy using ordinary least squares regression (n = 90,673 sites). Both predictors showed significant associations with occupancy (PWM β = +0.406, m⁵C β = −0.085), and the model explained a moderate fraction of variance (Pearson’s r between observed and fitted values = 0.48, F test p = 1.1 × 10^-16^). Density estimation was evaluated on a 300 x 300 grid.

We next tested whether sequence and epigenetic variation jointly provide more predictive power for CTCF occupancy. We found that a regression model combining PWM score and m⁵C methylation values explained more variance in CTCF occupancy than either factor alone (Pearson’s r between observed and fitted values = 0.48, PWM β = +0.406, m⁵C β = –0.085, n = 90,673, F test p = 1.1 × 10^-16^; Fig. 3c). This is consistent with prior work showing that SNPs and methylation of CTCF binding sites can affect CTCF binding (Wang et al. 2012; Zeng et al. 2023).

We also considered whether local chromatin accessibility predicted differential CTCF binding. Chromatin accessibility was determined as mean m⁶A methylation rates across the flanking 2 kb of each 1 kb bin containing a CTCF site. We found that local accessibility only weakly correlated with occupancy (Pearson’s r = 0.11, n = 262,282, p < 2.2 × 10^-308^; Supplementary Figure 17). Consistently, adding accessibility to the regression model increased Pearson’s r by 0.02 (Supplementary Figure 18). Given this weak association, we excluded accessibility from downstream analyses.

### CLASH loop scores enable comparative analysis of CTCF occupancy, genetic variation, and epigenetic variation on chromatin loop strength

Because differential pixels were associated with loop structures, we sought to refine the association between loop strength and genetic/epigenetic variation. Loops were identified on all five of our samples using Mustache (Roayaei Ardakany et al. 2020) and HiCExplorer (Wolff et al. 2022) at 2 kb resolution. Consistent with prior analyses of GM12878 (Roayaei Ardakany et al. 2020), 29.3% of pooled loop calls were loci anchored by two unique CTCF sites in at least one haplotype (Supplementary Figure 19). Consistency across callers was low (Fig. 4a, Supplementary Figures 20 and 21), as reported in prior within-sample benchmarking (Chowdhury et al. 2024; Forcato et al. 2017). We additionally observed inconsistent loop detection across individuals despite similar contact profiles (Figs 4a,b, Supplementary Figure 22). This indicates that binary loop-calling alone does not robustly capture biological variation or reliably represent presence across samples.

**Figure 4.**
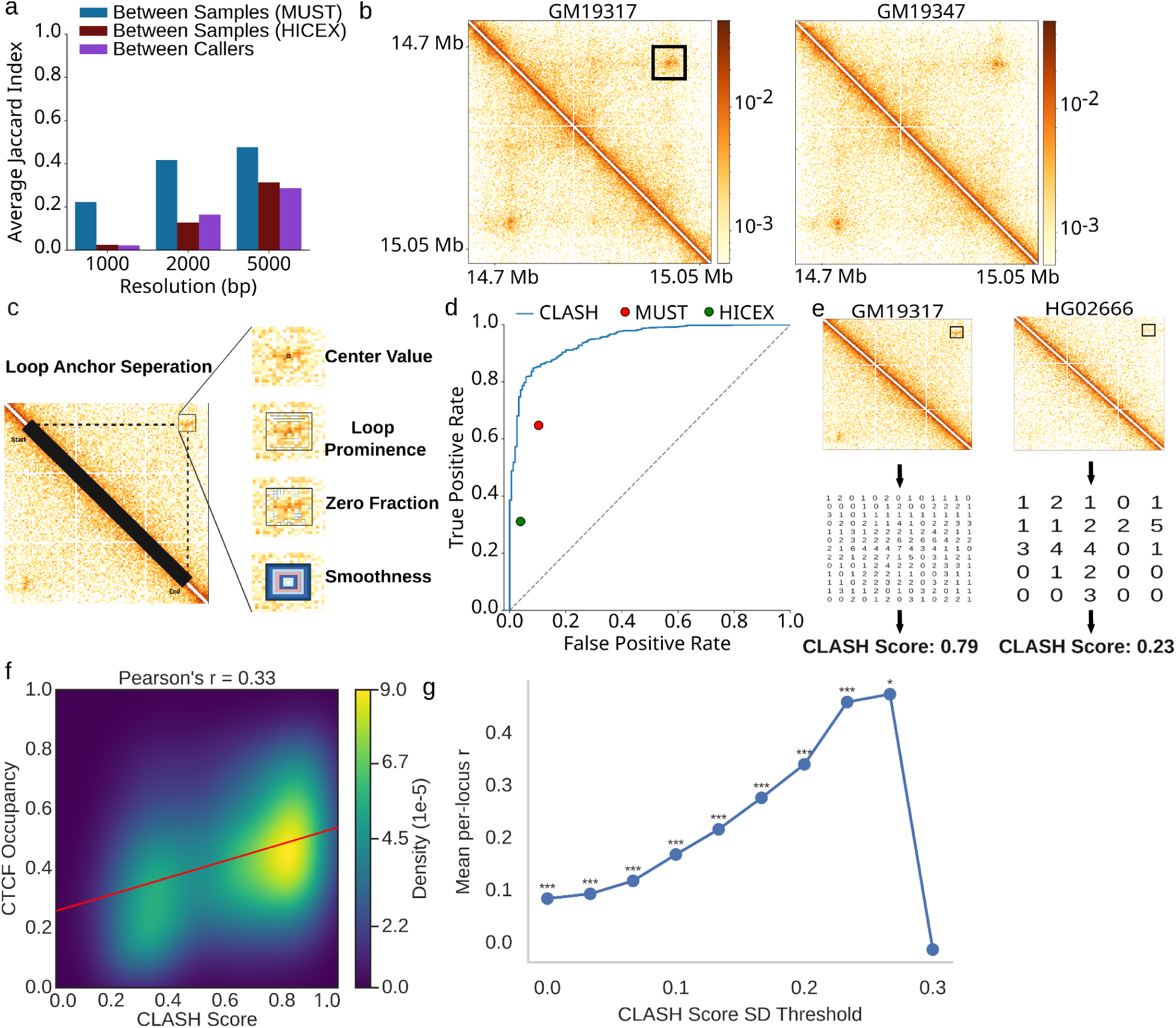
CLASH unification of loop calls and relation to CTCF occupancy. **a**, Average pairwise Jaccard index of loop calls between samples with a tolerance of 10 kb, shown separately for Mustache (MUST; blue) and HiCExplorer (HICEX; red), across 1 kb, 2 kb, and 5 kb resolutions. The average Jaccard index of within-sample loop calls between Mustache and HiCExplorer is also shown (purple). Values are displayed for loops filtered at a significance threshold of p = 0.1. **b**, Example chromatin loop for samples GM19317 (left) and GM19347 (right) from the locus chr5:14728000–15000000, illustrating loop-calling inconsistency across samples at the same locus. Both Mustache and HiCExplorer fail to identify the same loop across individuals; they call the loop only in sample GM19317 (black rectangle) despite clear contact enrichment in GM19347 (and the other three samples; see Supplementary Figure 28 for all five samples). **c,** Schematic illustration of the five features – loop anchor separation, center value, loop prominence, zero fraction, and smoothness – CLASH utilizes in addition to raw LC scores to score loops for the representative region of chr18:53883045-54145059 for GM19317. **d**, CLASH AUROC (blue, AUROC = 0.945) loop classification performance using out-of-fold predictions with a five-fold GroupKFold cross-validation, compared to the fixed (FDR, TPR) points of initial Mustache (MUST; red) and HiCExplorer (HICEX; green) loop calls on our curated training set. **e**, Schematic illustration of how CLASH scores loops at the same locus (representative region of chr18:53883045-54145059 for samples GM19317 and HG02666) by extracting the matrix around the local maximum count. Displayed counts were multiplied by 10^3^ to improve readability. **f,** Global correlation between CTCF occupancy and loop strength across all samples (Pearson’s r = 0.33, n = 21,050, p < 2.2 × 10^-308^). **g,** Distribution of Pearson correlation values between CTF occupancy and loop strength across loci with increasingly stringent CLASH score standard deviation thresholds (n = 3,144). Significance was assessed using a one-sided one-sample t-test against zero. Significance levels are denoted as p < 0.05 (*), p < 0.01 (**), and p < 0.001 (***).

Overcoming these limitations requires aggregating loop calls across samples and quantifying loop strength directly from the underlying Hi-C matrices as a continuous score. We initially considered the “LC scores” computed by the Mustache method which quantify the local contrast between a candidate loop and its background at multiple scales via a difference of Gaussians. These continuous scores were not previously used as a measure of loop-strength, but instead relegated as an intermediary internal component Mustache used to generate its final binary callset. To test if LC scores can be used to distinguish loops from non loops, we took an approach motivated by machine learning (Salameh et al. 2020) to first manually curate a truth of 1000 loops across 200 loci with loop presence or absence. We calculated LC scores following Mustache’s implementation, but assigned the maximum calculated LC score across scales to each putative loop. We calculated the AUROC using both these raw continuous scores (AUROC = 0.904) as well as using a logistic regression model trained on the raw LC scores with 5-fold cross validation, evaluating out-of-fold predictions (AUROC = 0.904; Supplementary Figure 23).

We also considered how other features and models performed on the same manually labelled dataset, using the same 5-fold cross validation framework. We first engineered five separate features derived from the Hi-C matrices at 2 kb resolution – loop anchor separation, center value, loop prominence, zero fraction, and smoothness (Fig. 4c). A logistic regression model trained on these five features (“Multifeature Logistic Regression”) performed similarly to LC with an AUROC of 0.894, while an XGBoost model trained on these features (“Multifeature XGBoost”) outperformed LC with an AUROC of 0.921 (Supplementary Figure 23). Including the raw LC scores as a sixth feature in the XGBoost model increased the out-of-fold AUROC to 0.945, indicating that LC scores and the other five features have complementary effects on loop classification performance (Supplementary Figures 23, 24, 25). We name this XGBoost model incorporating the six features Chromatin Loop Across-sample Score Harmonizer (CLASH). To enable further fair comparisons between CLASH and LC scores, we chose to use the logistic regression LC model (hereafter called “LC model”) for downstream analyses as opposed to the raw scores, but we note that we observed similar results when using the raw scores.

While both LC and CLASH outperform initial loop calls by Mustache and HiCExplorer (Fig. 4d), CLASH achieves a higher recall than LC (0.787 vs 0.715) when matching the false positive rate at 0.052 (Supplementary Figure 26). For both CLASH and LC model, we defined a threshold to separate loops from non-loops by selecting the threshold that maximized Youden’s J on the full training dataset (using out-of-fold predictions), which we used for all downstream analyses requiring binary calls. Under this optimized threshold, the harmonization of out-of-loop across individuals calls by CLASH improved classifications in a net 71 of 199 loci, while LC model harmonization improved a net 33 loci, demonstrating CLASH’s ability to refine loop labels at 5 kb resolution even when initial loop detection is underpowered (Wolff et al. 2022) (Supplementary Figures 27, 28). Using CLASH to harmonize loop calls across our full dataset revealed that 54% of loops that are present in at least one sample are shared in two or more samples (Supplementary Figure 29).

We extracted the min-max scaled logit of each CLASH classification to serve as a continuous loop-strength score, enabling comparison of loop strength across samples, and followed the same procedure for LC to enable score comparisons (hereafter called “LC model scores”). Although out-of-fold CLASH scores and LC model scores from our training set were highly correlated (r = 0.80), we observed that for the 141 CLASH (out-of-fold) misclassifications and 197 LC model (out-of-fold) misclassifications, CLASH scores were consistently closer to the decision boundary than LC model scores in each method’s score space (CLASH mean error = 0.21, LC model mean error = 0.31, Mann–Whitney U test p = 8.2 × 10⁻^5^; Supplementary Figure 30). This indicates that CLASH produces better-aligned scores with respect to the classification boundary, with misclassifications tending to occur near the decision threshold rather than far into the incorrect region, thereby supporting the use of CLASH scores for downstream quantitative interpretation. Importantly, CLASH scores also have minimal sample-specific bias when calculated on the full dataset (maximum absolute standardized deviation from the global mean = 0.12; Supplementary Figure 31), enabling locus-specific cross-sample comparisons (Fig. 4e).

As CTCF binding occupancy mediates boundaries of loops (Fudenberg et al. 2016; Rao et al. 2015), we considered the CTCF occupancy measured by Fiber-seq as an orthogonal support for CLASH scores. For the 6,377 CTCF-flanked loops, the CLASH score was positively associated with occupancy (Pearson’s r = 0.33, n = 21,050, p < 2.2 × 10^-308^; Fig. 4f). To assess locus-specific effects, we examined loci with increasing inter-sample variability in CLASH score (n = 3,140; Fig. 4g, Supplementary Figure 32). Higher occupancy was positively correlated with increased loop score, consistent with prior work showing that binary changes in CTCF binding coincide with corresponding gains or losses of chromatin loops (Pugacheva et al. 2020); our results extend this to a continuous model of CTCF occupancy. Repeating these correlation analyses using LC model scores yields similar results (Supplementary Figure 33).

Using CLASH scores as a metric of loop strength, we next sought to measure how genetic and epigenetic variation jointly affect loop strength. Using loops formed between two CTCF sites, we quantified the association between sequence or methylation variation and loop strength, averaging PWM scores and m⁵C methylation values across both anchor sites. PWM scores showed a positive correlation with loop strength (Pearson’s r = 0.242, n = 31,822, p < 2.2 × 10^-308^; Fig. 5a, Supplementary Figure 34) as expected (Gorkin et al. 2019; Monteagudo-Sánchez et al. 2024). Stratifying m⁵C values into groups shows that loop strength decreases with increasing methylation (n = 19,440, all pairwise Mann-Whitney U test p values ≤ 6.3 × 10^-38^; Supplementary Figure 35), with the lowest methylated group (≤ 0.1, n = 14,655) showing the highest median loop strength (0.709), while the highest methylated group (> 0.3, n = 1,230) has the lowest median loop strength (0.407).

**Figure 5.**
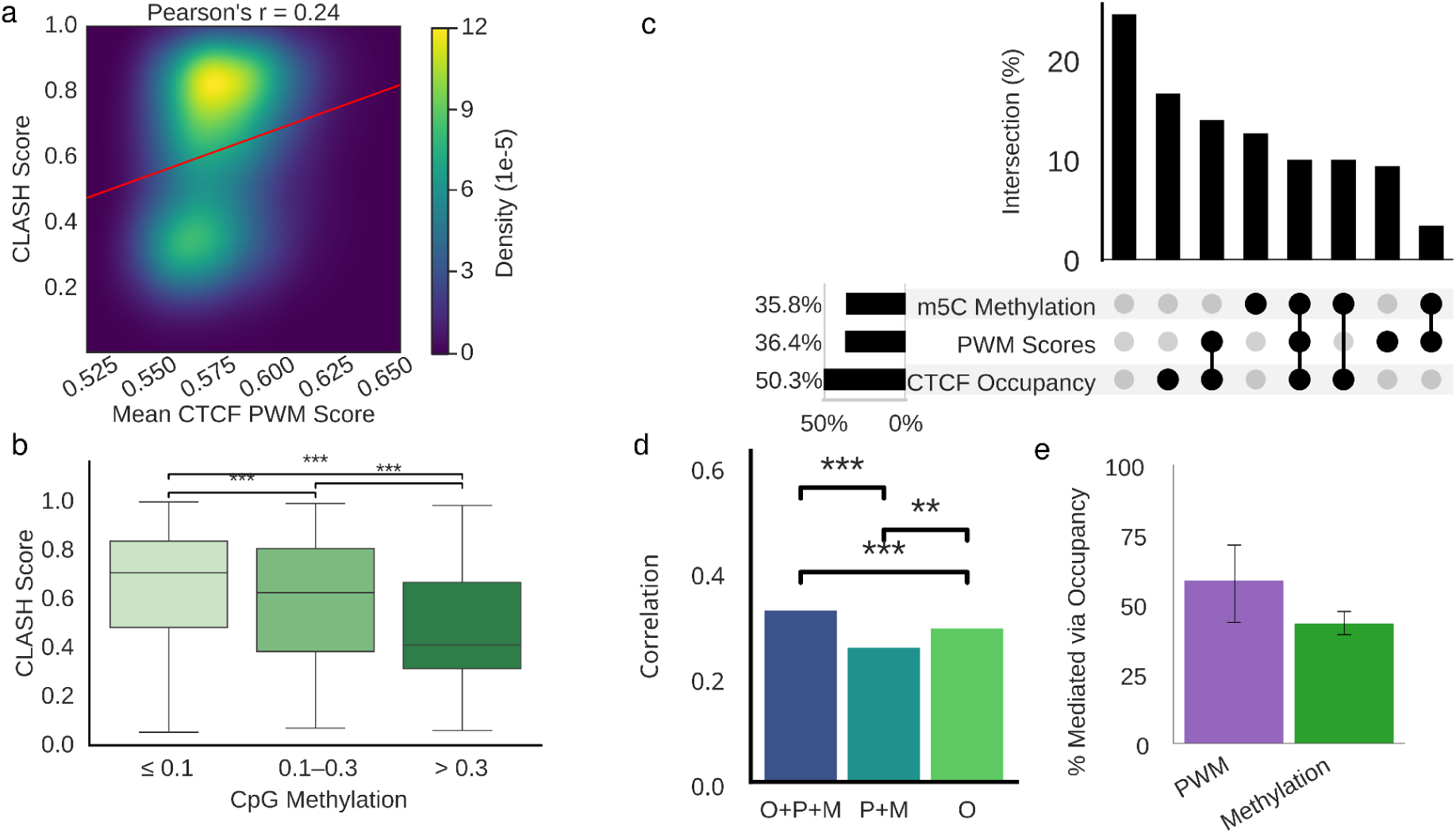
PWM score and CPG methylation correlations with loop scores and mediation analysis. **a,** Correlation between PWM scores (averaged across both haplotypes and both loop-associated CTCF sites) and loop strength (Pearson’s r = 0.242, n = 31,822, p < 2.2 × 10^-308^). **b,** Distribution of CLASH loop scores stratified by mean CpG methylation. CpG methylation is stratified into ≤ 0.1 (median loop score = 0.709, n = 14,655), 0.1 - 0.3 (median loop score = 0.632, n = 3,568), and > 0.3 (median loop score = 0.407, n = 1,230). Mann Whitney U tests yielded p values of 6.28 x 10^-38^ between the ≤ 0.1 and 0.1 - 0.3 groups, 4.85 x 10^-58^ between the 0.1 - 0.3 and > 0.3 groups, and 7.04 x 10^-146^ between the ≤ 0.1 and > 0.3 groups. **c,** Proportion of differential loops (n =151 loci with CLASH score range ≥ 0.3 and occupancy, PWM scores, and m⁵C methylation data for at least 3 of the 5 samples) explained by each mechanism using r ≥ 0.25 as the explanatory threshold per-locus (CTCF occupancy = 50.3%; m⁵C methylation = 35.8%; PWM score = 36.4%; together explaining 75.5% of differential loops). **d,** Comparison of correlation of loop strength with combined occupancy (O), genetic (P), and epigenetic (M) features (left) versus correlation of loop strength with combined genetic and epigenetic features (center) versus correlating loop strength with occupancy alone (right). Including all three mechanisms together leads to the highest correlation, which is 0.0663 more than just including genetic and epigenetic features (n = 13,094, Steiger p < 2.2 × 10^-308^) and 0.0408 more than just including occupancy as a feature (n = 13,094, Steiger p < 2.2 × 10^-308^). **e,** Product-of-coefficient mediation analysis quantifying the proportion of PWM scores (57.1%, n = 3,921) and m⁵C values (40.2%, n = 13,094) effects on loop strength that act through CTCF occupancy, averaged across samples. Black bars denote 95% bootstrap confidence intervals on the mediated proportion, estimated by pooled bootstrap of the indirect effect.

We next quantified the contribution of each mechanism with differential loops. Among the 151 loci where loop strength varied by more than 0.3 (chosen from Supplementary Figure 36) and where occupancy, PWM score, and methylation data were available for at least three of the five samples, 50.3% showed a correlation between occupancy and loop strength above r ≥ 0.25, while 35.8% were associated with m⁵C methylation and 36.4% with PWM score. Together, these mechanisms accounted for 75.5% of differential loops (Fig. 5c).

To determine if occupancy accounts for the observed mechanistic effect on loop strength, we calculated the joint association of PWM score, m⁵C methylation, and occupancy with loop strength, and compared this to models excluding each factor. Including PWM score and m⁵C methylation alongside occupancy yielded a higher correlation with loop strength than occupancy alone (Δ Pearson’s r = +0.0408 n = 13,094, Steiger p < 2.2 × 10^-308^), and excluding occupancy from the full model resulted in a significantly lower correlation (Δ Pearson’s r = -0.0663, n = 13,094, Steiger p < 2.2 × 10^-308^; Fig. 5d). To quantify how much of the PWM score and m⁵C methylation effects on loop strength are mediated by occupancy, we applied a global product-of-coefficients mediation analysis averaged across samples, finding that 57% of the PWM score effect (n = 3,921 observations from 1,174 variant CTCF loci) and 40% of m⁵C values (n = 13,094 observations) act through CTCF occupancy (Fig. 5e). In comparison, performing the PWM mediation analysis across all 21,050 observations genome-wide, which introduces stronger between-locus effects to which mediation estimates are particularly sensitive, reduced the mediated percentage to 45%. Although a within-locus mediation analysis would isolate within-locus effects more directly, per-locus mediation estimates were unstable with only five samples per locus.

Because the occupancy-based measurements did not incorporate SVs that insert or delete CTCF sites, we separately evaluated whether nonreference CTCF binding sites may influence genome structure through differences in loop strength. Using the T2T assemblies to provide a complete view of sequence variation (Logsdon et al. 2025), we determined that each haplotype contained 107 ± 12 CTCF-containing insertions and 61 ± 7 CTCF-containing deletions. For each locus where CTCF-affecting SVs overlapped a loop anchor, we compared average loop scores between samples carrying the variant and those without. Although it has been shown that the insertion of ectopic CTCF binding sites can induce additional domain boundaries (Zhang et al. 2020), we identified only 27 loop loci overlapping insertion CTCF variants and 23 loop loci overlapping deletion CTCF variants among our ten haplotypes, limiting our ability to extend this analysis to naturally occurring CTCF variants at the genome-wide level (Supplementary Figure 37). However, we did observe several individual loci that exhibited clear loop-strength changes associated with variant CTCF sites at relaxed CTCF detection thresholds (Supplementary Figures 38 and 39), indicating that nonreference CTCF binding sites can affect genome structure.

## Discussion

The comparison of three dimensional structure of genomes across individuals can reveal mechanistic insight into how non-coding genetic variation leads to phenotypic differences. To address this, we generated a multi-modal dataset of high-resolution contact maps with single-molecule chromatin accessibility and CTCF binding data complemented by T2T assemblies and haplotype-resolved methylation calls across five diverse samples. However, existing loop calling approaches demonstrated inconsistent calls between individuals. Thus, we developed CLASH, a method that generates continuous scores directly from high-resolution Hi-C contact maps to harmonize loop strength across samples.

The CLASH scores enabled comparative analysis of relative contributions of different forms of variation to loop formation. We found that the inclusion of epigenetic (m^5^C) and CTCF occupancy measurements provides a more complete understanding of differential loop formation than sequence variation alone. Crucially, the contribution of methylation to differential loop formation is nearly equivalent to that of sequence variation, while differential CTCF binding contributes substantially more (1.4x). Furthermore, in the assayed cell lines, the co-occurrence of differential methylation and sequence variation is minimal (3.3% of all differential sites). Although some of the observed epigenetic changes may be due to random silencing as a cell-line artifact (Mikkelsen et al. 2007), this provides advantageous variability for our analysis of changes that affect chromatin structure. Our measurements are consistent with the importance of CTCF binding to predict genome structure. However, because the inclusion of methylation and sequence variation improves association with loop strength, they reflect latent information that improves inference beyond open chromatin and CTCF binding, highlighting the importance of unified measurements of genetic and epigenetic sequence variation in future studies.

This study is primarily limited by the sample size; future studies with larger sample sizes will be necessary to find the per-locus mediation effects of PWM scores and methylation on loop strength through occupancy, significant relationships between naturally occurring CTCF insertions and deletions with loop formation/strength genome-wide, and the enrichment of expression quantitative trait loci with loops of increasing variability (Supplementary Figure 40). An increased sample size will also be valuable for training more advanced machine learning approaches to harmonize loop calling across individuals and improve model generalizability across datasets. For example, because CLASH features such as zero fraction and center value depend on sequencing depth and matrix sparsity, CLASH may require recalibration when applied across substantially different sequencing depths or resolutions. Accordingly, the current implementation is primarily intended to analyze high-coverage Hi-C datasets at 2 kb resolution. Finally, the low number of differential pixels identified by DiffHiC indicates that the development of new methods with more power to identify differential features could be helpful in comparing Hi-C maps between samples.

As the number of population and longitudinal studies such as the 4D nucleome project (Dekker et al. 2026) on genome structure grow, methods to quantify and harmonize features across conformation capture datasets will be increasingly important for relating 3D genome organization to genetic variation. The growing interest in feature quantification and harmonization is highlighted by emerging methods that harmonize loop positions across separate samples by pooling information (Liu et al. 2026) and quantify loop strengths by integration over focal regions of enrichment (Jusuf et al. 2025). Furthermore, as there is a sharp interest in predicting molecular phenotypes using machine learning (Tang 2025; Avsec et al. 2026), our data provides additional support that personalized epigenomes at the resolution of individual bases should be considered alongside personalized genomes to predict cell state. For example, we used AlphaGenome (Avsec et al. 2026) to predict the contact maps for each haplotype from sequence alone, centered around the same example loci that we used to demonstrate the effects of CTCF occupancy, genetic variation, and methylation on loop formation. As expected, while AlphaGenome accurately predicted the variation in observed loops due to sequence variation at CTCF sites (Supplementary Figure 41), variation in observed loops caused by differential CTCF occupancy due to methylation or other stochastic variation is not predicted (Supplementary Figures 42 and 43), further indicating that sequence-to-map models may benefit from incorporating these signals (Dubocanin et al. 2025).

## Data and code availability

Hi-C paired-end sequencing reads and Fiber-seq data are available from the 1000 Genomes ftp: https://ftp.1000genomes.ebi.ac.uk/vol1/ftp/data_collections/HGSVC3/working/.

Software is available at github.com/chaissonlab/clash. Tables of clash calls on all five genomes, features used for calling loops, and analysis scripts are available at https://zenodo.org/records/20412945.

## Acknowledgements

This work was supported by NHGRI grants U24HG007497 and HG011649. M.J.P.C. is an employee of Pacific Biosciences.

## Methods

### Data processing

All data used corresponded to five lymphoblastoid cell lines, GM19317, GM19347, HG01457, HG02666, and HG03248 (Logsdon et al. 2025). Hi-C data for all samples were generated, aligned to GRCh38, and processed using the distiller-sm pipeline (https://github.com/open2c/distiller-sm) to resolutions as high as 1 kb. Phased SNP and structural variant calls for all samples were obtained from the HGSVC Phase 3 dataset (Ebert et al. 2021) generated from telomere-to-telomere haplotype assemblies aligned to GRCh38, and processed with dipcall (Ebert et al. 2021; Li et al. 2018) and WhatsHap (Martin et al. 2023) to annotate CTCF-overlapping insertions, deletions, and single nucleotide polymorphisms (SNPs). Single-molecule Fiber-seq reads, generated with PacBio HiFi for all samples, were aligned to GRCh38, phased with WhatsHap, and processed with pbmm2 (Li 2018), whatshap, and fibertools (Jha et al. 2024) to determine CTCF occupancy, with occupancy defined as the fraction of fibers showing a footprint at each motif. Haplotype-specific CpG methylation tracks were downloaded from HGSVC via globus, aligned to GRCh38 with minimap2, phased with WhatsHap, and summarized per genomic bin and per CTCF site.

### A/B compartment analysis

A/B compartments were computed from ICE-balanced Hi-C matrices at 100 kb resolution using Cooltools eigs_cis, with GC content used to orient the first eigenvector such that positive values correspond to active (A) chromatin. Compartment similarity across individuals was assessed using pairwise eigenvector mean squared error (MSE) and sign concordance.

### Large-scale accessibility track analysis

For all samples, Fiber-seq reads across both haplotypes were partitioned into 100 kb genomic bins, the proportion of methylated adenines/total adenines within each bin was computed, and the resulting profiles were mean-centered. For each pair of samples, we computed the average MSE between accessibility profiles and sign concordance as the percentage of bins in which both samples showed deviations from the mean accessibility in the same direction (positive or negative relative to their respective means).

### DiffHic analysis

Genome-wide differential chromatin interactions were identified using diffHic (Lun and Smyth 2015) by following the protocol detailed in the “diffHic User’s Guide”. We generated differential interaction lists from raw Hi-C contact matrices at 1 kb, 2 kb, and 5 kb resolutions by filtering out non-relevant pixels, normalizing for bias with normOffsets(), estimating dispersion, and applying quasi-likelihood tests across all samples. We retained differential pixels that exhibited sufficient coverage (logCPM > –4) and that passed Bonferroni multiple-test correction at genomic distances between 10 kb and 1 Mb. Each differential interaction was then annotated for four potential mechanisms – large structural variants, small sequence changes, m⁶A methylation, and m⁵C CpG methylation – using outlier frameworks to identify the sample or haplotype exhibiting the most extreme deviation for each feature. For each mechanism, we quantified whether the sample with the strongest molecular deviation corresponded to the sample with the largest Hi-C log-fold change and assessed significance using binomial tests. Loci explained by multiple mechanisms were then removed to confirm that each mechanism remained significantly enriched.

### CTCF site determination and PWM score calculation

Because 85% of chromatin loops form at CCCTC-binding factor (CTCF; (Rao et al. 2014), CTCF sites were identified in each haplotype by running FIMO (Grant et al. 2011) on haplotype-resolved assemblies and mapping the resulting coordinates back to GRCh38 using the Long Read Aligner software (Grant et al. 2011; Ren and Chaisson 2021), yielding ∼50,000 sites per haplotype. This matches expected CTCF site counts per haplotype (The ENCODE Project Consortium 2012).

For each motif, the underlying 19 bp sequence was scored using a position weight matrix (PWM) derived from the JASPAR MA0139.1 probability-odds matrix (Khan et al. 2017), producing a continuous measure of motif strength that reflects the impact of SNPs and small indels. PWM scores were computed by summing position-specific probability weights and normalizing by motif length to facilitate comparison across haplotypes and individuals. Sequence mismatches relative to the canonical 15 bp CTCF consensus motif were enumerated by aligning each haplotype sequence to the consensus and counting non-matching bases.

### CTCF m⁵C CpG methylation

CpG methylation levels were quantified at single-base resolution from phased ONT methylation calls produced by the HGSVC, with per-position methylation fractions computed as the proportion of reads supporting a methylated cytosine. For each haplotype-resolved CTCF motif, all CpG positions within the motif boundaries were extracted and averaged to obtain a site-level m⁵C methylation value. Additionally, a genome-wide hidden Markov model was used to classify each CpG as hypomethylated or hypermethylated. Similar results could be achieved from the Fiber-seq m^5^C reads, however the ONT reads were used as orthogonal support.

### CTCF site accessibility

m⁶A rates (the number of m⁶A adenines / the total number of adenines) were calculated within each 1 kb bin for each sample, as a proxy for chromatin accessibility. The average m⁶A rate of the four bins (two upstream, two downstream) surrounding each bin that contained a CTCF site was calculated. To account for technical biases from sequencing, the m⁶A rates were Z-scored within sample and sigmoid-transformed into a score between zero and one, representing how accessible the chromatin near each CTCF site was compared to the other CTCF sites in the sample. Z-scores were calculated within CTCF sites rather than genome-wide as downstream analysis was focused only on CTCF sites.

### Quantifying correlations between genetic and epigenetic factors and occupancy

Each CTCF site across all ten haplotypes was annotated with its PWM score, number of sequence mismatches, CpG methylation level and state, accessibility, and CTCF occupancy value. Global pairwise Pearson correlation coefficients were computed to assess how each genetic or epigenetic feature individually relates to CTCF occupancy. To evaluate their combined contributions, an ordinary least squares model was constructed using PWM scores, accessibility, and CpG methylation as joint predictors of occupancy after appropriate normalization. Because the goal was to quantify association rather than perform prediction, the model was fit to the full dataset and significance was evaluated using standard linear regression statistics.

### Calling loops with Mustache and HiCExplorer

Chromatin loops were initially identified using both Mustache (Roayaei Ardakany et al. 2020) and HiCExplorer (Wolff et al. 2022) across 1 kb, 2 kb, and 5 kb resolutions, filtering for p-values of 0.1 and 0.01. To assess agreement between callers and across individuals, we compared loop sets using Jaccard indices after standardizing loop coordinates within 10 kb. We used the 2 kb set of loop calls with the p = 0.1 threshold for downstream loop analysis.

### CLASH

We developed CLASH, a classifier XGBoost model that re-evaluates pooled loop calls across samples. For each sample, a refined loop center is determined and a dynamic matrix around each loop center is extracted based on the local matrix structure. The model was then trained on six features – “LC scores”, loop anchor separation, center value, loop prominence, zero fraction, and smoothness – to classify each sample at each locus as a loop or not, and the min-max scaled logits of each classification were used as the CLASH score. These continuous scores can replace inconsistent binary loop calls and enable robust cross-sample comparison of loop strength. Full model details are provided in the Supplementary Methods.

### CLASH validation

We compared CLASH’s AUROC classification performance to other model and feature combinations, and the classification performance of raw LC scores, Mustache initial calls, and HiCExplorer initial calls. To enable fair comparisons, we tested CLASH’s and LC model’s recall ability at a matched false positive rate. To demonstrate CLASH’s harmonization capability, we evaluated how many training loci initially called by Mustache and HiCExplorer were worsened, improved, or perfectly improved by CLASH loop predictions (evaluated out-of-fold) and compared this to the performance of the LC model. To validate the usage of CLASH scores, we computed the error severity of out-of-fold misclassifications on the training set and compared the results to those of the LC model. We also externally validated CLASH loop-scoring performance by computing Pearson correlation efficients between CLASH scores with CTCF occupancy globally and within-locus across samples.

### Calculating loop conservation

Loop presence was defined as loops exhibiting a CLASH score of > 0.59. For each locus whose two interaction bins each contained a CTCF site within 10 kb, we identified the number of individuals (max N = 5) in which a loop was present (k). The corresponding k-of-N conservation curve was generated and smoothed using cubic interpolation. The area under the curve (AUC) of the portion of the graph with k ≥ 2 corresponds to the conditional conservation probability.

### Quantifying the correlation between CTCF occupancy and CLASH loop score

To assess how CTCF protein binding relates to chromatin loop strength, we restricted the analysis to loops whose two interaction bins each contained a CTCF site within 10 kb and computed a per-loop occupancy value by averaging the Fiber-seq–derived occupancy of its two interacting CTCF sites. If multiple CTCF sites were located within 10 kb of a loop anchor, we chose CTCF sites by sequentially optimizing for CTCF orientation (such that anchors would have convergent orientations), motif strength (PWM scores), and minimal distance from the anchor bin. These occupancy values were paired with CLASH loop scores across all samples to quantify the global relationship between CTCF binding and loop intensity.

To evaluate locus-specific effects, we also quantified the distribution of Pearson’s correlation coefficients between CTCF occupancy and CLASH loop strength between samples at each locus. Loci with fewer than four samples containing valid occupancy measurements were excluded. A binomial sign test was used to calculate statistical significance.

### Quantifying the correlations between PWM scores, m⁵C methylation, CTCF occupancy, and CLASH loop scores

Per-loop PWM scores and CpG methylation values were obtained by averaging interaction bin measurements and pairing them with CLASH loop scores and occupancy. We quantified individual and joint associations using Pearson correlation coefficients, Mann-Whitney U tests, and Steiger’s test for dependent correlations, enabling direct evaluation of whether PWM scores and m⁵C methylation values contributed explanatory power beyond occupancy alone, including in a model excluding occupancy.

### Mediation analysis

To quantify how much of the effect of PWM scores and CpG methylation on loop strength is transmitted through CTCF occupancy, we performed a global product-of-coefficients mediation analysis. For PWM scores, we only pooled observations from variant CTCF loci. Uncertainty was measured using a pooled nonparametric bootstrap.

### Structural variation analysis

To assess whether structural variants (SVs) that add or remove CTCF motifs influence chromatin loop formation, we identified insertions and deletions overlapping loop-associated CTCF sites and quantified the resulting gain or loss of motif instances. For each loop and individual, we compared CLASH loop scores between samples carrying a CTCF-altering SV and those without, and evaluated the direction and magnitude of loop strength changes across the genome.

### Alpha Genome

To evaluate whether sequence variation alone is sufficient for current sequence-to-map models to accurately predict observed loops and assess whether they could benefit from incorporating methylation and CTCF occupancy data in addition to sequence data, we used Alpha Genome (Avsec et al. 2026)

## Supplemental Methods

### Data processing

Hi-C data were generated for five male lymphoblastoid cell lines (GM19317, GM19347, HG01457, HG02666, and HG03248), each sequenced in three independent runs by Phase Genomics (Seattle, WA), with one additional run being included from previous Human Genome Structural Variation Consortium (HGSVC) data (Logsdon et al. 2025). Raw FASTQ files were aligned to the GRCh38 reference genome and processed using the distiller-sm pipeline, excluding unplaced contigs. PCR duplicates, self-circles, and dangling ends were removed following standard Hi-C quality-control steps.

The lower-limit of contact resolution imposed by molecular byproducts for all samples was 1-2 kb, as summarized using MultiQC (v1.20; (Ewels et al. 2016)). To determine if the sequencing depth of our samples supported analysis at this high resolution, we adapted the Juicer (Durand et al. 2016) script provided by https://github.com/aidenlab/juicer/blob/main/misc/calculate_map_resolution.sh to guide plausible Hi-C resolution analysis for our samples, which yielded map resolutions ranging from 2.45 kb – 2.95 kb (Table S1). These values are reasonable given prior work that has shown that a read depth of ∼5 billion read pairs is necessary for Hi-C analysis at 1 kb resolution (Rao et al. 2015), ∼1 billion read pairs allows for slightly underpowered Hi-C analysis at 2 kb (Rao et al. 2015; Lee et al. 2022), and ∼500 million read pairs is sufficient for 5 kb analysis (Rao et al. 2015). Thus, we generated contact matrices at resolutions as high as 1 kb, but with a focus on 2 kb and 5 kb resolutions for downstream analysis, using the cooler (v0.10.3) framework, and applied iterative correction (ICE) for normalization via cooler balance. Across the samples, cooler balance assigned NaN weights to an average of 12.84% ± 0.26% of bins at 2 kb resolution (1,544,155 bins total) and 13.12% ± 0.30% of bins at 5 kb resolution (617,669 bins total).

Phased variant call files (VCFs) containing both single-nucleotide polymorphisms and structural variants were obtained from the HGSVC Phase 3 dataset, and were generated from the pav2 pipeline (docker://becklab/pav:latest; (Ebert et al. 2021). These VCFs were produced by aligning telomere-to-telomere haplotype assemblies generated from PacBio HiFi and Oxford Nanopore reads to GRCh38, calling variants with dipcall (v0.3) and phasing with WhatsHap (v2.8). The resulting phased VCFs were used to annotate deletions, insertions, and SNPs overlapping CTCF motifs and chromatin loops.

Single-molecule chromatin Fiber-seq data were produced using PacBio Revio HiFi sequencing (30 h movie time per SMRT Cell) at the University of Washington. Each sample achieved a mean genome-wide coverage of 33.5x, an average read length of 20.7 kb, HiFi yields of ∼103 Gb, and read-quality scores ranging from Q30–Q33 (Supplementary Table S2). Reads containing pre-annotated N⁶-methyladenine (m⁶A) modifications were aligned to GRCh38 using pbmm2 (v1.10.0). The resulting BAM files were phased using WhatsHap phase guided by corresponding variant calls. Processed reads were analyzed using the standard fibertools command suite (v0.6.4) pipeline. M⁶A bases and nucleosomes were identified using the following command:

~~~
ft extract GM19317/GM19317_m6_nuc.bam --reference \
 -q -a GM19317_all.tsv -n GM19317_nucleosomes.bed \
 --m6a GM19317/GM19317_m6a.bed -t 8
~~~

Footprinting was performed against known CTCF motif coordinates (JASPAR MA0139.1) using the following command:

~~~
ft footprint “GM19317_m6_nuc.bam” \
 --bed “GM19317H1_CTCF_COORDINATES_clean.bed” \
 --yaml “ctcf.yaml” \
 --out “GM19317H1_ctcf_footprint.bed”
~~~

For each motif, both the total number of fibers spanning the site and the number containing a CTCF-sized footprint were counted. The ratio of footprinted to total fibers defined the CTCF occupancy frequency for that site.

Per-base m⁵C CpG methylation calls for all samples were obtained from the HGSVC. Base-called reads were aligned to the telomere-to-telomere assembly of each sample using minimap2 (v2.30-r1287). Methylation profiles were phased using WhatsHap phase (v2.8) guided by dipcall-derived VCFs, yielding haplotype-specific methylation tracks. For downstream integration with Hi-C and Fiber-seq, the number of methylated CpG sites per fiber per genomic bin and per CTCF site was used as a proxy for total methylation signal.

### AB compartment analysis

A/B compartments were computed using the Cooltools (v0.7.0) eigs_cis function applied to ICE-balanced Hi-C matrices at 100 kb resolution using the following command:

~~~
eigvals, eig_df = cooltools.eigs_cis(
 GM19317_100000.cool,
 phasing_track=bins[[’chrom’, ’start’, ’end’, ’frac_gc’]],
 n_eigs=1
)
~~~

GC content was used as the phasing track to orient the first eigenvector (E1), such that positive values correspond to GC-rich, transcriptionally active (A) compartments. The resulting E1 eigenvalues for each genomic bin were compared between samples using pairwise MSE and sign concordance to quantify compartment similarity. A null expectation for MSE was estimated as the genome-wide variance of E1 values pooled across all samples, corresponding to the expected mean squared difference under random alignment of compartment eigenvectors. For visualization, E1 profiles across a representative 35 Mb region were plotted for all samples. Hi-C maps showing compartment state and inter-sample compartment changes were generated using matplotlib from sample cooler files. For all A/B compartment, TAD domain, chromatin loop, and chromatin interaction-related analyses in this manuscript, we highlight representative loci and regions that illustrate the general trends observed genome-wide.

### Chromatin accessibility with Fiber-seq

For all samples, Fiber-seq reads across both haplotypes were partitioned into 100 kb genomic bins (only relevant fragments of each read were kept in the case of reads that overlap multiple bins). For each read or fragment of a read, the total number of adenines and m⁶A methylated adenines were determined and summed across every read for every bin. The proportion of methylated adenines/total adenines within each bin was computed. For between-sample comparisons, these percentages were mean-centered within each sample to remove global shifts in modification levels and focus analyses on relative spatial variation along the genome. For each pair of samples, we computed the average MSE between mean-centered m⁶A levels at the same genomic bin across samples and sign concordance as the percentage of bins in which both samples showed deviations in the same direction (positive or negative relative to their respective means), excluding bins with 0 deviation. Final statistics reported averaged MSE and sign concordance across each pair of samples. A permutation-based null model was constructed where for each sample pair, m⁶A values from one sample were randomly permuted across bins 1,000 times while preserving the empirical value distribution, and the MSE was recomputed to generate a null distribution of similarity values.

### Identification of differential interactions using diffHic

Genome-wide differential chromatin interactions between the samples were identified using the diffHic package (v1.38.0) following the protocol detailed in the diffHic User’s Guide (https://bioconductor.posit.co/packages/devel/bioc/vignettes/diffHic/inst/doc/diffHicUsersGuide.pdf). The steps performed were:

1. Generate raw Hi-C contact matrices for the 5 samples at 1 kb, 2 kb, and 5 kb resolution using the cooler dump() function. These matrices were used as input for diffHic.
2. Filter out centromeres, telomeres, interactions < 10 kb, interactions > 1 Mb, and interactions where across all 5 samples, the sum of contacts was < 2.
3. Normalize for bias using the normOffsets() function, which allows for cross-sample pixel comparisons. Following the documentation, the correctedContact() was not implemented as ICE normalization is not necessary for cross-sample pixel comparisons.
4. Estimate dispersion and fitting the quasi-likelihood negative binomial GLM model to the data. This step was repeated with each iteration comparing one sample to the remaining samples, generating one list of differential interactions for each sample. Our setup kept deletions in the Hi-C dataset as 0 counts, allowing them to be identified as differential between samples.

The resulting interaction lists were filtered to retain pixels passing logCPM > –4 in at least one of the five sample and interaction distances between 10 kb and 1 Mb. The logCPM threshold was selected based on the distributions of logCPM and logFC values, optimizing retention of informative contacts while preserving the expected unimodal logFC distribution centered around zero at 5 kb resolution (Supplemental Figures 4-8). Across filtered loci, logFC standard deviations followed a negative binomial distribution with a mean σ = 0.36, reflecting that most interactions are conserved and a minority are differential. Significant interactions were identified using Bonferroni correction at each resolution, and genome-wide logFC profiles were visualized for each sample. The set of 367 significant interactions at 5 kb was used for mechanistic analyses.

### Mechanistic analysis of diffHic differential interactions

Four mechanisms were evaluated for their association with diffHic-identified differential Hi-C pixels (aka interaction-pairs):

1. Large-scale sequence differences between interaction bins
2. Small-scale sequence differences within interaction bins
3. m⁶A methylation
4. m⁵C methylation

For genetic variation, a leave-one-out (LOO) approach was used to identify the sample with the most extreme value at each differential pixel. For each interaction locus, the sample with a differential number of total bases changed (via SNPs, insertions, and deletions) both between interaction anchor bins and in interaction anchor bins was determined. Samples were considered differential if they exhibited a median absolute deviation (MAD)-based Z-score ≥ 2 and their LOO t-test p-value fell below a Bonferroni-adjusted 0.02 threshold. This was chosen as the threshold because it corresponds to strong outlier deviation in a MAD-based framework while maintaining sensitivity given the sample size (n = 5). For bins with a differential genetic alteration sample we quantified the match rate between that sample and the sample with the largest absolute value logFC. Statistical significance was assessed using a two-sided binomial test under a null probability of 0.2 (reflecting the 1-in-5 chance of a match by random expectation).

M⁶A methylation rates were computed by determining, for each fiber and each bin, the number of methylated adenines divided by the total adenines. Rates were averaged across fibers and across both interacting bins for each differential Hi-C contact. To normalize baseline differences between individuals, genome-wide expected methylation ratios were computed and compared to the observed ratios. Bins with observed log-ratios deviating from expectation by > 0.2 were flagged as differential, and the direction of the deviation was recorded. Because increased m⁶A methylation is expected to correlate with reduced chromatin contacts, loci were classified into four categories (+m⁶A, +logFC; +m⁶A, -logFC; -m⁶A, +logFC; -m⁶A, -logFC) based on whether the differential-methylation sample aligned with the minimum or maximum logFC value. The match rate of each set was calculated and statistical significance was assessed using a two-sided binomial test under a null probability of 0.2. This calculation was repeated while excluding loci where the identified differential sample contained a CTCF binding site within 10 kb of either interacting bin to differentiate the effects of low m⁶A values caused by closed chromatin from the effects of low m⁶A values caused by CTCF binding.

CpG methylation levels were obtained from phased ONT reads. For each resolution, per-bin methylation fractions were computed as the number of methylated cytosines divided by the total number of cytosines within that bin, separately for each haplotype (H1, H2). The methylation fractions of the two interaction bins were averaged to obtain a single value per haplotype per differential interaction. To identify whether a sample exhibited aberrant m⁵C levels at a locus, we applied a similar robust outlier framework as before. Across the 10 haplotypes, methylation values were converted into MAD Z-scores, and the haplotype with the largest absolute Z-score was considered a candidate differential sample. A leave-one-out (LOO) Z-score was then computed by recalculating the median and MAD excluding the candidate haplotype sample. A haplotype was marked as differential if both the MAD Z-score and the LOO Z-score were ≥ 2, and the direction of the deviation (“+” for higher methylation, “–” for lower) was recorded. For each differential m⁵C event, we compared the direction of methylation deviation with the Hi-C log fold-change (logFC) of the haplotype-sample. Because increased CpG methylation is expected to reduce chromatin contacts, loci were classified into four categories (+m⁵C, +logFC; +m⁵C, -logFC; -m⁵C, +logFC; -m⁵C, -logFC) based on whether the differential-methylation sample aligned with the minimum or maximum logFC value. For each category, we computed the proportion of loci in which the m⁵C deviation correctly predicted the logFC extremum and assessed significance using a two-sided binomial test with null probability 0.2.

To assess joint explanatory power, differential bins were annotated with any mechanism for which a differential sample was detected, and match-rate analyses (total bases changed between interaction bins and in interaction bins; +m⁶A, -logFC; +m⁵C, -logFC) were calculated. To evaluate independence, match-rate analyses were recomputed after removing any differential interactions explained by two or more mechanisms, demonstrating that each mechanism retains significance when controlling for the others. A Venn diagram generated using the venn Python package (v0.1.3) summarized the overlap between the match rates of each mechanism in bins that exhibited at least one differential mechanism.

Next, we computed the frequency with which each unique genomic bin appeared across all differential interactions and quantified the proportion that fell within 10 kb of a chromatin loop in the 2D Hi-C matrix as well as within homozygous bin-eliminating deletions, calculating the null through circular permutation (n = 1000) by randomizing differential interaction anchors at identical genomic distances along the chromosome and testing against fixed homozygous deletion intervals with empirical permutation p-values. We also clustered differential pixels within 10 kb × 10 kb of each other in the 2D Hi-C matrix and plotted the distribution of the number of differential pixels within each cluster, stratified by clusters within 10 kb of a loop or within homozygous deletions.

### CTCF site determination and PWM score calculation

Because 85% of chromatin loops form at CTCF sites (Rao et al. 2015), CTCF motif locations in each sample were derived by running the FIMO software (v5.3.0) on haplotype-resolved assemblies for each sample using the following command:

~~~
fimo CTCF.meme GM19317.vrk-ps-sseq.asm-hap1.fasta
~~~

The least stringent CTCF motif instance across all haplotypes retained a FIMO p-value of 3.85 × 10^-6^, which approaches the suggested FIMO p-value threshold of p < 1 × 10^-6^ (Dozmorov et al. 2022) Because this process yielded 55,422 ± 1,180 CTCF sites per haplotype, consistent with prior estimates of expected human CTCF site counts (Chen et al. 2012), we did not further filter CTCF sites to those satisfying p < 1 × 10^-6^, which would have eliminated ∼29,166 sites from each haplotype. We note that although FIMO uses a default p-value threshold parameter (“--thresh”) of p < 1 × 10^-4^, FIMO also uses a default “--max-stored-scores” parameter of 100,000 sites, and when those 100,000 sites are filled, the weakest sites are truncated to make room for new sites. This led to slight variation in the maximum p-value retained by each haplotype (3.48 × 10^-6^ – 3.85 × 10^-6^) despite running FIMO with identical commands to that shown above. Importantly, we validated that within each haplotype, this truncation process did not remove any CTCF sites that satisfied the maximum p-value after running FIMO with --max-stored-scores set to 1,000,000 sites. Results were mapped back to the reference genome GRCh38 using the Long Read Aligner software (v1.3.7.2). This process successfully mapped ∼90% of CTCF sites for each sample back to the reference genome, averaging ∼49,983 sites per haplotype.

The FIMO output also included the base sequence for each CTCF site. The position weight matrix (PWM) scores were calculated for each sequence for each haplotype. A PWM probability matrix with per-base probabilities at each position in the motif was constructed from JASPAR’s 2018 MA0139.1 CTCF frequency matrix. For each haplotype-resolved CTCF site, the corresponding 19 bp sequence was scored by summing the position-specific probability weights associated with each base and dividing the total by the motif length to obtain a motif probability score, which enables direct comparison across the ten haplotypes in our dataset and integrates naturally with downstream quantitative models. Importantly, the minimum PWM scores across all 10 haplotypes were exactly the same (0.514), indicating that the slight variation in maximum p-values between haplotypes did not change which CTCF sequence motifs were considered for downstream analysis.

For each CTCF sequence, the number of bases mutated compared to the canonical 15 bp CTCF consensus sequence 5′-NCANNAGRNGGCRSY-3′ (Hashimoto et al. 2017) was calculated. To accomplish this, the 15 bp window was aligned across each haplotype sequence to identify the highest-scoring local match. Within the best alignment, bases that violated the consensus rules were counted as mismatches, and the number of mismatches per sequence was recorded.

### CTCF site m⁵C methylation

CpG methylation was quantified at single-base resolution using ONT-derived per-nucleotide methylation calls for each haplotype. This data was processed to determine, for each genomic position, the number of reads overlapping that cytosine and the number of reads supporting a methylated call for each sample and haplotype. For each position, a methylation fraction was computed as the number of methylated cytosines / the total number of cytosines. For every haplotype-resolved CTCF motif, all CpG positions falling between the start and end coordinates were retrieved. Multiple CpG methylation events on the same site were averaged, producing a site-level methylation profile for every motif and haplotype.

In parallel, a hidden Markov model (HMM) was applied genome-wide to classify each CpG position as hypomethylated or hypermethylated. These HMM state calls were merged into the CTCF annotations using the same coordinate-based procedure, yielding for each haplotype-specific CTCF site a categorical label of hypomethylated, hypermethylated, or mixed (if CpG positions within the motif belonged to different HMM states). Similar results could be achieved from the Fiber-seq m^5^C reads, however the ONT reads were used as orthogonal support.

### CTCF site accessibility

The m⁶A rates (the number of m⁶A adenines / the total number of adenines) were calculated within each 1 kb bin for each sample. The average m⁶A rate of the four bins (two upstream, two downstream) surrounding each bin that contained a CTCF site was calculated. To account for technical biases from sequencing, the m⁶A rates were Z-scored within sample and sigmoid-transformed into a score between zero and one, representing how accessible the chromatin near each CTCF site was compared to the other CTCF sites in the sample. Z-scores were calculated within CTCF sites rather than genome-wide as downstream analysis was focused only on CTCF sites.

### Quantifying correlations between genetic and epigenetic factors and occupancy

Each CTCF site across all ten haplotypes was annotated with by PWM score, the number of sequence mutations relative to the consensus motif, the mean CpG methylation level, the CpG methylation state (hypomethylated, mixed, hypermethylated), the average accessibility rate, and the Fiber-seq–derived CTCF occupancy value. To enable cross-haplotype comparisons of the same CTCF site, we matched CTCF sites that mapped back to the GRCh38 reference genome within 50 bp together. Global pairwise correlations between each feature and occupancy were computed to assess individual relationships. To quantify the joint association between sequence strength, accessibility, and CpG methylation and their relationship to CTCF occupancy, we fit an ordinary least squares (OLS) regression model using PWM score and methylation as predictors of occupancy and calculated the correlation between observed and fitted values. PWM scores were min–max normalized to the range [0,1], whereas methylation values were transformed to a z-score across all motif instances. Rows containing missing values for any of the predictors or occupancy were excluded. Because the goal of this analysis was association rather than prediction, the model was fit to the full dataset without a train–test split. Global significance of the association model was assessed using the F-test for linear regression with two predictors. All analyses were performed in Python using pandas, numpy, scikit-learn, scipy, matplotlib, and seaborn.

### Loop discovery using Mustache and HiCExplorer

The following commands were used to discover loops, with appropriate length parameters and adjustments per sample:

~~~
hicDetectLoops -m GM19317_5000.cool -o \ GM19317_HICEX_5000_01.bedpe --maxLoopDistance 2000000 \
 --windowSize 10 --peakWidth 6 --pValuePreselection 0.1 \
 --pValue 0.1
mustache -f GM19317.mcool -r 2kb -pt 0.1 \
 -o GM19317_2000_01.tsv -p 50 -st 0.7
~~~

Chromatin loops were initially called on each resolution of each sample using both Mustache (v1.3.3; MUST) and HiCExplorer (v3.7.5; HICEX) software with p-values of both 0.1 and 0.01. Mustache loops called at 1 kb and 2 kb resolutions included an additional st parameter which was set at 0.7 following recommended settings. All HiCExplorer loops called with the hicDetectLoops() function included additional parameters of maxLoopDistance = 2000000, windowSize = 10, and peakWidth = 6. To compare loop sets, we computed Jaccard indices between callers and between samples after expanding each loop to a ±10 kb neighborhood to account for minor positional differences in peak localization. The same comparisons were also performed on two independent GM19317 technical replicates at 10 kb resolution.

We also calculated Jaccard index values with respect to varying sequencing depth by splitting the read pairs of unbalanced GM19317 cooler files at 1 kb, 2 kb, 5 kb, and 10 kb resolutions in half, balancing both output halves, and calling loops on both halves for all four resolutions using Mustache (HiCExplorer failed to call loops on either of the halves at any of the resolutions). Jaccard index values were then calculated between halves at each resolution, as they were above.

### CLASH

CLASH is a method that assigns a loop strength score to a provided Hi-C locus based on the structure of Hi-C signal at that locus, accounting for the roughly radial symmetry of decaying contacts from the loop center (Eagen 2018). The input for CLASH is pooled loop call sets of genomic coordinates where a previous method has called a chromatin loop. For each candidate loop, bins within 10 kb of the diagonal were excluded to avoid including windows with invalid ranges of separation and crossing over the diagonal of the Hi-C matrix. Then, for each called loop, the maximum balanced contact count within a dynamic search window ranging from a 5×5 matrix (±2 bins) for short-range loops (<100 kb) up to an 11×11 matrix (±5 bins) for long-range loops (>200 kb) was selected as the refined center of the candidate loop.

For each refined loop center, we extracted a local Hi-C submatrix with adaptive sizing. Shorter loops (loop anchor separation < 35 kb) always used a 5×5 window (±2 bins). For longer loops, the following procedure was implemented:

1. Start at a minimum radius r = 2, corresponding to the 5x5 ring centered around the loop center.
2. The average balanced contact count of pixels, ë, in the rings corresponding to r, r+1, and r+2 was determined.
3. One sided Welch’s t-tests were used to determine if λ_r_ > λ_r+1_ or if λ_r_ > λ_r+2_ with a p-value ≤ 0.05.
4. R is incrementally increased and the process is repeated from step 2. Matrix expansion stops when three consecutive radii fail the test in step 3, returning r = max(2, the max r that passed the test in step 3).
5. The (2r + 1, 2r + 1) matrix centered around the refined loop center is extracted.

Six features were included to describe each putative loop:

1. Loop anchor separation: The genomic distance (bp) between the two interacting bins.
2. Center value: the balanced contact count of the center pixel for each matrix.
3. Loop prominence: The Z-score of the size of matrix extracted around the loop via our adaptive procedure, computed relative to the matrices extracted for the other samples at the same locus.
4. Zero fraction: the ratio of pixels within the matrix that have a balanced contact count = 0.
5. Smoothness, determined by:

a. Calculating the Euclidean distance of each pixel in the matrix from the loop center and grouping pixels at the same distance.
b. Calculating the mean contact intensity of each radial distance to produce a radial intensity profile.
c. Fitting a univariate spline to the intensity profile.
d. Calculating the mean squared residual between the observed radial intensities and the fit spline.
6. LC scores: the maximum local contrast between a candidate loop and its background, calculated from Gaussian-smoothed interaction maps across multiple spatial scales. Our implementation was heavily based on Mustache’s implementation (Roayaei Ardakany et al. 2020) and calculated 23 LC scores across between 24 scales at each putative loop, of which we chose the maximum.

An XGBoost binary classifier model, CLASH, was trained using these six features to predict a loop or no loop, minimizing binary cross-entropy. The model used 300 trees with a maximum depth of 4, a learning rate of 0.05, and subsampling of 0.8 for both rows and features. Training data consisted of 1,000 putative loops across 200 loci (199 of which were identified by either/both of Mustache or HiCExplorer in at least one sample, with the other locus, chr7:96076000-97028000, being added after visual observation of the Hi-C map despite not being called in either Mustache or HiCExplorer in any sample). Loops across these 200 loci were manually assigned binary labels of 0 (no loop) and 1 (loop) based on visual inspection of the corresponding Hi-C maps, with 845 true loops spread out across 199 loci and 155 false loops spread out across 71 loci. Our curated set incorporated loci with and without CTCF anchor sites, and with loop anchor separations spanning genomic distances of 36,000-1,214,000 bp. Similar procedures of manual loop presence validation have been used by other machine learning methods for loop detection (Salameh et al. 2020).

Model performance was evaluated using out-of-fold prediction from five-fold cross-validation with grouping by locus to prevent different loops from the same locus appearing in both training and validation sets. Feature importance was determined using SHAP (Lundberg and Lee 2017). For each classification the logit was min-max scaled to yield a continuous CLASH score, and the decision boundary was determined by mapping the classification probability that maximized Youden’s J (true positive rate - false positive rate) to min-max scaled space.

Although the primary benefit of CLASH is that it provides this continuous score, we partitioned CLASH scores into five groups to facilitate their visual interpretation on Hi-C maps. Roughly, scores from 0-0.4 represent the absence of a loop, scores from 0.4–0.55 represent non-loop regions with limited enrichment, scores from 0.55-0.65 – which straddle the decision boundary of ∼ 0.6 (calculated from the operating point that maximized Youden’s J) for this dataset – represent ambiguous loops, scores from 0.65 - 0.85 represent loops, and scores from 0.85 - 1 represent strong loops; these ranges are intended as visual guides rather than strict thresholds.

### CLASH Validation

We validated CLASH’s testing AUROC by permuting labels and retraining the model, and we compared CLASH’s AUROC classification performance to XGBoost and Logistic Regression models that exclude LC scores as a feature, and only include LC scores as feature, as well as the classification performance of raw LC scores. To enable fair comparisons, we tested the recall for CLASH and the Logistic Regression LC at a matched false positive rate. We also computed the recall and false positive rate of both Mustache and HiCExplorer on the loci comprising the training set.

To demonstrate CLASH’s ability to harmonize across samples using the decision boundary corresponding to the maximum Youden’s J from the full dataset, we evaluated how many training loci initially called by Mustache and HiCExplorer were worsened, improved, or perfectly improved by CLASH loop predictions (evaluated out-of-fold) and compared this to the LC model performance.

To validate the usage of CLASH scores, we computed the error severity of out-of-fold misclassifications on the training set, defining error severity as the normalized distance from the decision boundary relative to the extent of the incorrect region, and compared the results to those of LC model scores. We also performed orthogonal tests to externally validate CLASH loop-scoring performance by computing Pearson’s correlation efficients between CLASH scores with CTCF occupancy globally and within-locus across samples, and compared this to the LC model score correlations.

### Quantifying the correlation between CTCF occupancy and CLASH loop-score

To quantify how CTCF protein occupancy globally relates to chromatin loop strength, CLASH-scored loops were first filtered to retain only those in which both loop-associated interaction bins contained a CTCF site within 10 kb. For each loop, the CTCF occupancy values of its two CTCF sites were averaged to generate a single occupancy value. If CTCF occupancy was missing, the CTCF occupancy value from the other site was used, and if both were missing then the datapoint was excluded from analysis. This per-loop occupancy metric was then paired with the corresponding CLASH loop score, and Pearson’s correlation coefficients were computed across all loci and samples to assess the global relationship between CTCF binding and loop strength.

We also quantified the distribution of Pearson’s correlation coefficients between CTCF occupancy and CLASH loop-strength between samples at each locus. Loci with fewer than four samples containing both valid occupancy and loop score measurements were excluded. We collected all per-locus correlations and performed a binomial sign test to calculate statistical significance. All analyses were conducted in Python using Pandas, Numpy, Scipy, Seaborn, and Matplotlib.

### Quantifying the correlation between PWM scores, m⁵C methylation, CTCF occupancy, and CLASH loop scores

For each chromatin loop in each sample, PWM scores and CpG methylation levels from the two loop-associated CTCF sites were averaged to generate per-loop measures of motif strength and methylation. If multiple CTCF sites were located within 10 kb of a loop anchor, we chose CTCF sites by sequentially optimizing for CTCF orientation (such that anchors would have convergent orientations (Rao et al. 2015), motif strength (PWM scores), and minimal distance from the anchor bin. This process resulted in 79% of our CTCF-associated loop set being anchored by CTCF sites in convergent orientations. Orientation relationships between paired loop-anchor CTCF sites were highly conserved across haplotypes for the same loop locus, with only 0.2% of loops exhibiting variation in orientation state across haplotypes. These values were paired with CLASH loop scores and occupancy values from earlier, and global Pearson’s correlation coefficients were computed to quantify the individual relationships between motif strength and loop intensity. Due to the skewed distribution of methylation values, we stratified loop score distributions by methylation values and used Mann Whitney U tests to quantify the individual relationships between methylation and loop intensity.

We next quantified the number of loci where the CLASH score varied by 0.3, indicating a substantial difference in loop strength, and where occupancy, PWM scores, and methylation data existed for at least 3 of the 5 samples. From this set, we calculated the proportion of loci where the observed difference in loop strength could be explained (defined as having a correlation of ≥ 0.25) by each of the three mechanisms.

To test whether PWM scores and methylation values provided explanatory power beyond CTCF occupancy, we compared the correlation between occupancy and loop-strength to the multiple correlation of a combined association model incorporating occupancy, PWM scores, and CpG methylation using Steiger’s test for dependent correlations, which accounts for shared outcomes and predictor non-independence. This analysis provided a direct statistical assessment of whether adding genetic (PWM) or epigenetic (m⁵C) features improved explanatory power relative to occupancy alone. We performed an analogous comparison using PWM scores and methylation values as the only predictors to evaluate their joint contribution independent of occupancy.

### Mediation analysis

To quantify how much of the effect of CTCF motif strength (PWM score) and CpG methylation on loop strength is transmitted through CTCF occupancy, we performed a global product-of-coefficients mediation analysis. For PWM scores, we only pooled observations from variant CTCF loci. We treated PWM scores or CpG methylation as the predictor, CTCF occupancy as the mediator, and CLASH loop strength as the outcome. For each predictor, mediated effects were estimated using linear regression to obtain the product of the predictor-occupancy (a) and occupancy-loop (b) paths, along with the corresponding direct (c’) and total (a×b+c’) effects and the proportion of the total effect mediated. The percent mediated was calculated as (a×b)/(a×b+c’) and averaged across all 5 samples. Uncertainty was measured using a pooled nonparametric bootstrap in which loci were resampled within each sample and per-sample indirect effects were recomputed, averaged across samples for each bootstrap iteration (1,000 iterations), and compared to the mean total effect to derive bootstrap confidence intervals.

### Structural variation

We investigated whether large SVs (insertions and deletions) that introduce or remove CTCF binding sites lead to measurable changes in chromatin loop formation. Structural variant calls for each sample haplotype, aligned to the GRCh38 reference genome, were filtered to retain only those altering at least 19 bp (the length of the CTCF consensus motif). FIMO motif scanning was then performed on each SV sequence to identify the number and positions of CTCF motifs gained or lost. We only considered CTCF sites that passed a FIMO p-value threshold of 3.85e-6, matching the least stringent p-value threshold used to identify non-variant CTCF sites across the haplotypes. These SV-associated motif changes were subsequently intersected with CLASH-derived loop scores to assess whether the addition or removal of CTCF sites corresponded to loop strengthening or weakening.

For each loop locus, and for each sample, we then determined whether either haplotype carried an SV that added at least one new CTCF site (insertion) or removed at least one CTCF site (deletion) within either loop-associated interaction bin. Across the individuals, this yielded two groups of loop strength measurements at each locus: (i) samples whose haplotypes contained a CTCF-altering SV at the interaction bin, and (ii) samples without such an SV. For each locus and for each SV class, we computed the change in loop strength as the difference between the mean CLASH loop score of the SV group and the non-SV group. Statistical significance of these differences genome-wide was assessed using a Wilcoxon test comparing loop scores between SV and non-SV samples at each locus. We later relaxed the FIMO p-value threshold to 1e-5 as part of an exploratory search to include additional loop-associated CTCF variants.

### Alpha Genome

To evaluate whether sequence variation alone is sufficient for current sequence-to-map models to accurately predict observed loops and assess whether they could benefit from incorporating methylation and CTCF occupancy data in addition to sequence data, we used Alpha Genome (commit 6973cfe32c8f7d692350b2063c1f7cd611d1cd4f, (Avsec et al. 2026) to predict contact maps for all our haplotypes based on their genetic sequence. We focused on generating maps around the three loci that served as examples for instances where CTCF occupancy (Supplementary Figure 32), SNPs in CTCF sites (Supplementary Figure 34), and methylation in CTCF sites (Supplementary Figure 35) affected loop formation.

To generate the maps, we first extracted the 1,048,576 bp of GRCh38 centered around the midpoint of each pair of interacting bins that formed each loop that we were focused on. We then adjusted the input sequence using variant calls for each of our haplotypes, re-centered the locus of interest, and re-adjusted the overall length of the sequence to be 1,048,576 bp. The adjusted sequence of each haplotype was then used to predict the corresponding contact map using the following function call:

~~~
output = dna_model.predict_sequence(sequence=hap_seq,
requested_outputs=[dna_client.OutputType.CONTACT_MAPS],
ontology_terms=[“EFO:0002784”])
~~~

The output was then plotted as a Hi-C map in a manner similar to the original examples, including cropping the dimensions of the output to visualize the same region as the original examples. We were unable to extract a matrix-like quantity from the output in order to use CLASH to score the predicted loops.

