## Supplementary Methods for "CLASH (Chromatin Loop Across-sample Score Harmonizer) quantifies the relative contributions of genetic variation, methylation, and CTCF occupancy on chromatin loop strength across individuals"

**Table S1.** Hi-C sequencing and mapping summary statistics across the five samples.

| Sample | GM19317 | GM19347 | HG02666 | HG01457 | HG03248 |
|---|---|---|---|---|---|
| Total read pairs (millions) | 1813.6 | 1946.7 | 1440.9 | 1789.8 | 1449.8 |
| % unmapped | 31.40% | 32.30% | 28.50% | 31.10% | 30.10% |
| % one-sided | 10.30% | 10.20% | 10.70% | 9.90% | 10.00% |
| % two-sided | 58.30% | 57.50% | 60.80% | 59.00% | 59.90% |
| % duplicated | 12.80% | 14.20% | 12.10% | 19.20% | 11.90% |
| Unique read pairs (millions) | 824.9 | 842.8 | 701.1 | 712 | 695.9 |
| Cis interactions (count) | 658065376 | 648963062 | 576398312 | 596436231 | 616383635 |
| Trans interactions (count) | 166854257 | 193862696 | 124683354 | 115560791 | 79549255 |
| FF (1-2kb) | 24.60% | 25.10% | 24.90% | 24.60% | 24.70% |
| RF (1-2kb) | 20.60% | 20.90% | 21.10% | 20.60% | 21.30% |
| FR (1-2kb) | 30.00% | 28.90% | 29.10% | 30.00% | 29.20% |
| RR (1-2kb) | 24.80% | 25.10% | 24.90% | 24.80% | 24.80% |
| Coverage (x) | 175.76 | 188.65 | 139.64 | 173.44 | 140.50 |
| Suggested map resolution (Rao, bp) | 2500 | 2450 | 2950 | 2900 | 2950 |

**Table S2.** Fiber-seq summary statistics across the five samples.

| Sample | GM19317 | GM19347 | HG02666 | HG01457 | HG03248 |
|---|---|---|---|---|---|
| Coverage (x) | 34 | 31.1 | 33.8 | 35.3 | 33.3 |
| Mean read length (bp) | 20850 | 22591 | 24462 | 21151 | 14236 |
| Hifi yield (gbp) | 105 | 96 | 105 | 109 | 103 |

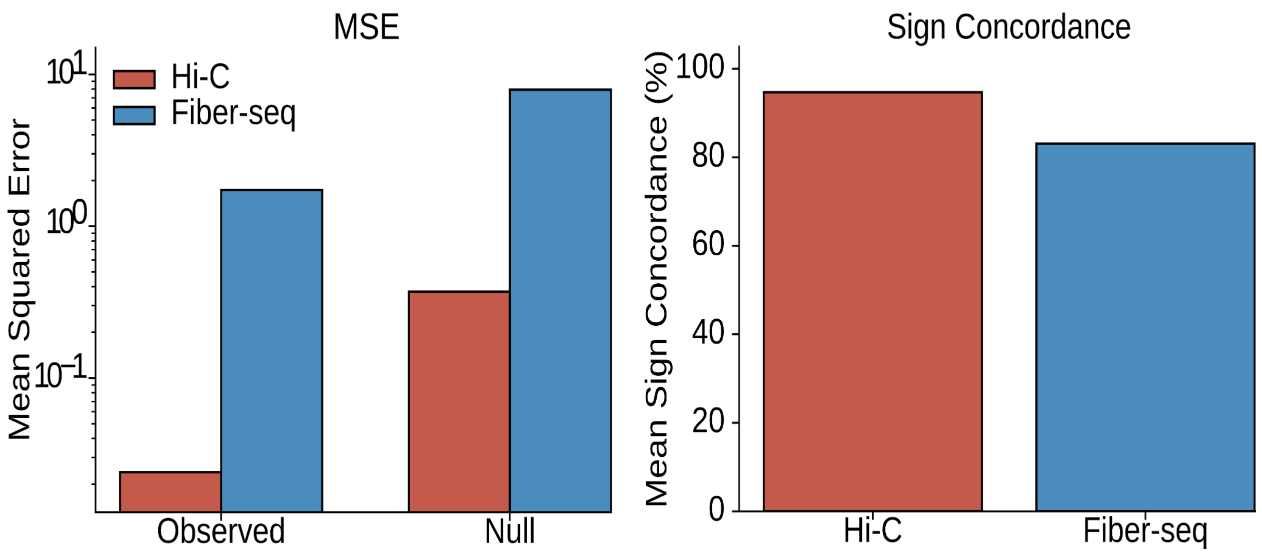

**Supplementary Figure 1. Pairwise comparisons of A/B compartment eigenvector values (E1) and accessibility profiles across all five samples.** Hi-C A/B compartment profiles computed at 100 kb resolution show high similarity with an average MSE of 0.024 ± 0.006 (red) and sign concordance, calculated as the fraction of bins with the same E1 sign, of 94.7% ± 0.9% across sample pairs. The null MSE of 0.371 was defined by the expected MSE if A/B compartments between samples were not correlated and calculated as E1 variance. Similarly, mean-centered average Fiber-seq accessibility profiles at 100 kb resolution show high similarity, with an average pairwise MSE of 1.731 ± 0.988 (red) and sign concordance of 83.1% ± 2.2% across sample pairs. MSE was calculated by mean-centered per-bin signals ($m^6A$ methylated adenines/total adenines), and the sign at each bin was determined as the deviation from the mean of that sample. The null MSE of 7.929 was calculated by randomly permuting mean-centered per-bin accessibility across genomic bins.

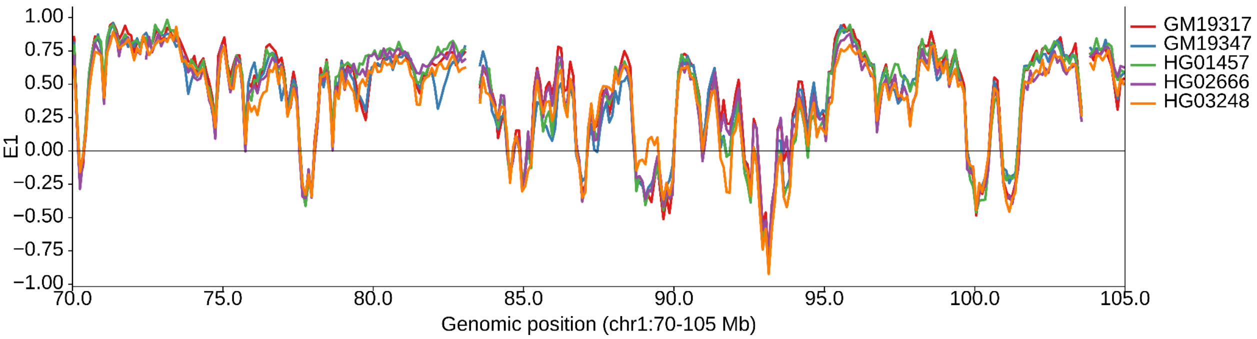


**Supplementary Figure 2. E1 compartment eigenvector profiles at 100 kb resolution for all five samples plotted across a representative region of chromosome 1 (chr1:70–105 Mb).** Colors indicate GM19317 (red), GM19347 (blue), HG01457 (green), HG02666 (purple), and HG03248 (orange), and samples exhibit largely similar eigenvector profiles. The region chr1:81,800,000-82,600,000 shows a decrease in GM19347 E1 scores compared to the other samples, and this can be attributed to GM19347 H1 containing >5× more total bases deleted in that region than the other 9 haplotypes.

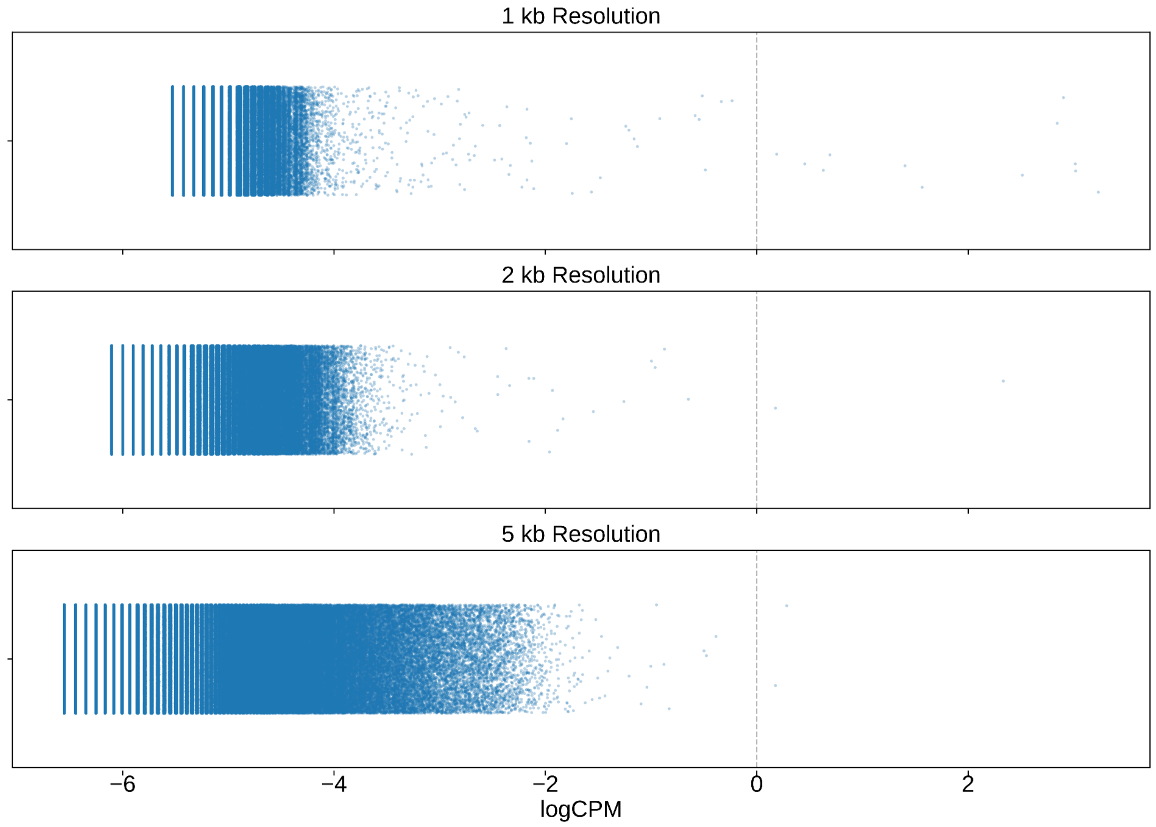


**Supplementary Figure 3. Distribution of logCPM values across all loci for GM19317 at 1 kb, 2 kb, and 5 kb resolution.** At 5 kb resolution, using a logCPM threshold of -4 retains sufficient data points to support statistical power. Thus, we used the set of data points at 5 kb resolution filtered for datapoints with logCPM > -4 for downstream QC and analysis.

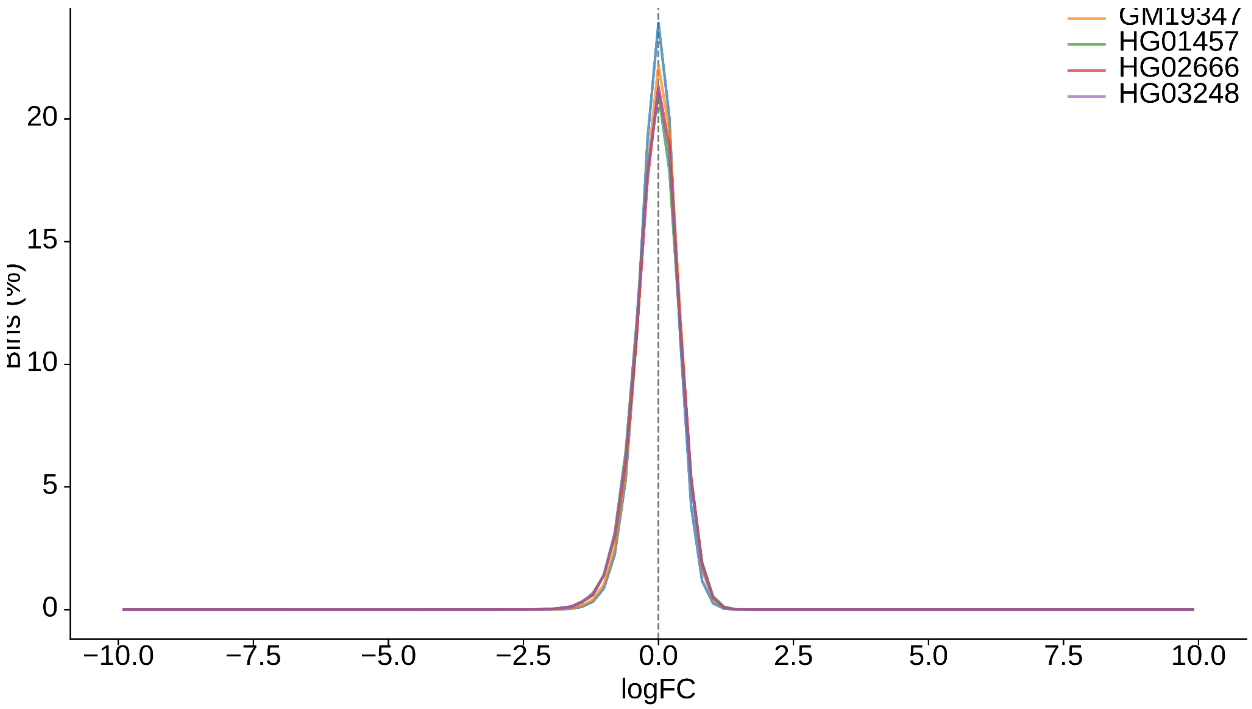


**Supplementary Figure 4. Distribution of filtered logFC values for each sample, showing the expected unimodal peak centered near zero at 5 kb resolution.**

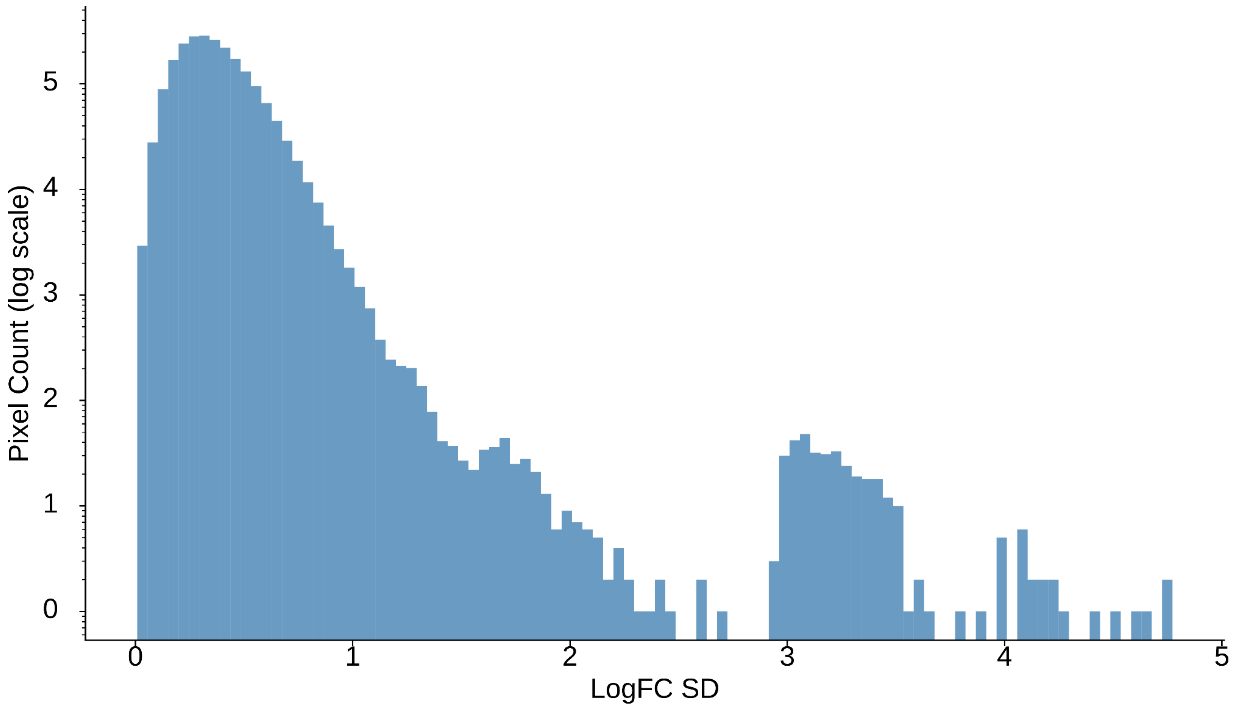

**Supplementary Figure 5. Distribution of logFC standard deviations across filtered pixels at 5 kb.** The distribution approximates a negative-binomial with most interactions conserved (mean σ = 0.36), and a minority showing strong differential signals.

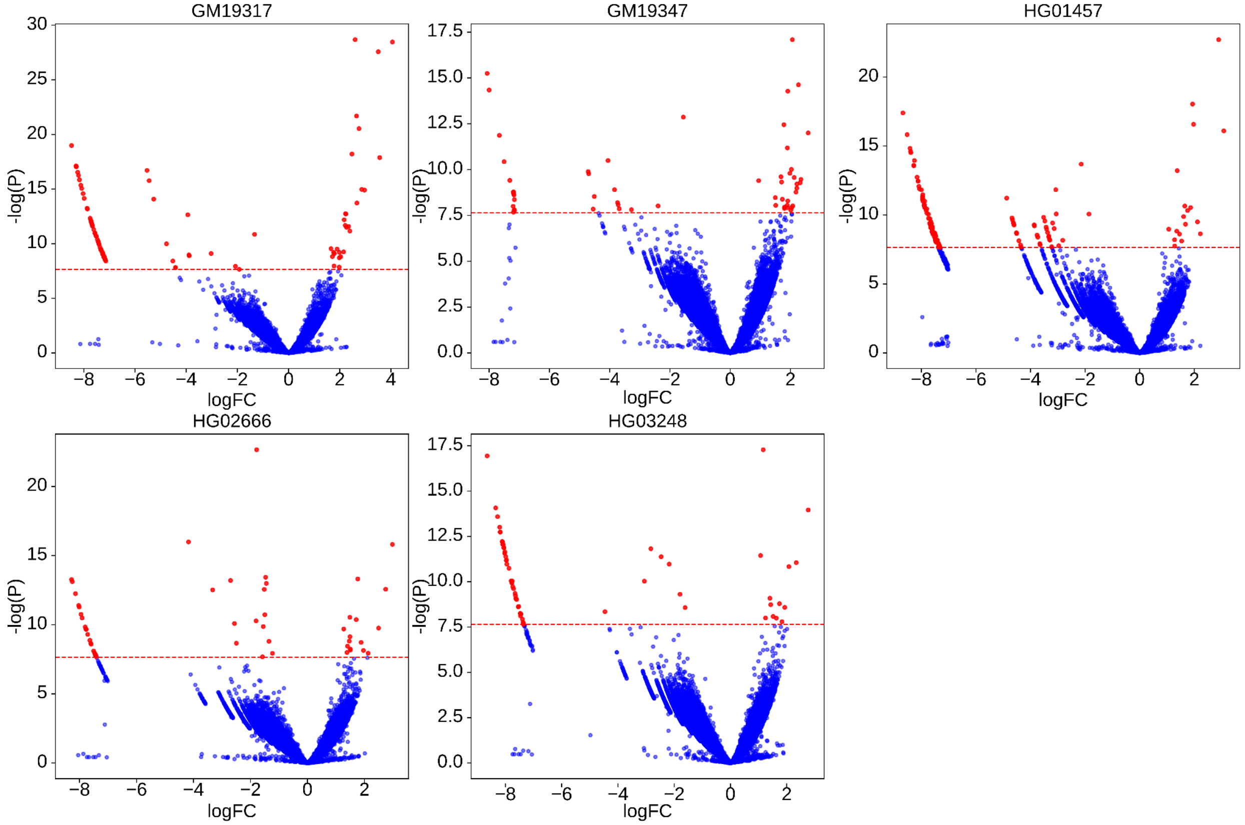


**Supplementary Figure 6. LogFC versus log(p-value) for filtered pixels at 5 kb resolution.** Bonferroni multiple-test correction was applied with Bonferroni $p = 2.3 \times 10^{-8}$, and pixels that passed the threshold are colored red.

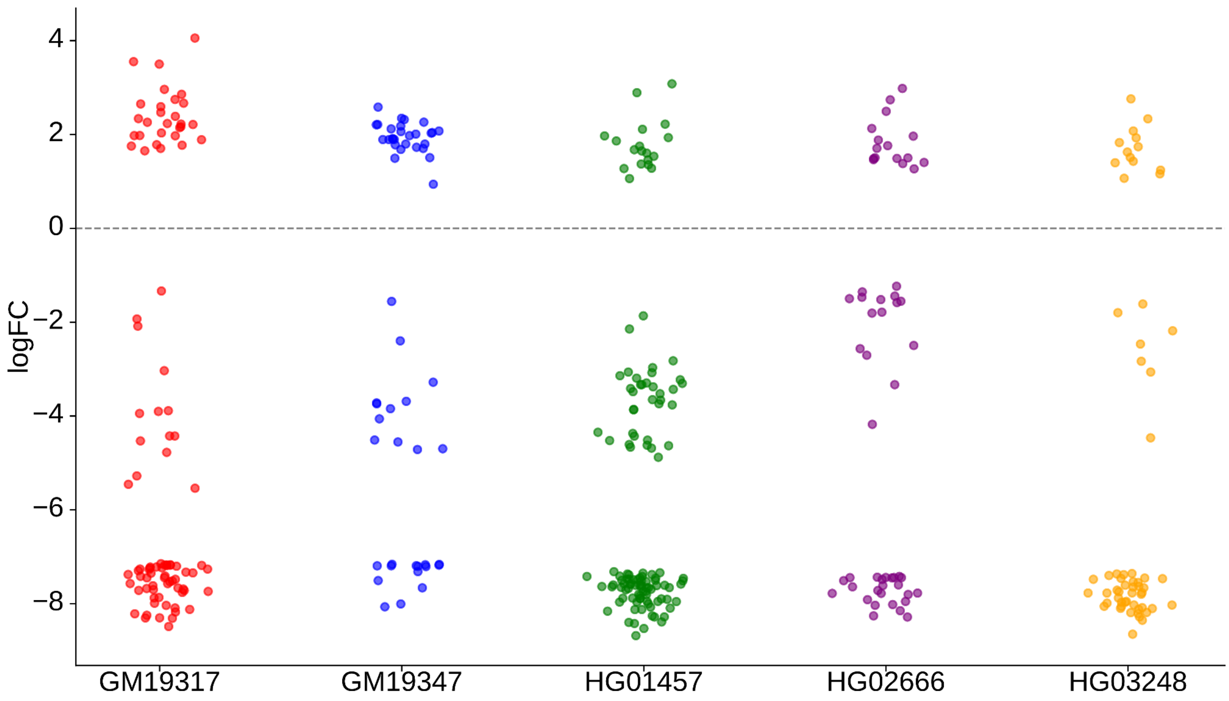


**Supplementary Figure 7. LogFC values for each filtered differential interaction for each sample at 5 kb resolution that pass the Bonferroni p-value.**

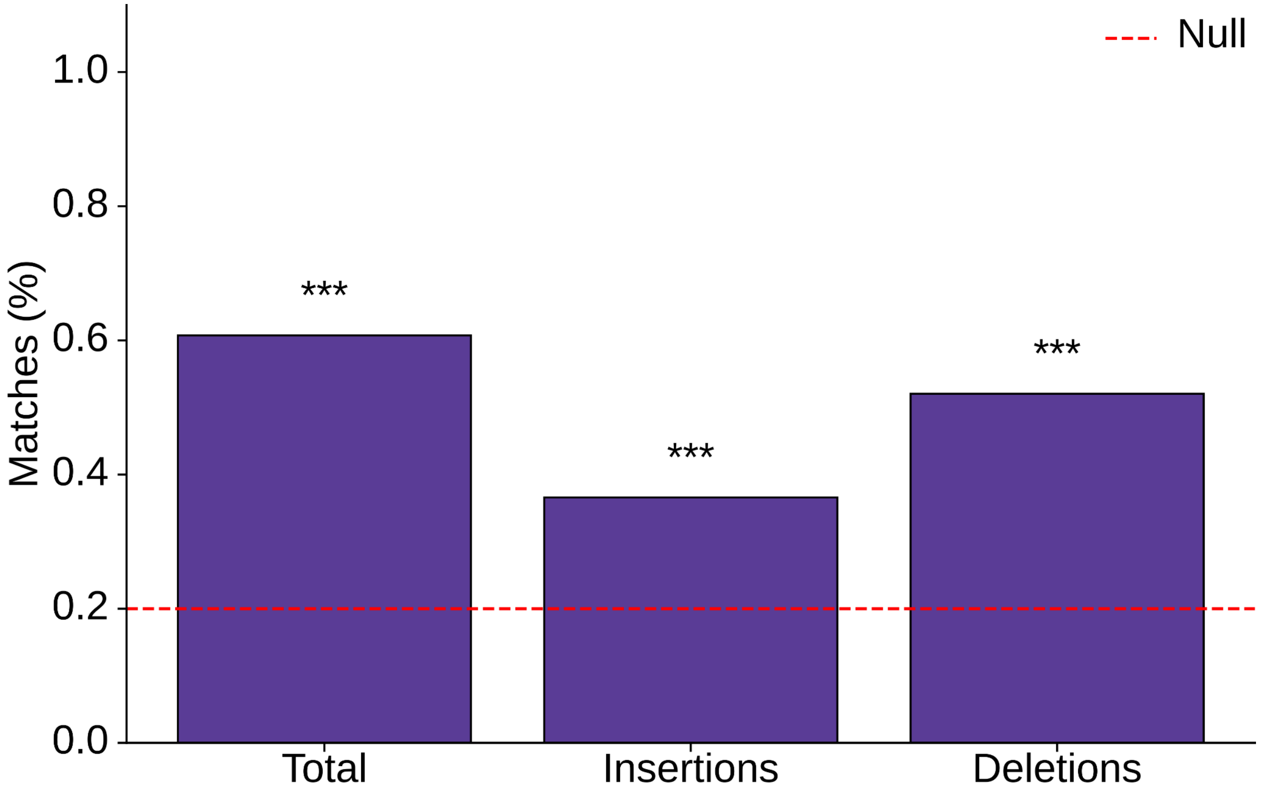


**Supplementary Figure 8. Match rate of differential loci where the sample with the most sequence changes between interacting bins also has the maximum log-fold change of contacts.** Match rates are compared to a 0.2 random expectation (5 kb).The match rate for total base changes is 60.7% ($n = 107$, $p = 3.95 \times 10^{-20}$, binomial test assuming independent effects of variant and fold-change). Significance levels are denoted as $p < 0.05$ (*), $p < 0.01$ (**), and $p < 0.001$ (***).

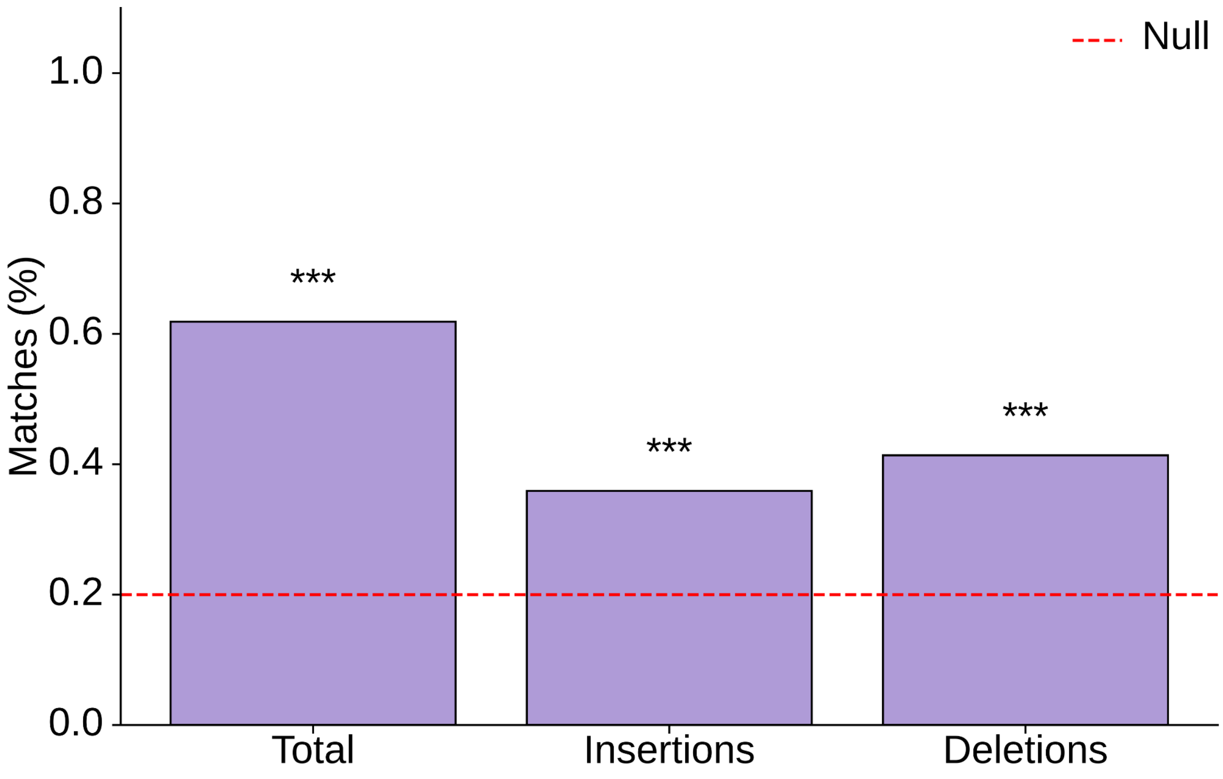


**Supplementary Figure 9. Match rate of differential loci where the sample with the most sequence changes inside bins also has the greatest log-fold change.** Match rates are compared to a 0.2 random expectation (5 kb).The match rate for total base changes is 61.9% ($n = 118$, $p = 4.25 \times 10^{-23}$, binomial test). Significance levels are denoted as $p < 0.05$ (*), $p < 0.01$ (**), and $p < 0.001$ (***).

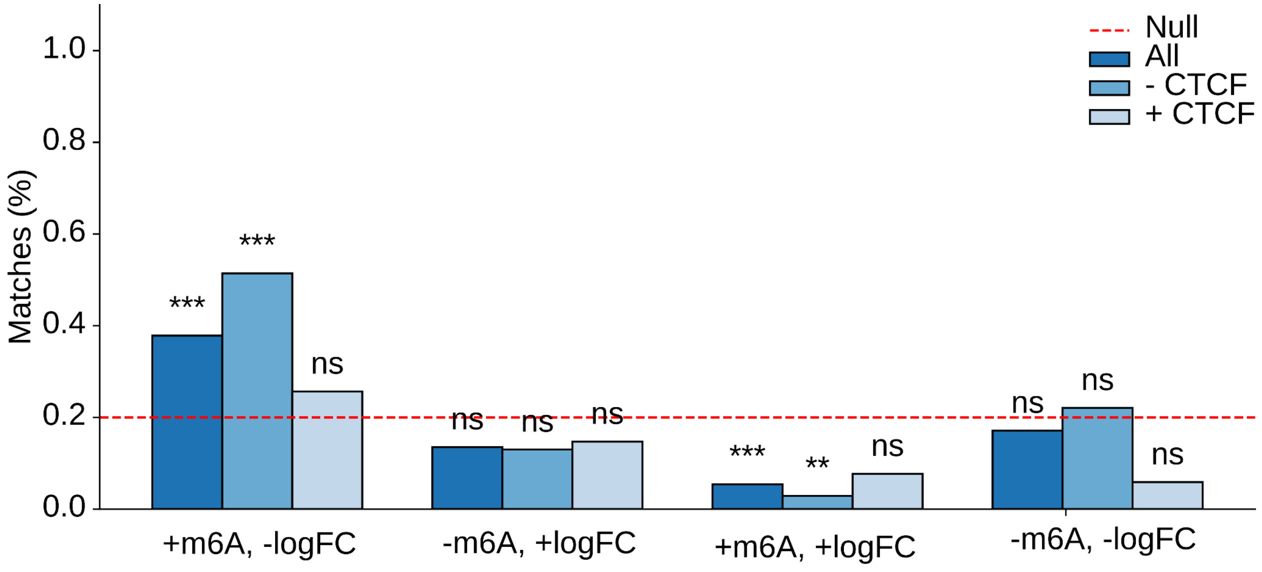


**Supplementary Figure 10. Match rates of differential loci in accessible chromatin (dark blue: all loci; blue: loci without a CTCF site within 10 kb of either interacting bin; light blue: loci with a CTCF site within 10 kb of either interacting bin) where the sample with the highest (+) or lowest (-) logFC value matches the sample with the highest (+) or lowest (-) m$^6$A methylation level compared to the null 0.2 under random matching at 5 kb resolution.** The match rate is defined as the count of sites where the sample with the highest measure of chromatin accessibility (m$^6$A) matches the sample with the lowest logFC. For all loci, the match rate is is 37.8% ($n = 74$, $p = 3.80 \times 10^{-4}$, binomial test), and when excluding differential contacts where decreases in m$^6$A methylation are associated with CTCF binding, the match rate is 51.4% ($n = 35$, $p = 3.42 \times 10^{-5}$, binomial test). Significance levels are denoted as $p < 0.05$ (*), $p < 0.01$ (**), and $p < 0.001$ (***).

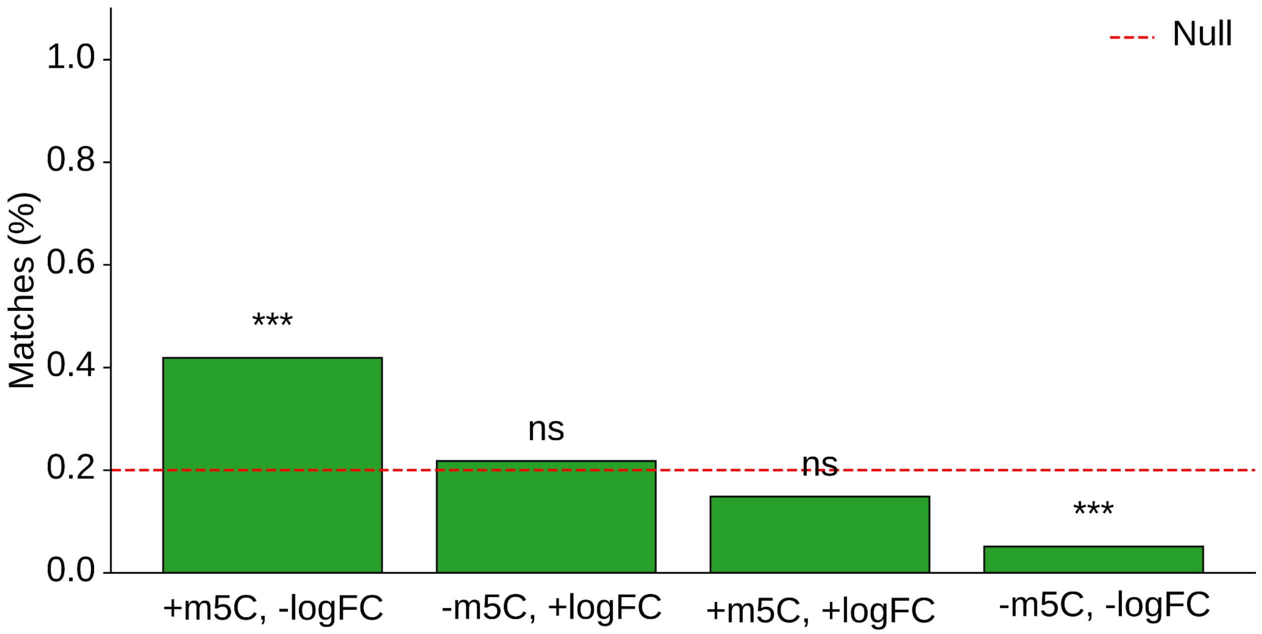


**Supplementary Figure 11. Match rates of differential loci with differential methylation.** Matches are defined as differential loci with differential methylation where the highest (+) or lowest (-) differential $m^5C$ methylation level is the same as the sample with the highest (+) or lowest (-) logFC. Significance is measured assuming independence (probability of match = 0.2) at 5 kb resolution. The match rate between the sample with the lowest logFC and highest $m^5C$ is 41.9% ($n = 74$, $p = 1.58 \times 10^{-5}$, binomial test). Significance levels are denoted as $p < 0.05$ (*), $p < 0.01$ (**), and $p < 0.001$ (***).

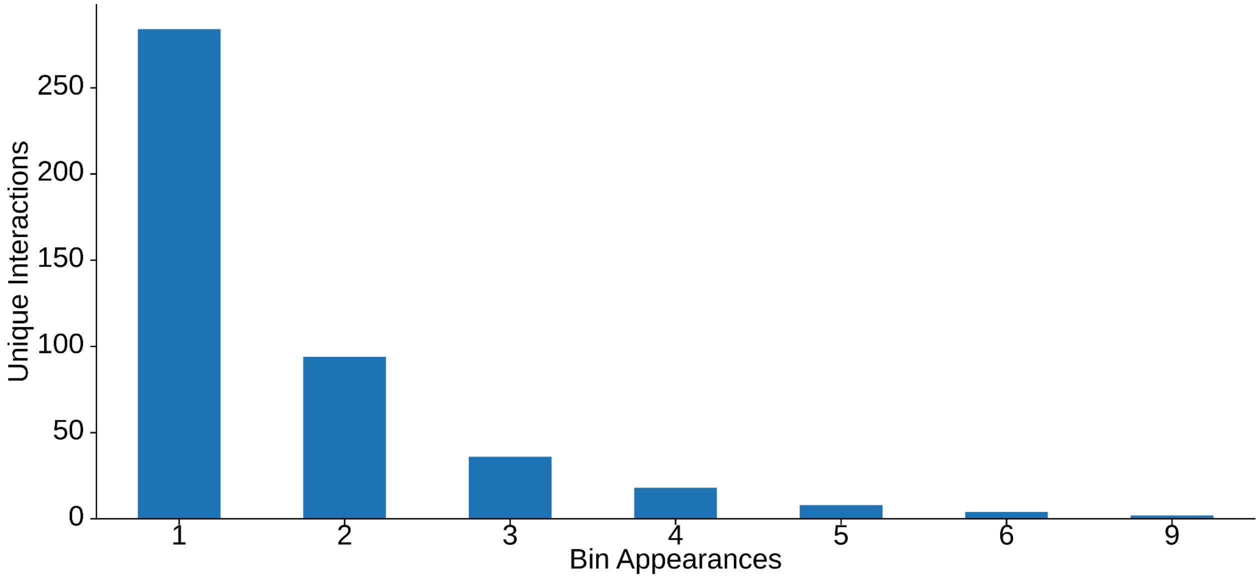


**Supplementary Figure 12. Frequency distribution of diffHic identified bins among the 367 differential pixels identified at 5 kb resolution across the genome.**

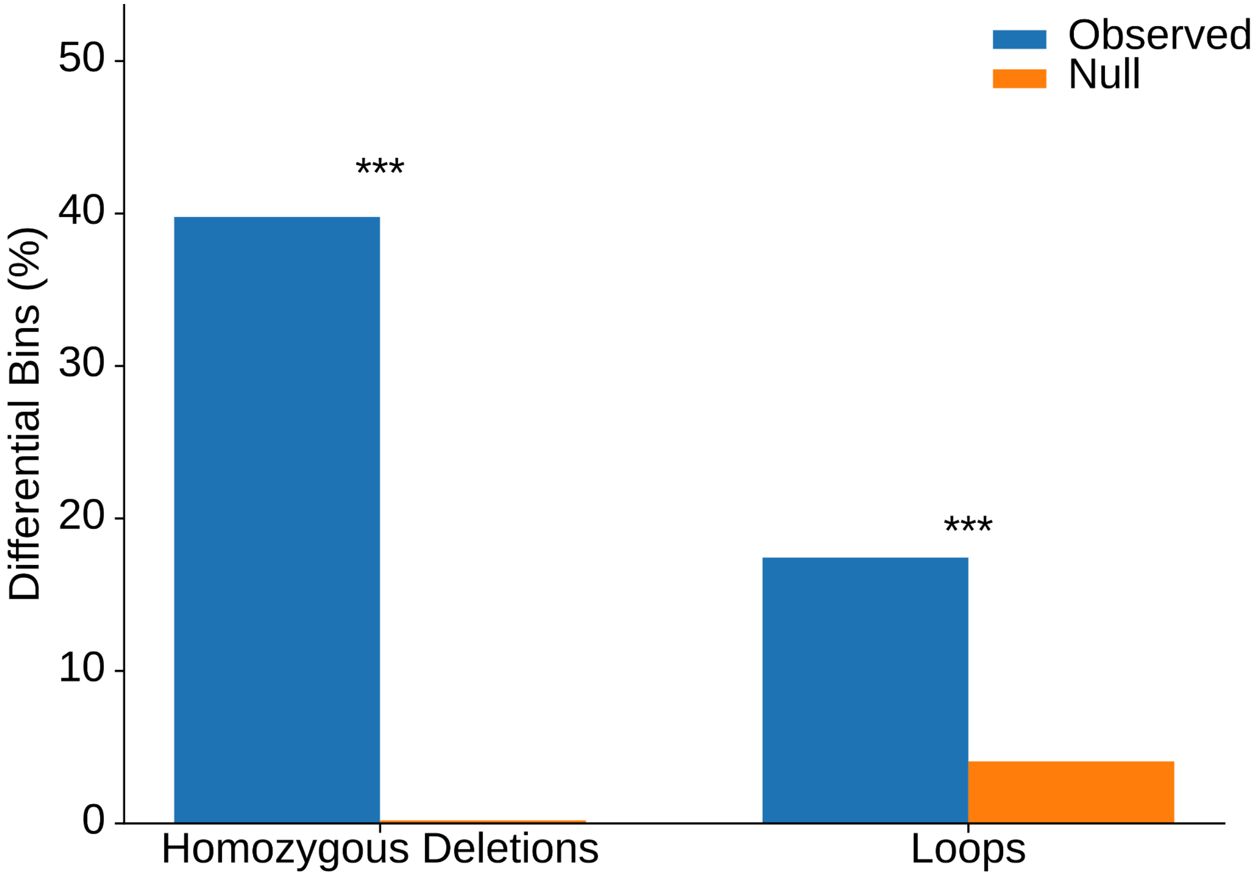


**Supplementary Figure 13. Proportion (blue) of differential pixels located within homozygous deletions (40%) or within 10 kb of chromatin loops (17%) at 5 kb resolution.** Null proportions (orange) for loops (n = 1000) were calculated through circular permutation (n = 1,000) randomizing loop anchor positions at identical genomic distances with empirical permutation p-values. Null proportions for deletions (n = 1,000) were calculated by circularly permuting differential interaction anchors along each chromosome and testing against fixed homozygous deletion intervals. For both, the $p < 0.001$. Significance levels are denoted as $p < 0.05$ (*), $p < 0.01$ (**), and $p < 0.001$ (***).

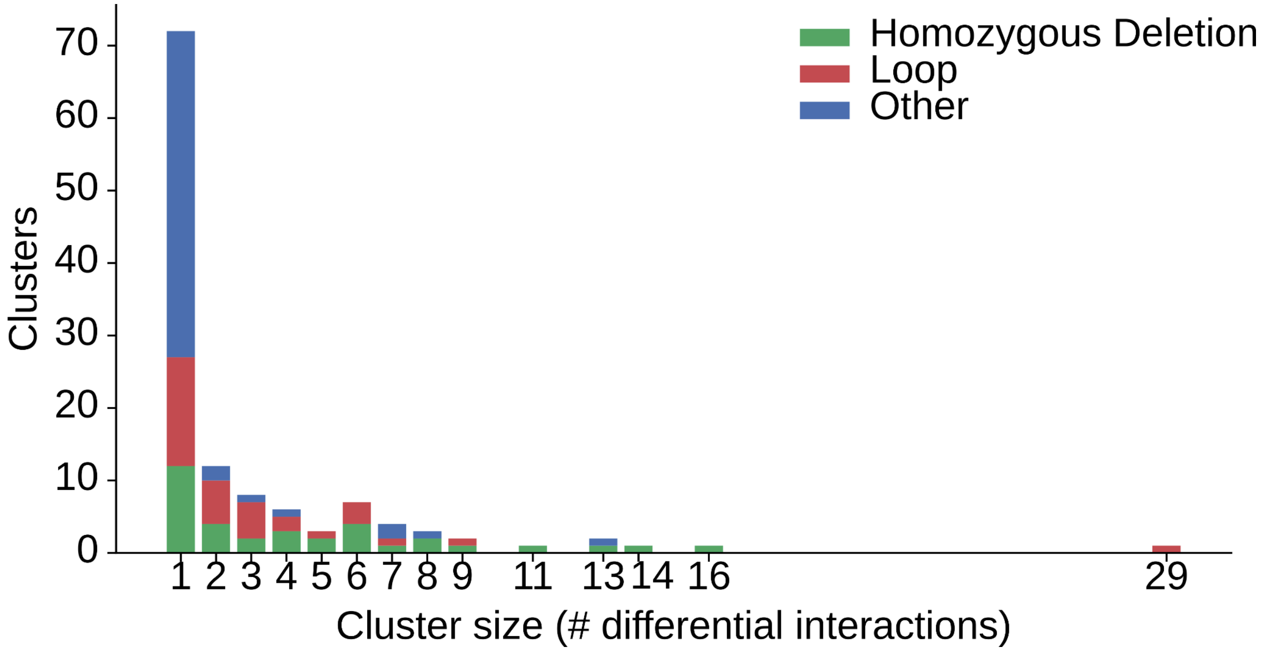


**Supplementary Figure 14. Number of diffHic clusters associated with homozygous deletions (green), loops (red), and neither deletions nor loops (blue) at 5 kb resolution, stratified by number of differential pixels within clusters (n = 367 pixels, as 123 distinct clusters).** Clusters were formed by grouping differential pixels located within a 10 kb × 10 kb window of each other in the 2D Hi-C matrix. Clusters associated with both a deletion and a loop were classified as being associated with loops.

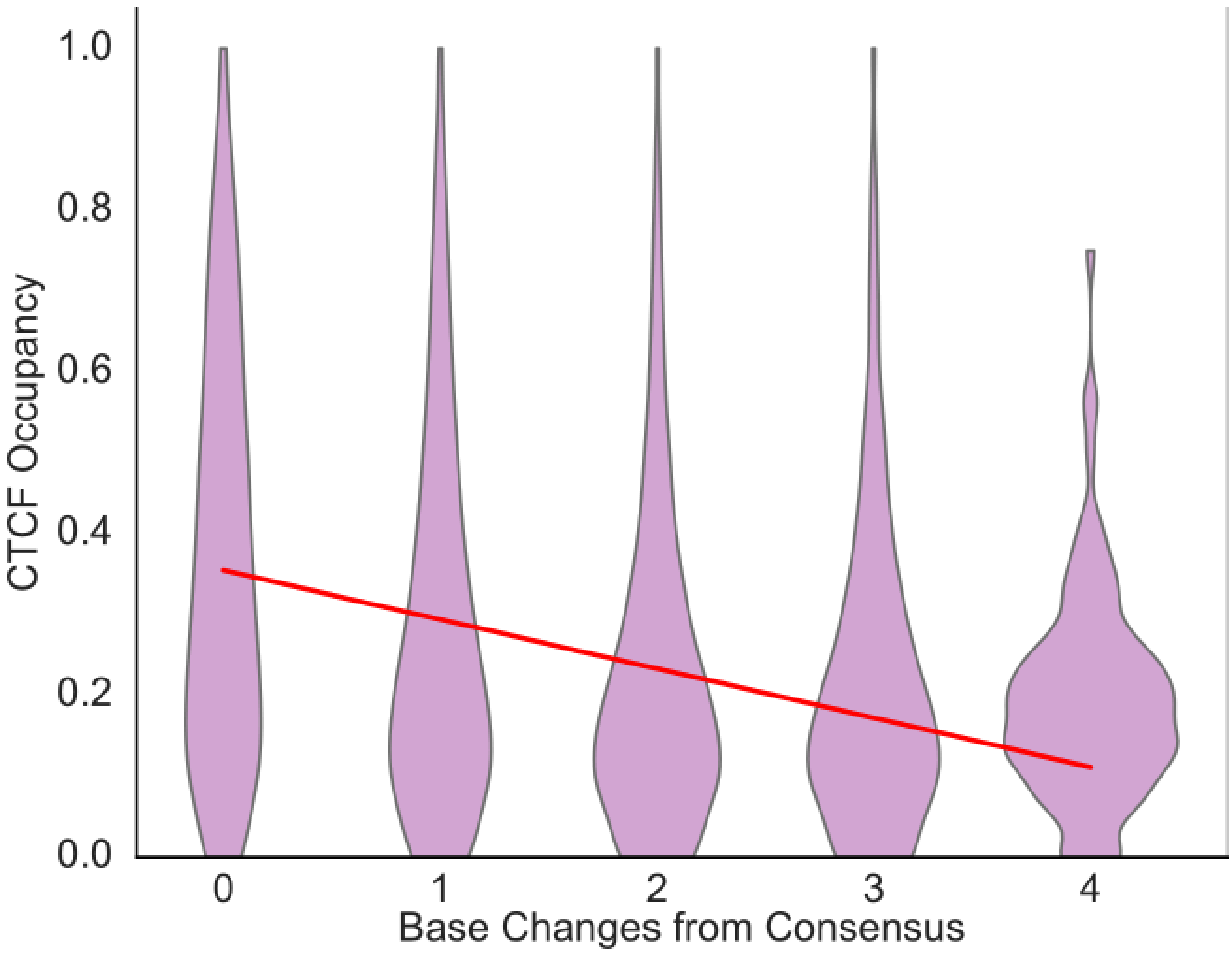


**Supplementary Figure 15. Correlation between the number of base substitutions within each CTCF motif and CTCF occupancy.** Pearson's r = -0.21 (n = 262,465 sites; $p < 2.2 \times 10^{-308}$). Significance levels are denoted as $p < 0.05$ (*), $p < 0.01$ (**), and $p < 0.001$ (***).

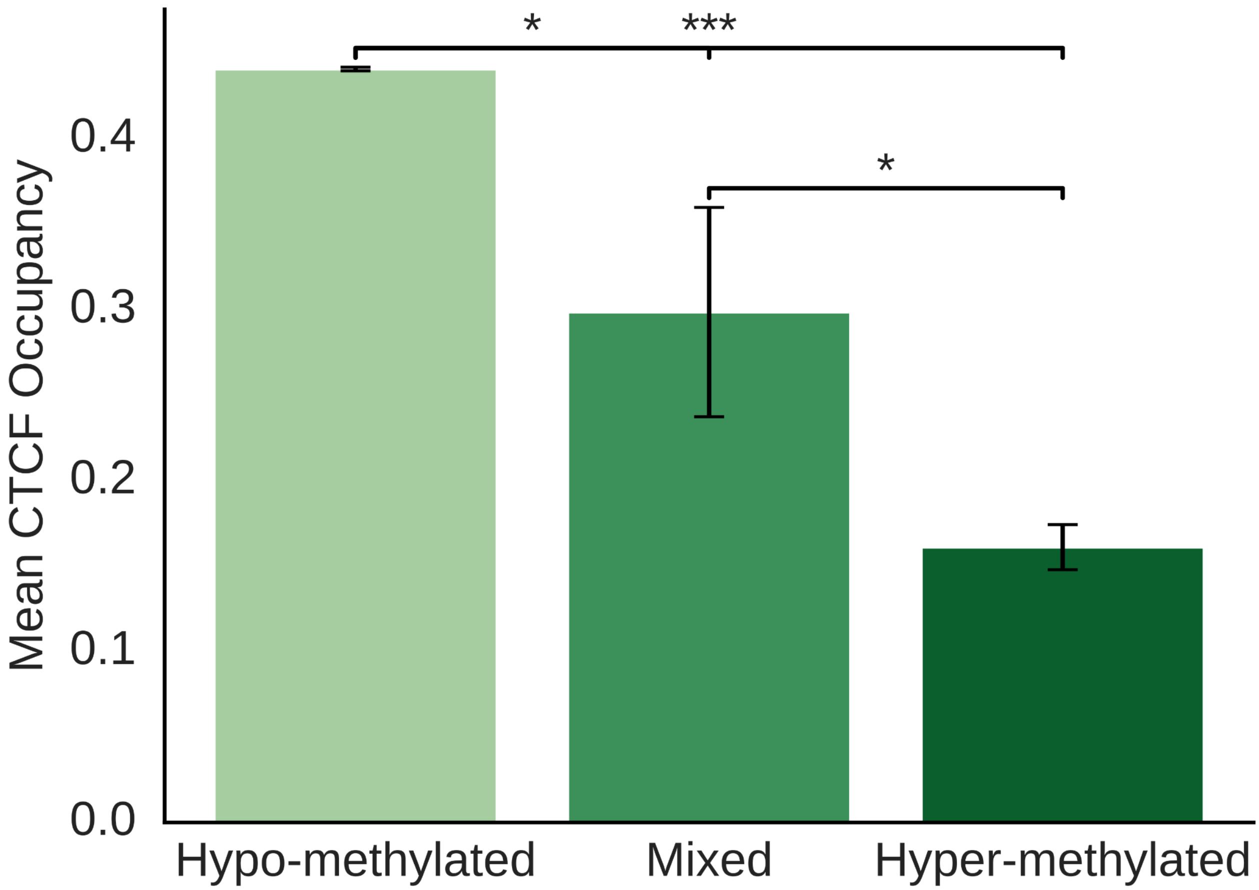


**Supplementary Figure 16. Haplotype-resolved relationship between CTCF m$^5$C CpG methylation state and CTCF occupancy.** There is a clear decrease in mean occupancy from hypo-methylated sites to mixed sites to hyper-methylated sites (hypomethylated n = 58,094 sites; hypermethylated n = 140 sites; mixed n = 14 sites; Mann Whitney $p = 4.4 \times 10^{-39}$). Significance levels are denoted as $p < 0.05$ (*), $p < 0.01$ (**), and $p < 0.001$ (***).

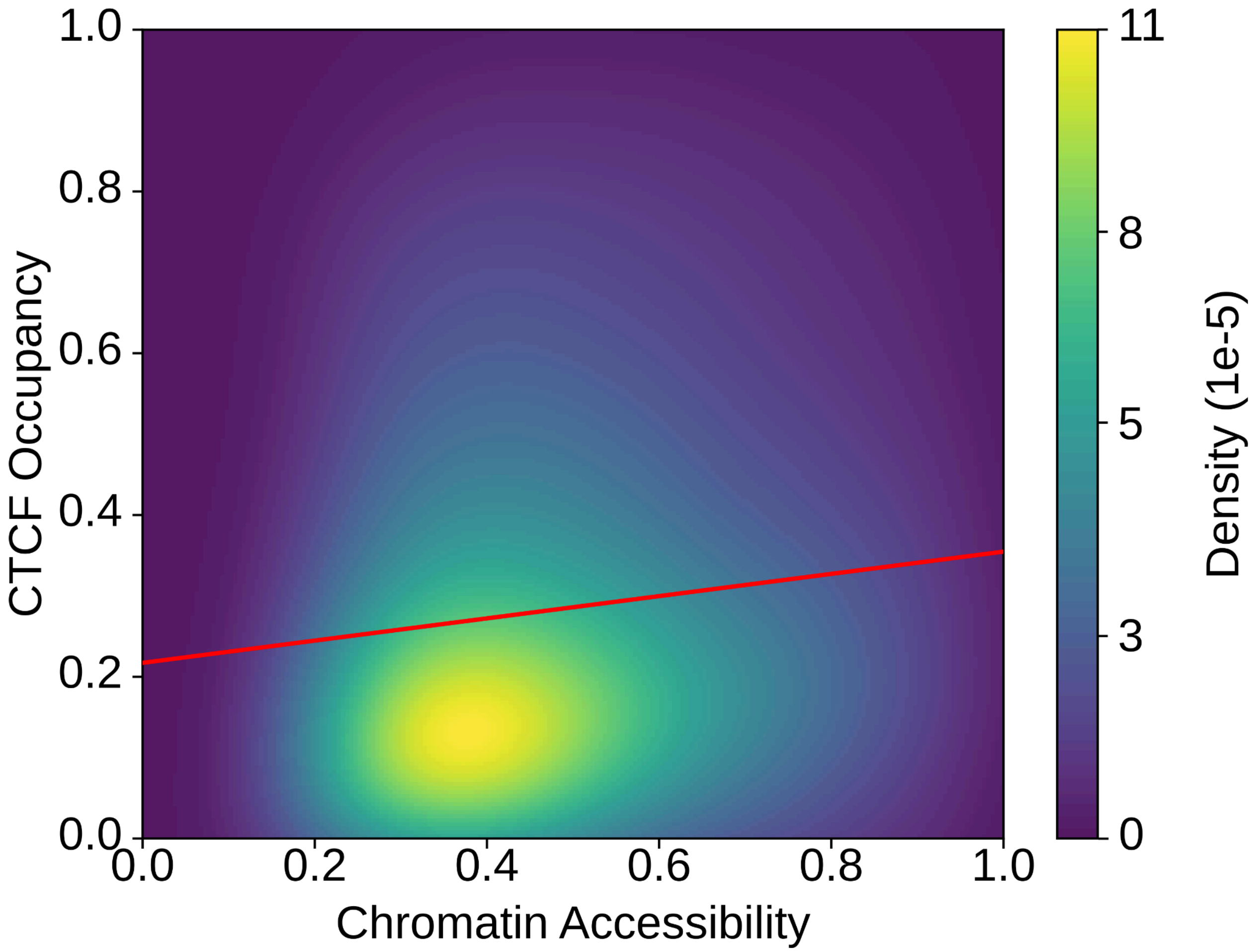


**Supplementary Figure 17. Correlation between local chromatin accessibility and CTCF occupancy (Pearson's r = 0.11, n = 262,282, $p < 2.2 \times 10^{-308}$).** Chromatin accessibility was determined from averaged $m^6A$ methylation rates from the 2 kb upstream and downstream of each 1 kb bin containing a CTCF site.

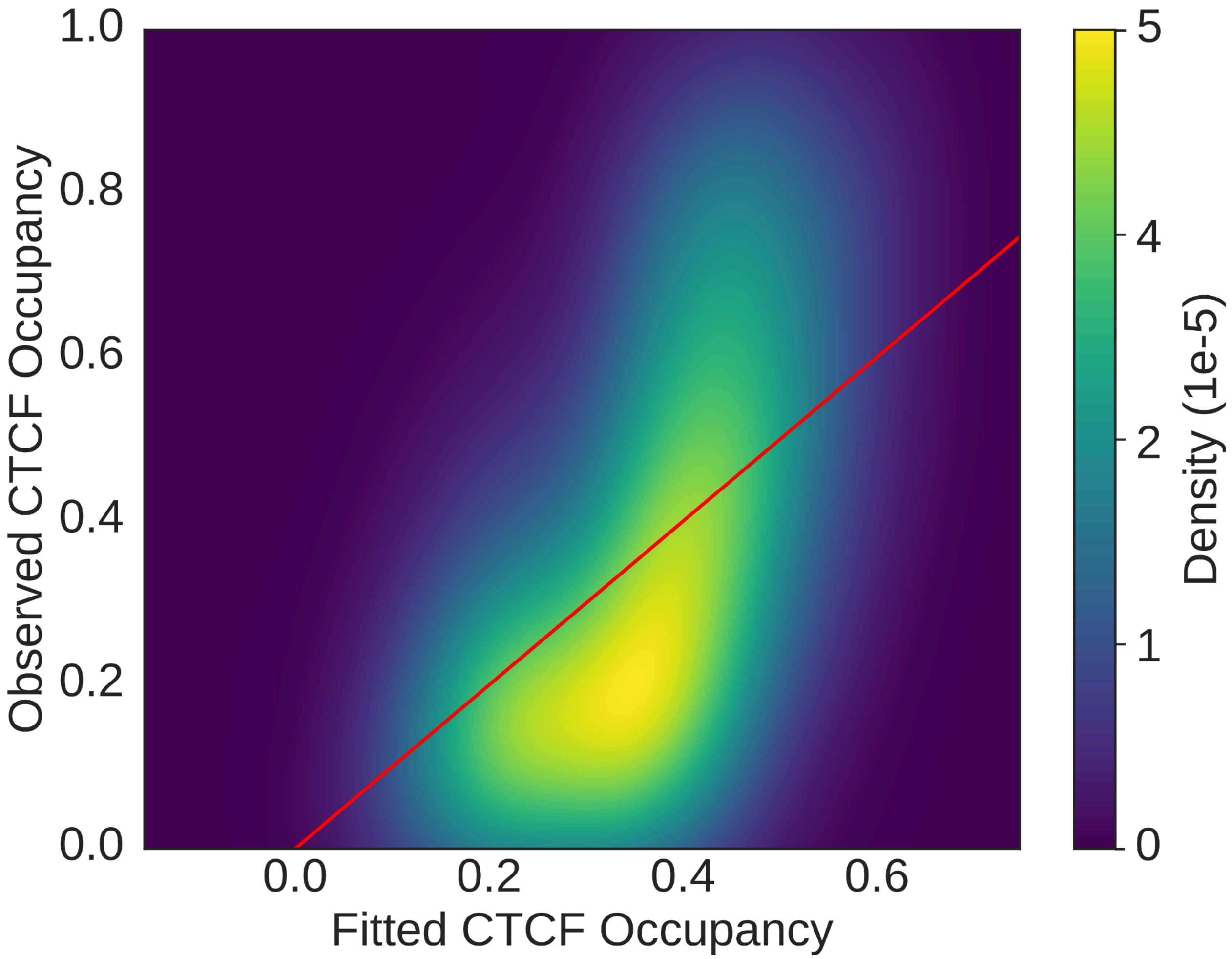


**Supplementary Figure 18. Linear model estimating the combined effects of PWM score, m[5]C methylation, and local chromatin accessibility on CTCF occupancy using ordinary least squares regression (n = 90,531 sites).** All predictors showed significant associations with occupancy (PWM β = +0.408; m[5]C β = −0.085, accessibility β = +0.143), and the model explained a moderate fraction of variance (Pearson's r between observed and fitted values = 0.50, F test  p = 1.1 × 10-16). Density estimation was evaluated on a 300 x 300 grid.

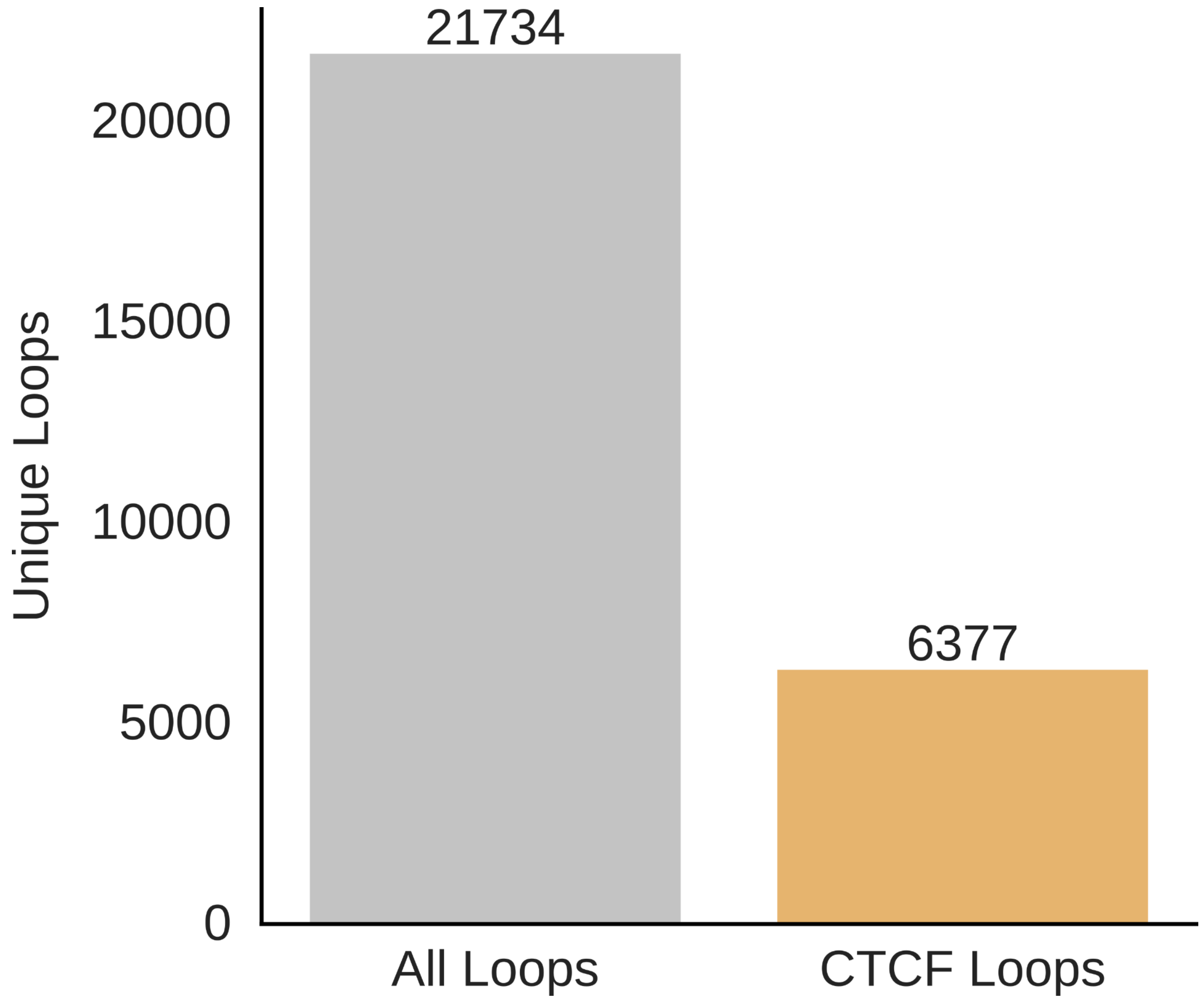


**Supplementary Figure 19. Total number of unique loci with at least one loop call in the pooled callset, identified by Mustache and HiCExplorer across all five samples at 2 kb resolution using a p-value threshold of 0.1.** The loop callset contained 21,734 loops (grey), and among these, 6,377 loops (yellow; 29.3%) were formed between two CTCF sites (where each loop-anchor had one unique CTCF site within 10 kb) in at least one haplotype. For reference, Mustache called 18,068 total loops in GM12878 at 5 kb resolution, with 5,318 loops (29.4%) forming between two CTCF anchors (Roayaei Ardakany et al. 2020).

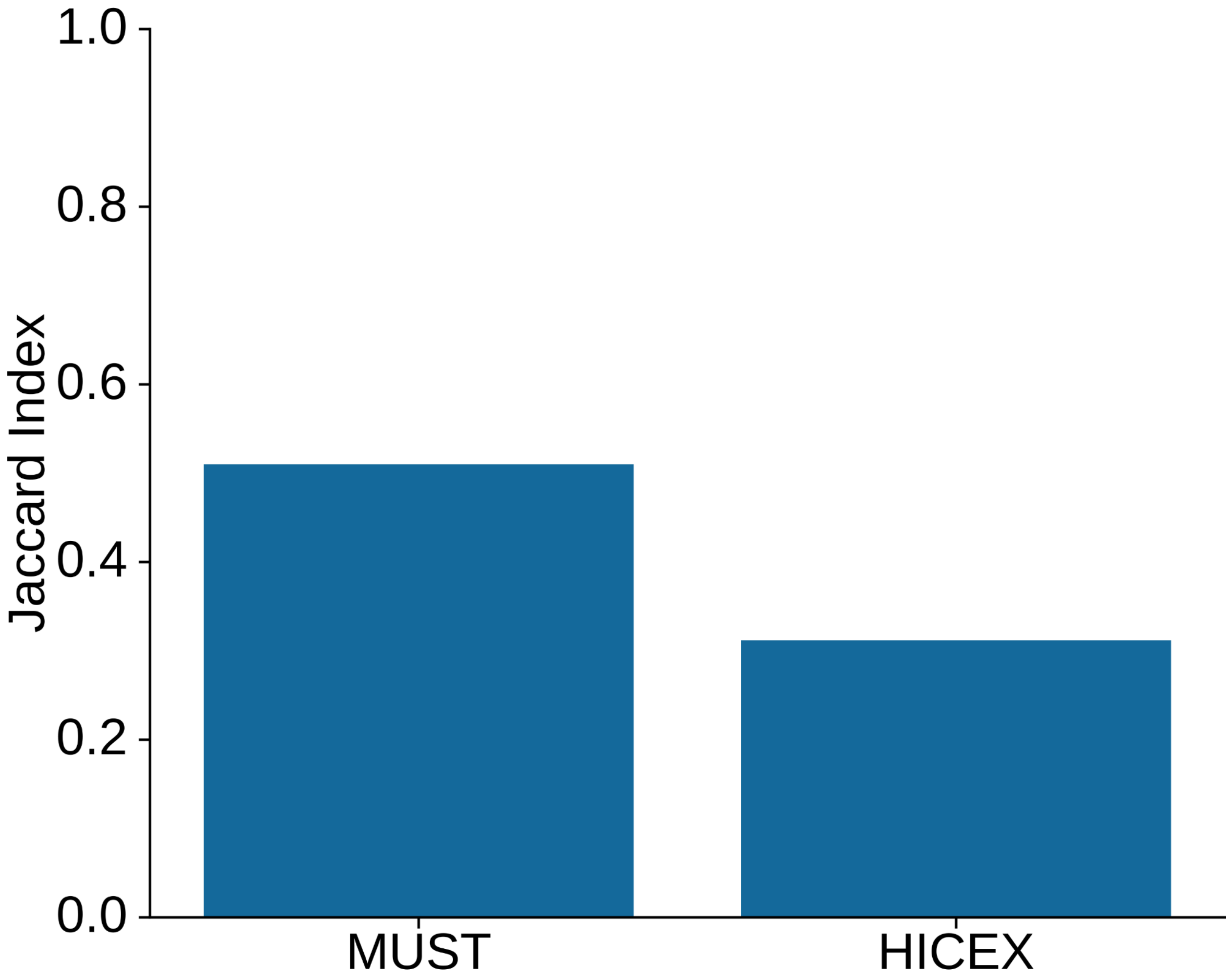


**Supplementary Figure 20. Average Jaccard index of loop calls between technical runs for Mustache (MUST) and HiCExplorer (HICEX) at 10 kb resolution.** Technical run IDs 1113258 and 1149781 correspond to sample GM19317.

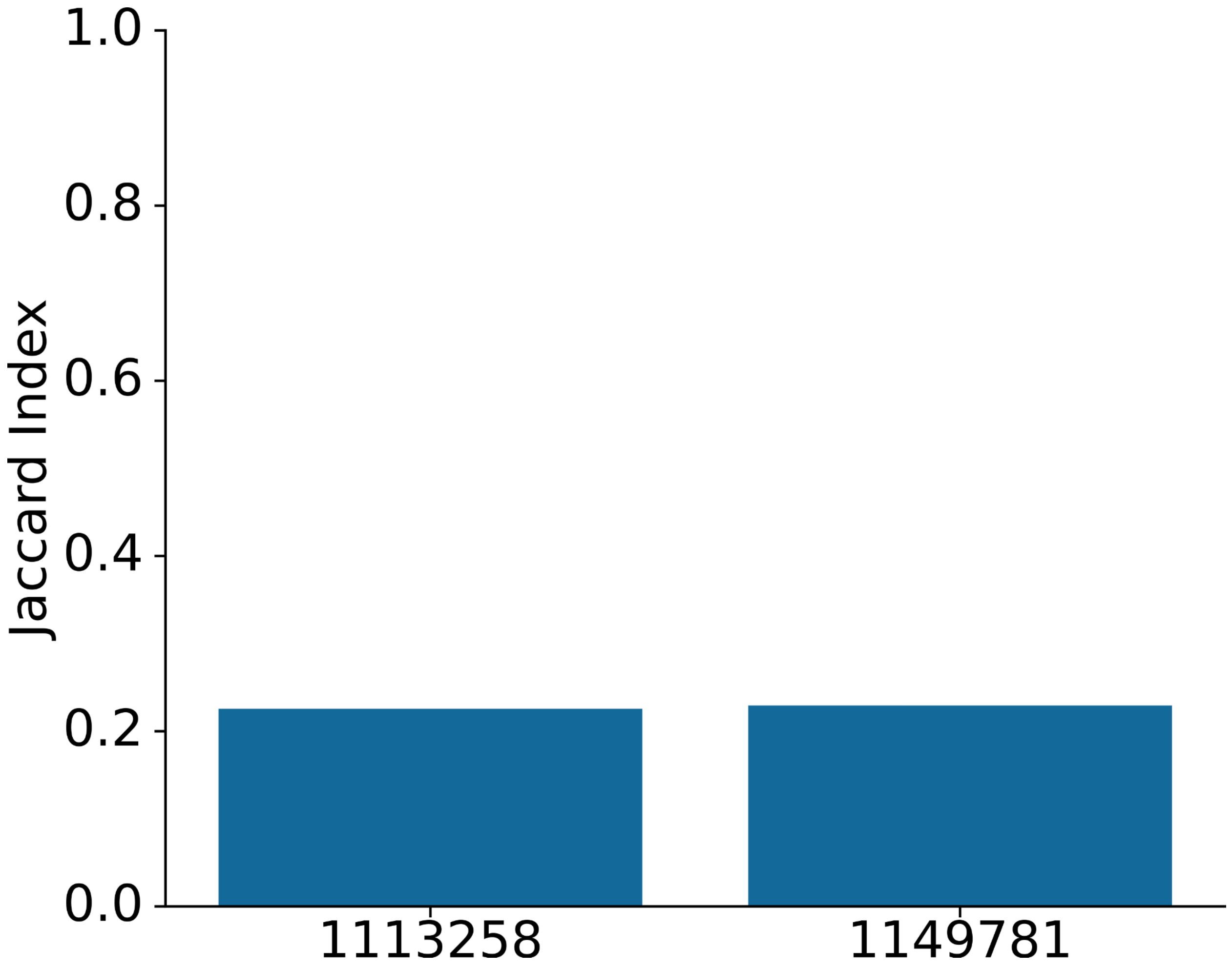

**Supplementary Figure 21. Average Jaccard index of loop calls between Mustache and HiCExplorer across two technical runs at 10 kb resolution.** Technical run IDs 1113258 and 1149781 correspond to sample GM19317.

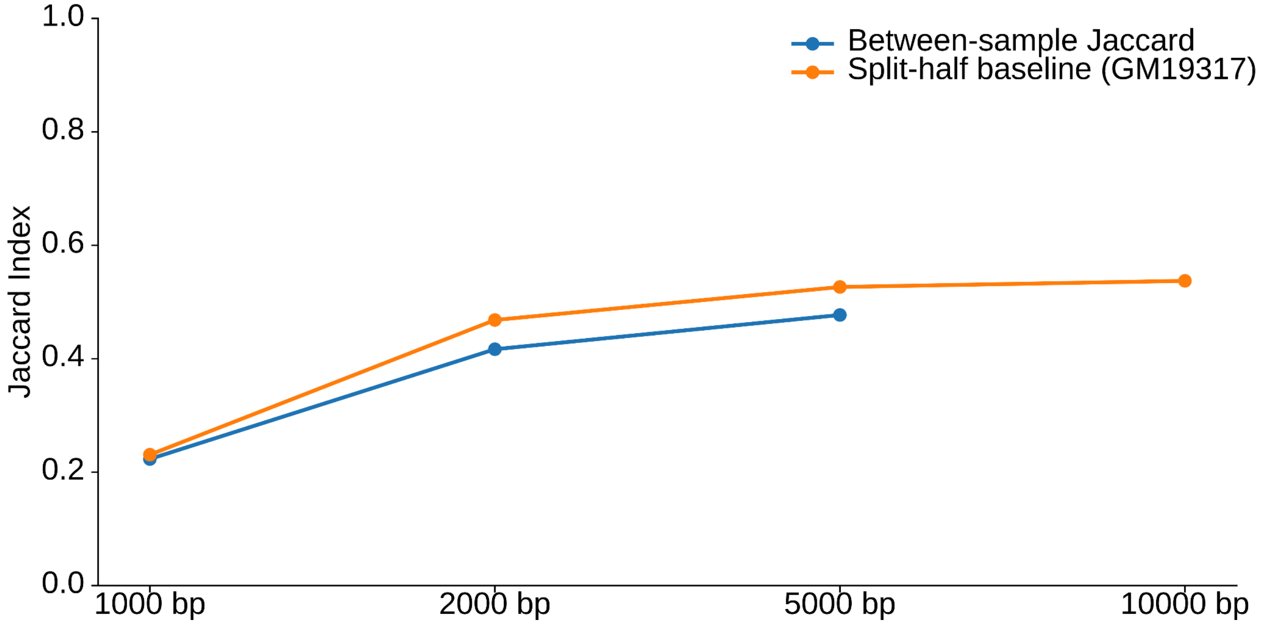


**Supplementary Figure 22. Average between-sample Jaccard index values at 1 kb, 2 kb, and 5 kb resolutions (blue) and maximum Jaccard index values expected due to sequencing depth for GM19317 at 1 kb, 2 kb, 5 kb, and 10 kb resolutions (orange).** The expected Jaccard index given perfect loop-callers with perfect data would approach 1.0, but the observed baseline Jaccard index doesn't substantially increase from 2 kb to 10 kb resolutions, despite the dataset having more than sufficient sequencing depth power at 10 kb resolution. Thus, the between-sample Jaccard indices approach the limit set by the calling range of Mustache rather than by sequencing depth.

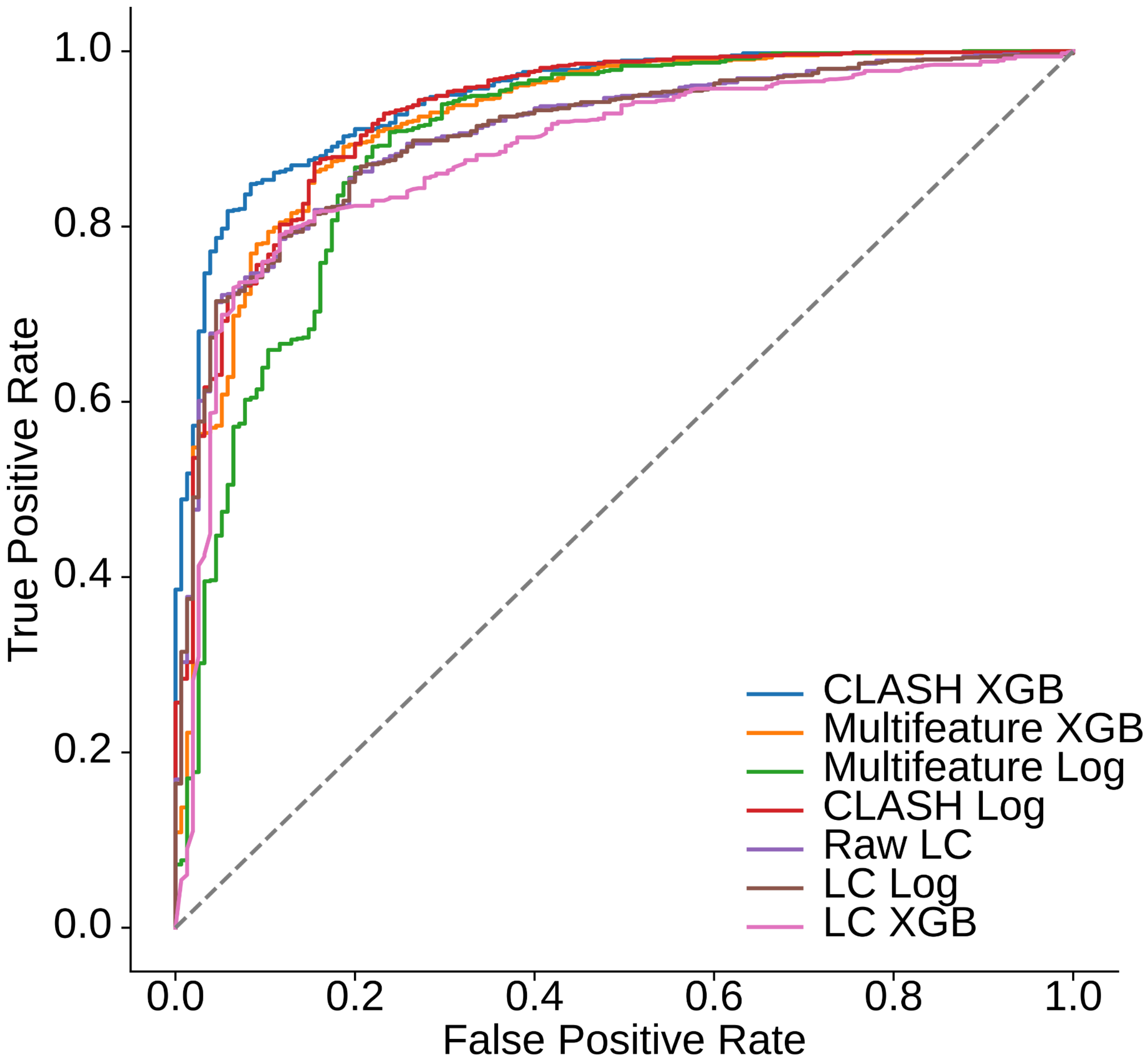


**Supplementary Figure 23. AUROC comparison of classifier methods and features, including CLASH XGBoost ("CLASH"; blue, AUROC = 0.945), Multifeature XGBoost (orange, AUROC = 0.921), Multifeature logistic regression (green, AUROC = 0.894), CLASH logistic regression (red, AUROC = 0.929), raw LC scores (purple, AUROC = 0.904), LC logistic regression ("LC model"; brown, AUROC = 0.904), and LC XGBoost (pink, AUROC = 0.882).** All models were trained using a 5-fold GroupKFold cross-validation framework, where the 5 samples at each locus were grouped, and displayed curves represent test ROCs on out-of-fold predictions. Raw LC evaluated LC scores directly as a continuous ranking score without model training.

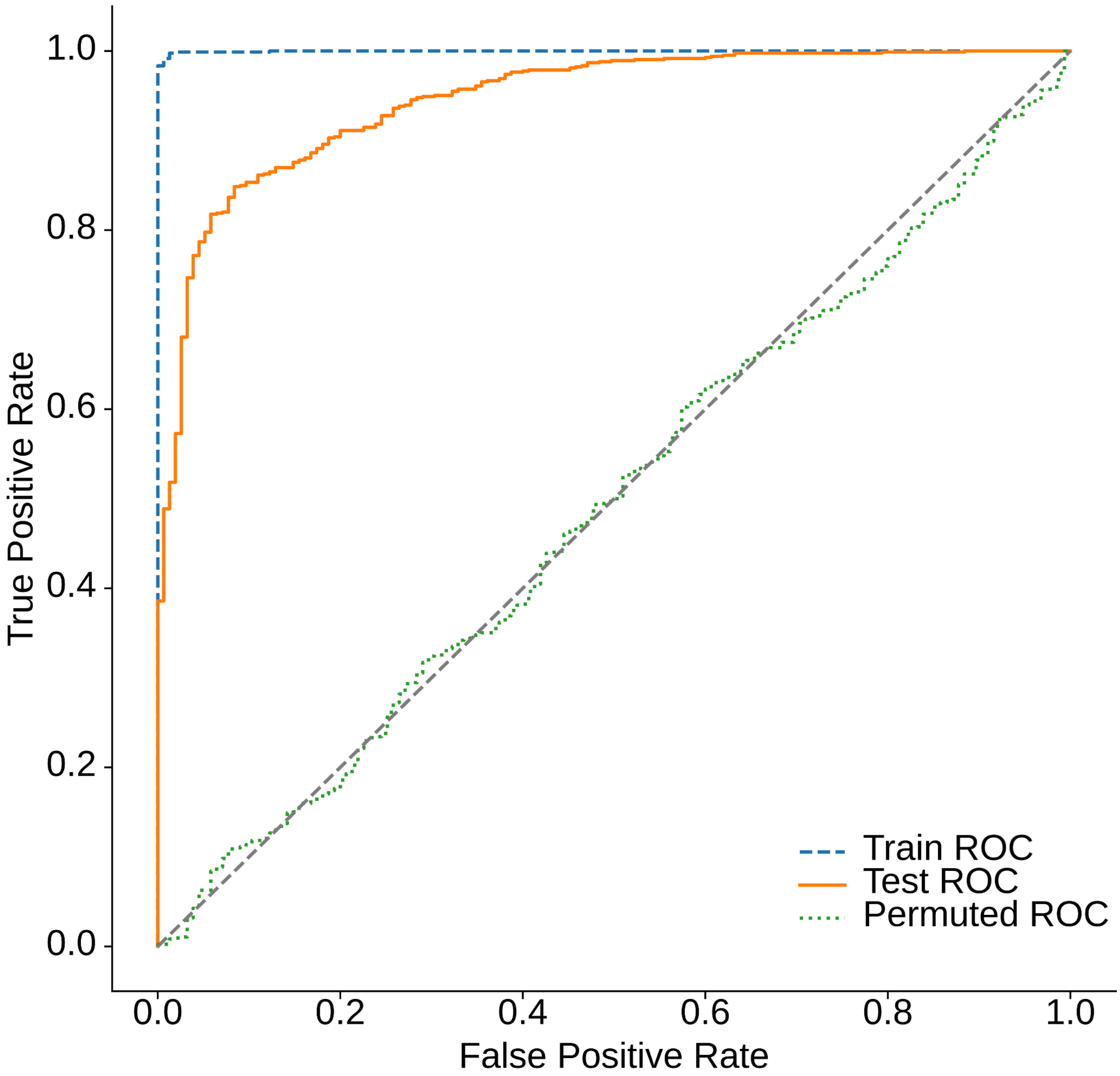


**Supplementary Figure 24: Cross-validated CLASH ROC curve analysis with permutation control.** ROC curves were generated comparing CLASH model performance on training data, held-out test data, and a permutation-based null model. The model was trained using a 5-fold GroupKFold cross-validation framework, where the 5 samples at each locus were grouped to prevent information leakage across folds. Training ROC (blue) was computed by fitting the model on the full dataset and evaluating predictions on the same data (AUROC = 1.0). Test ROC (orange) was computed using out-of-fold predictions, and all predictions were aggregated to compute a single ROC curve (AUROC = 0.945). Permuted ROC was generated by randomly shuffling class labels across all loci prior to cross-validation, and out-of-fold predictions were again used to compute the ROC curve (AUROC = 0.495). With perfect training AUROC, high test AUROC, and permuted ROC ≈ 0.5, the model has the capacity to fit the data, generalizes well to unseen data, and learns real signals.

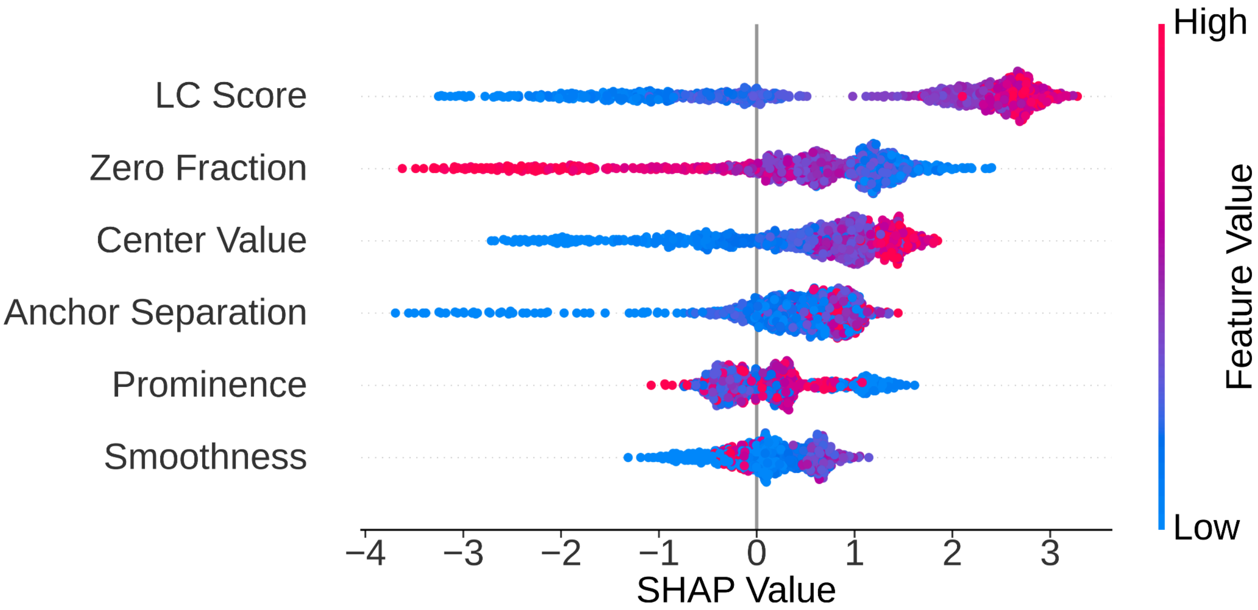


**Supplementary Figure 25. SHAP Value Decomposition of feature effect on CLASH output.** Although LC scores have the largest single impact, each of the other features adds meaningful discrimination power allowing the full model to achieve the highest AUROC.

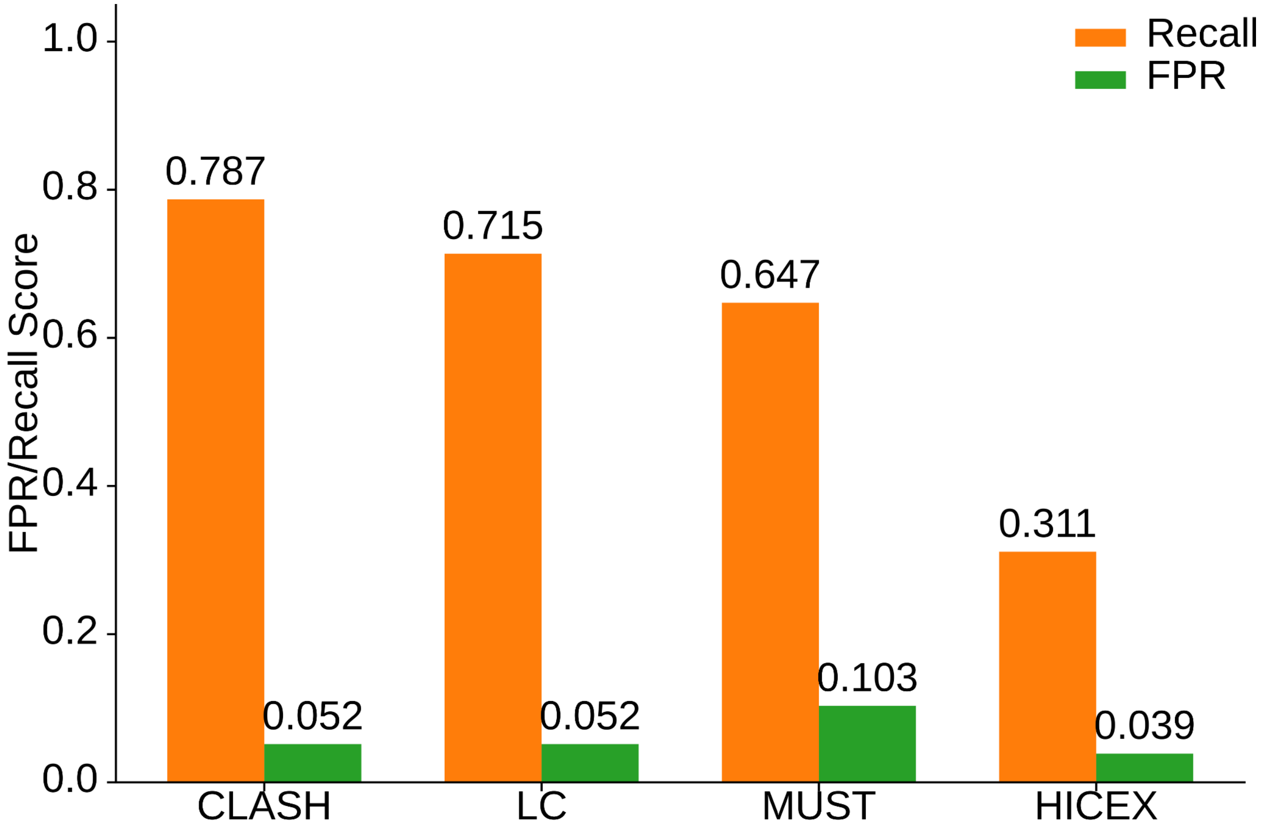


**Supplementary Figure 26. Comparison of CLASH loop call recall on the held-out training set against LC model, binary Mustache (MUST) calls, and binary HiCExplorer (HICEX) calls.** At a matched FPR ≈ 0.05, CLASH has a higher recall of +0.072 than LC model. For the binary calls, there was only a single FPR as calls were already made.

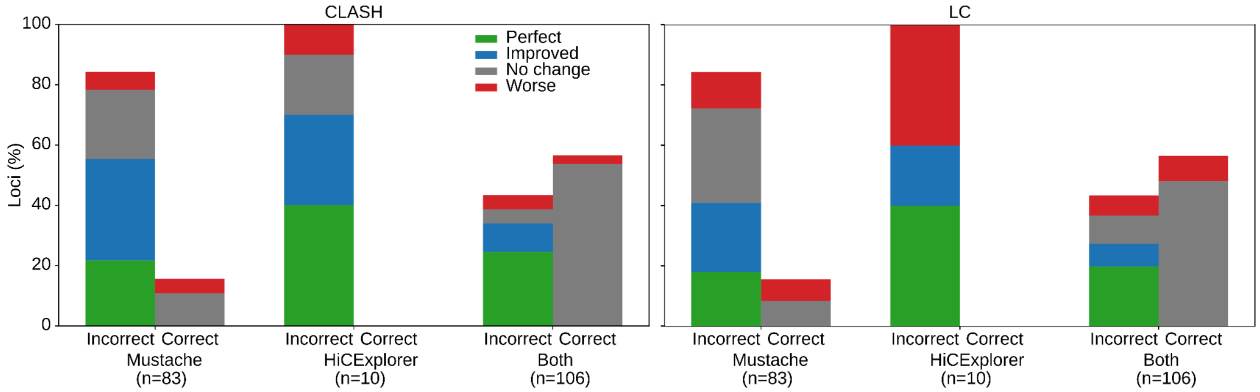


**Supplementary Figure 27. Stratification of CLASH harmonization effects (perfect improvement: green, improvement: blue, no change: grey, worse: red)  on Mustache discovered, HiCExplorer discovered, or both Mustache and HiCExplorer discovered loops,**

**separated by loci that initial callers called correctly vs incorrectly.** Out of the 199 loci, Mustache and HiCExplorer collectively incorrectly call loops in 126 of these loci. CLASH harmonization (left), using out-of-fold predictions and the decision boundary that maximizes Youden's J, improves loop calls in 89 of these loci, while worsening calls in 18 loci. Comparatively, using LC model as the classifier (right), also using the decision boundary that maximizes Youden's J, improves 69 loci while worsening 36 loci worse.

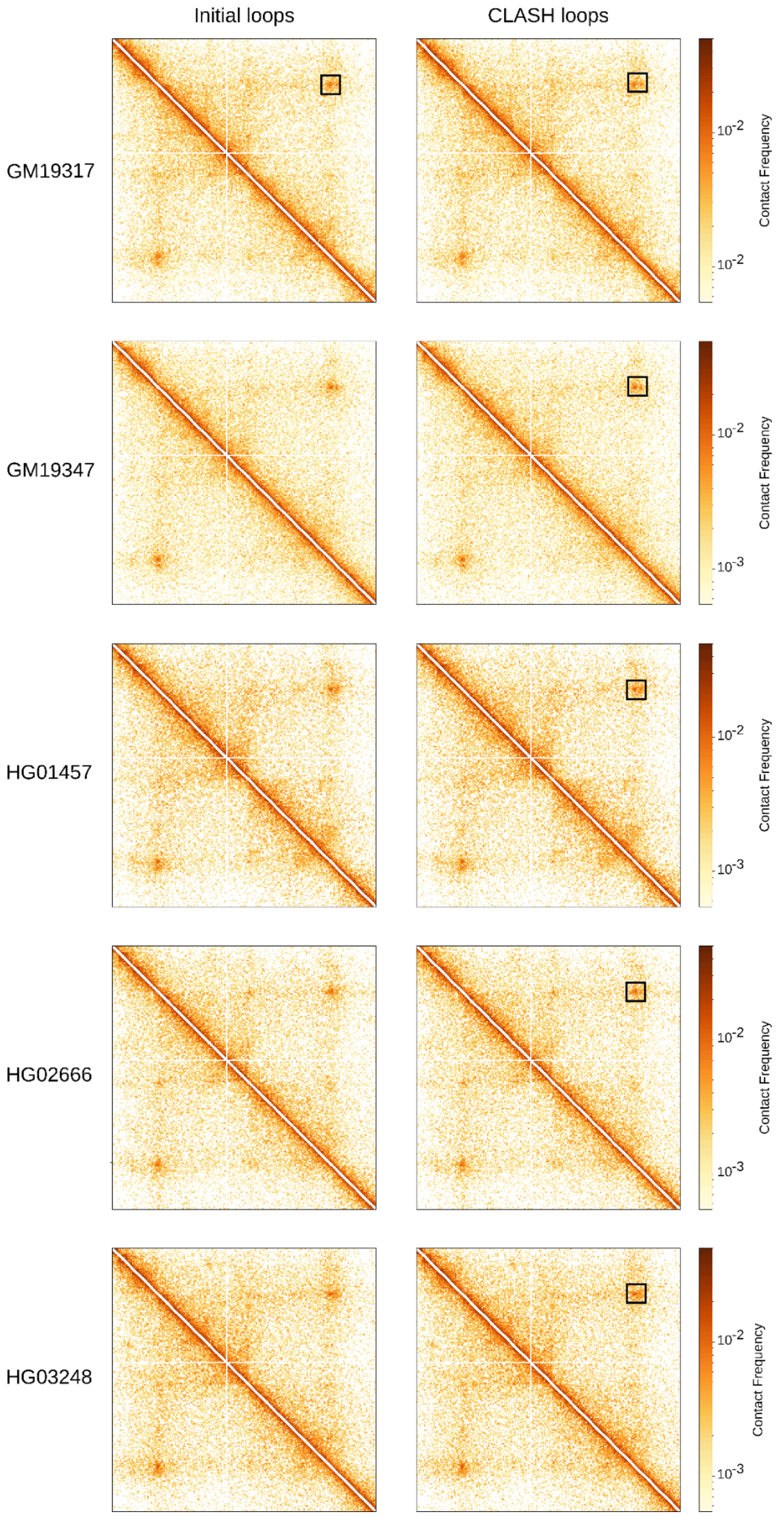
Initial loops
CLASH loops
GM19317
GM19347
HG01457
HG02666
HG03248
Contact Frequency
10-2
10-3

**Supplementary Figure 28. Example chromatin loop before and after CLASH harmonization.**

The representative locus (chr5:14728000–15000000) illustrates loop-calling inconsistency across samples prior to CLASH harmonization (left), where Mustache and HiCExplorer fail to identify the same loop across individuals. Collectively, they call the loop only in sample GM19317 (black rectangle) despite clear contact enrichment across all samples. After CLASH harmonization (right), the same locus is consistently labelled as a loop across all five samples.

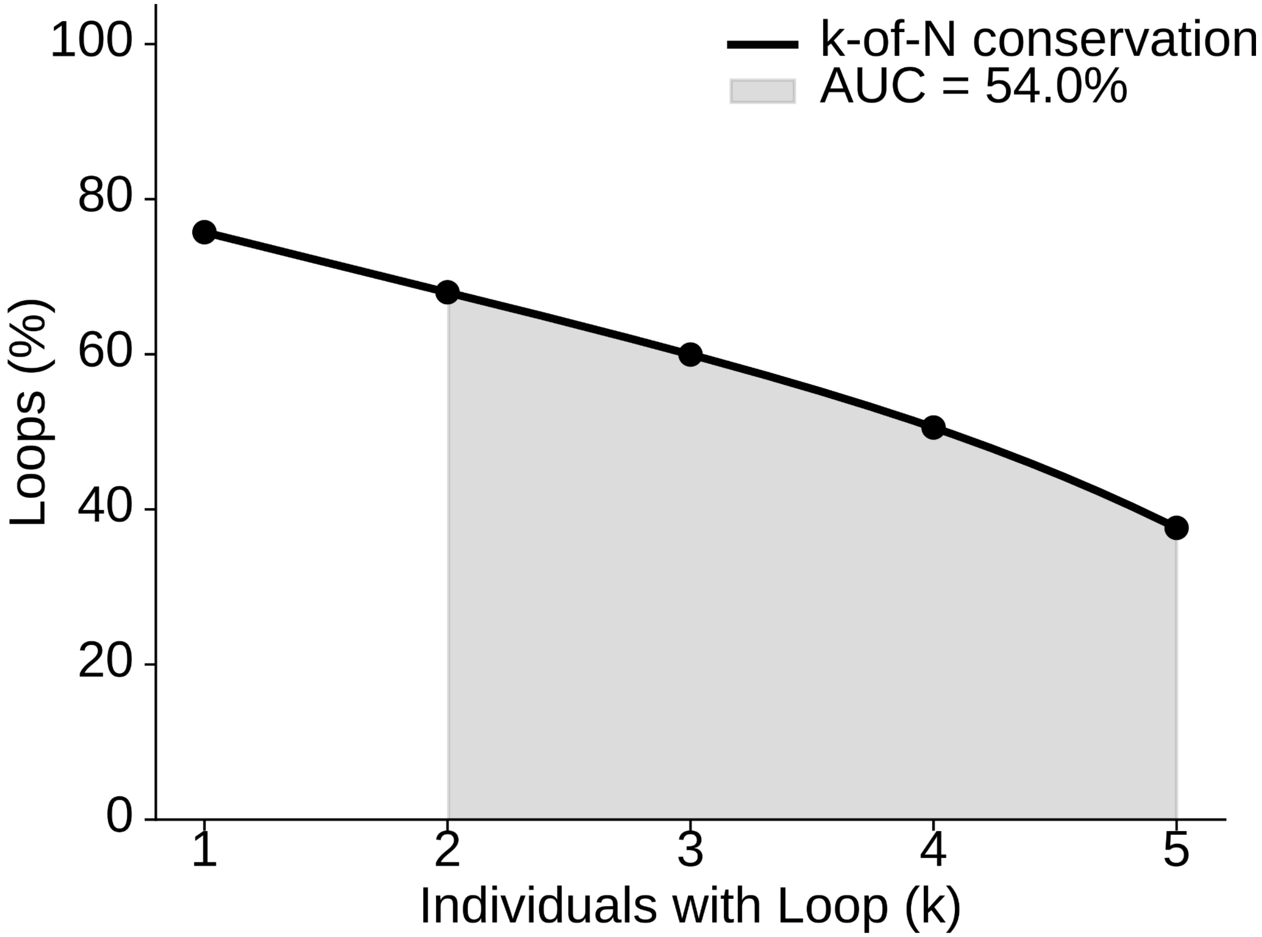


**Supplementary Figure 29. Smoothed k-of-N conservation curve of binarized CLASH-scored loops.** The area under the curve (AUC = 0.54) of the portion of the graph with k ≥ 2 corresponds to the conditional conservation probability. Total n = 6,377 loci.

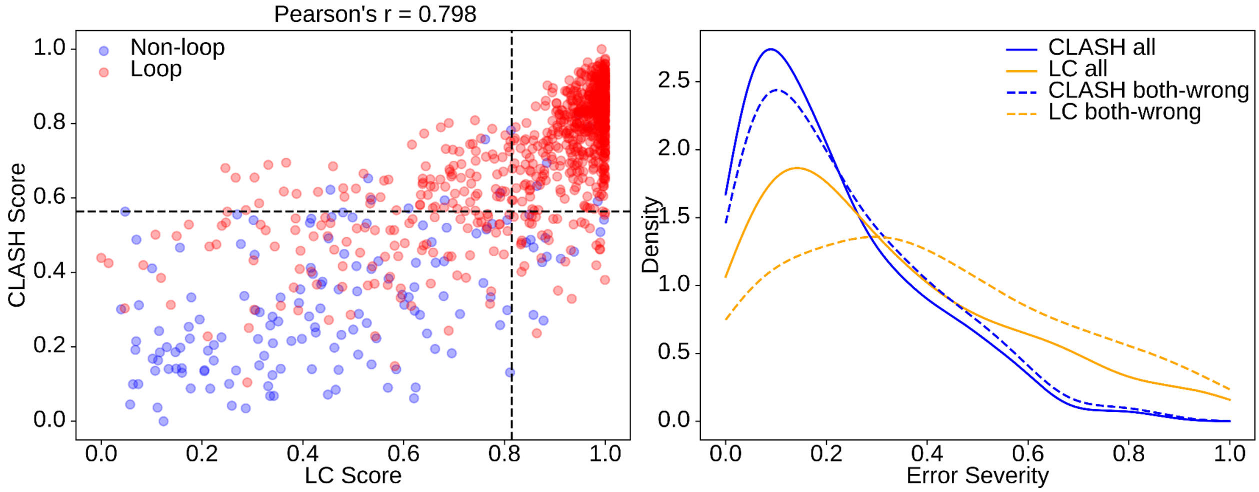


**Supplementary Figure 30: CLASH misclassifications have continuous scores closer to the decision boundary than LC misclassifications within each method's score space. Left:** Scatter plot showing CLASH and LC model classification (out-of-fold) predictions, and true labels (red = loop, blue = no loop) of each putative loop in the training set, as well as the decision boundary for both CLASH scores and LC model scores – both calculated as the min-maxed logits of each classification – as determined by the operating point that maximizes Youden's J (CLASH = 0.56, LC = 0.81). CLASH misclassifies 141 of the 1000 loops while LC model misclassifies 197, sharing 99 misclassifications in common. The correlation between CLASH scores and LC model scores is 0.80 ($p = 1.71 \times 10^{-221}$). **Right:** Distribution of error severity for CLASH and LC model misclassifications, defined as the normalized distance from the decision boundary relative to the extent of the incorrect region. CLASH score errors are significantly smaller than LC model score errors across all misclassified loci (mean = 0.21 vs 0.31; median = 0.16 vs 0.24; Mann–Whitney U test $p = 8.1 \times 10^{-5}$), and this difference remains when restricting to loci misclassified by both methods (Wilcoxon signed-rank test $p = 8.60 \times 10^{-7}$). These results indicate that CLASH not only reduces the number of misclassifications relative to LC, but also produces scores that are better calibrated with respect to the decision boundary, with errors occurring closer to the classification threshold.

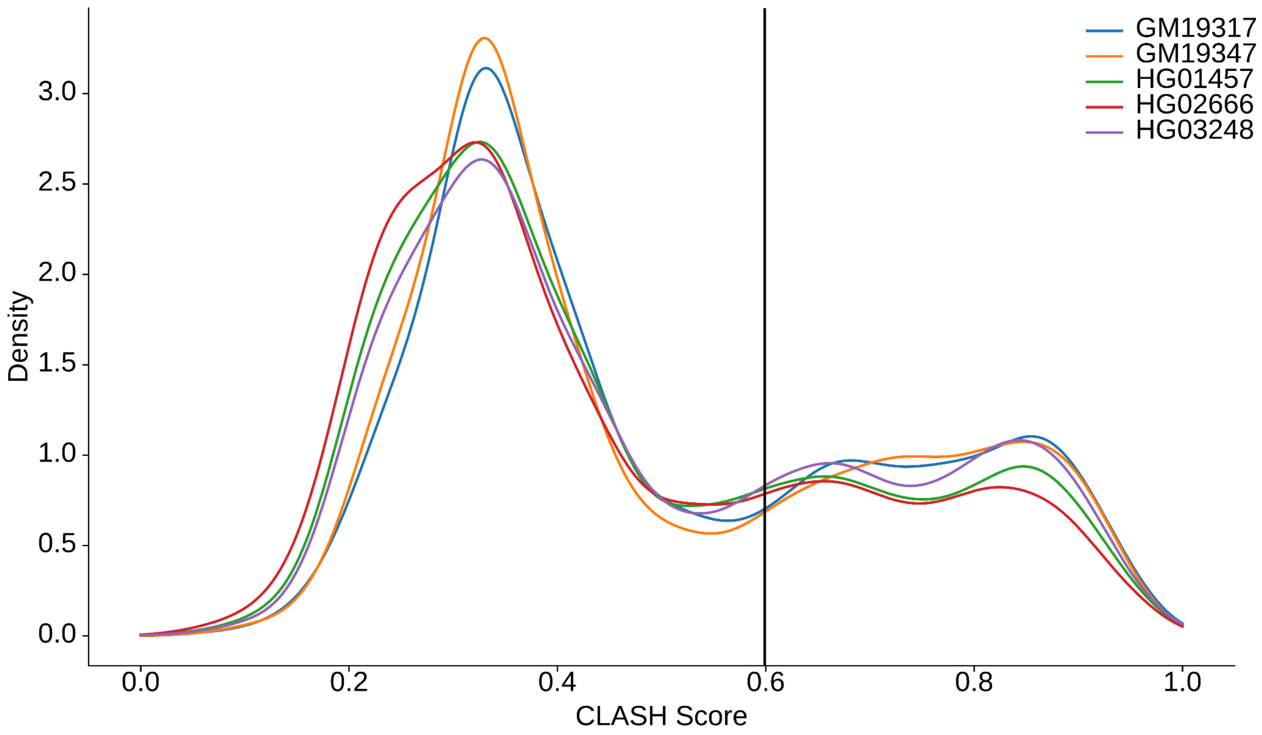


**Supplementary Figure 31. CLASH scores display minimal sample-dependent biases.** Distribution of CLASH scores on the full dataset, stratified by sample. Sample distributions show minimal differences (mean scores range from 0.45 - 0.50, median scores range from 0.37 - 041, standard deviations range from 0.22-0.23, and the maximum absolute standardized deviation from the global mean = 0.13). The decision boundary is shown as a black line.

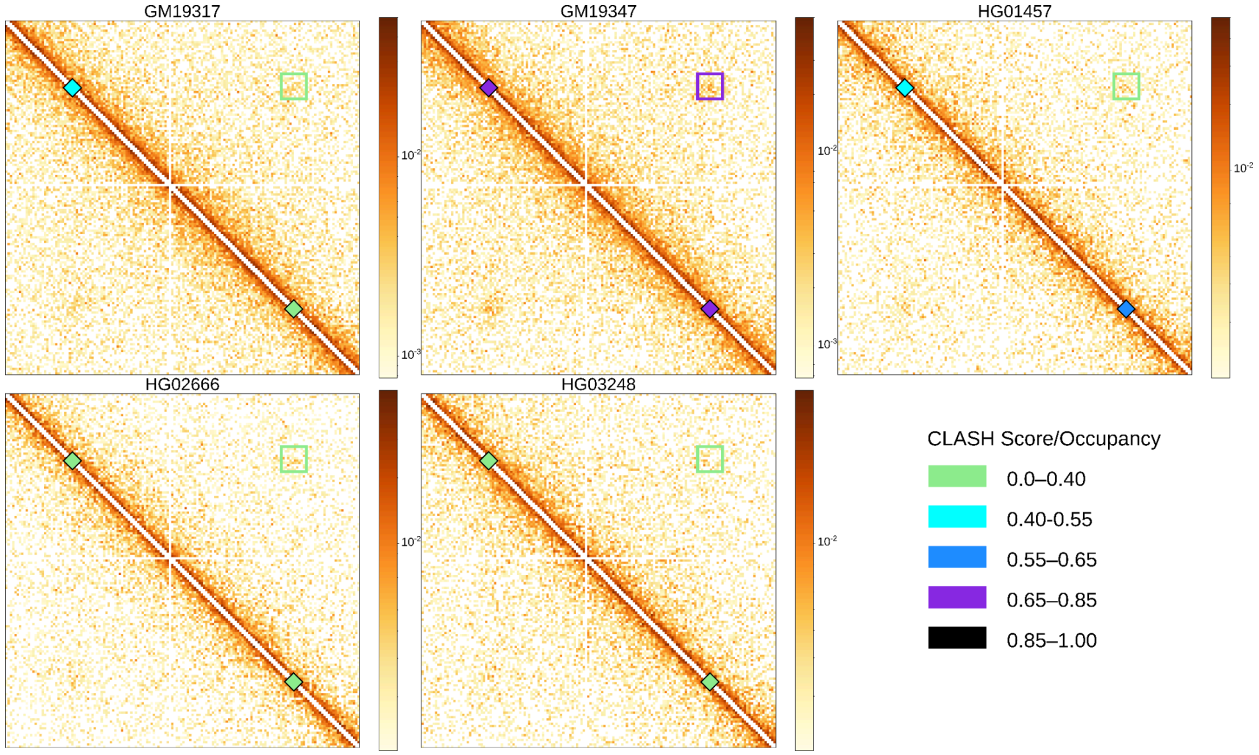


**Supplementary Figure 32. Example locus (chr1:115216363-115527735) across all five samples, illustrating a positive relationship between CTCF occupancy and loop scores.** Both CTCF occupancy and CLASH loop scores are annotated for each sample with light green = 0-0.40, light blue = 0.40-0.55, blue = 0.55-0.65, purple = 0.65-0.85, and black = 0.85-1.00. The loop score is annotated by the color of the rectangle surrounding the loop, and CTCF occupancy of each CTCF site is annotated as the color of the rectangles along the diagonal. GM19347 has higher CTCF occupancy in both loop-associated CTCF sites than the other samples, corresponding to an increase in loop score.

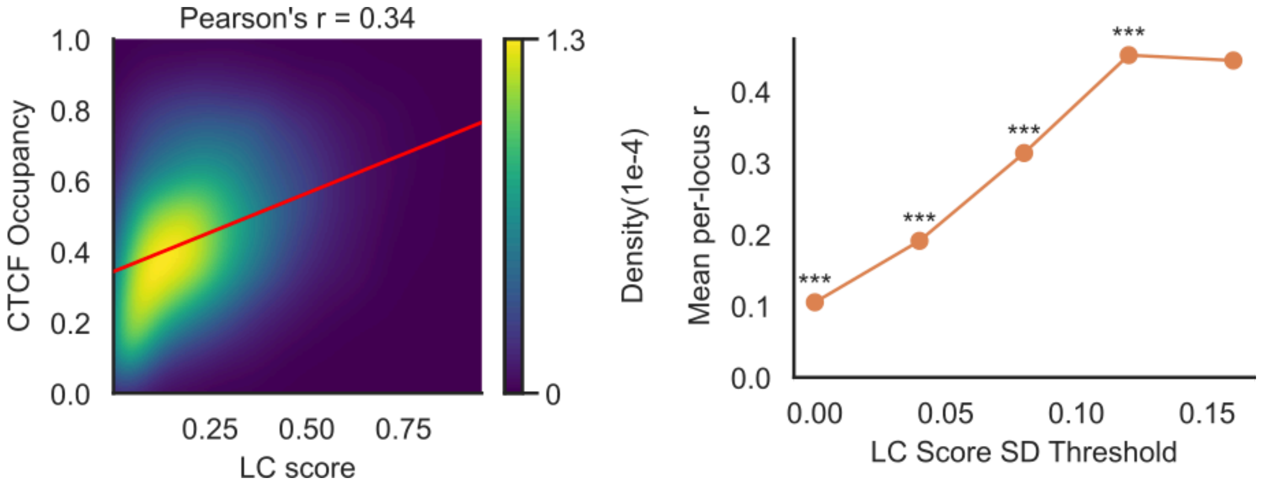


**Supplementary Figure 33**. **Global (left) and per locus correlations of LC model scores with CTCF occupancy are similar to CLASH score correlations with CTCF occupancy.** Globally, LC model scores have a Pearson's correlation of r = 0.342 (n = 21,050, $p < 2.2 \times 10^{-308}$). Per locus correlations (n = 3,144) increase as loop variability increases. Significance was assessed using a one-sided one-sample t-test against zero.

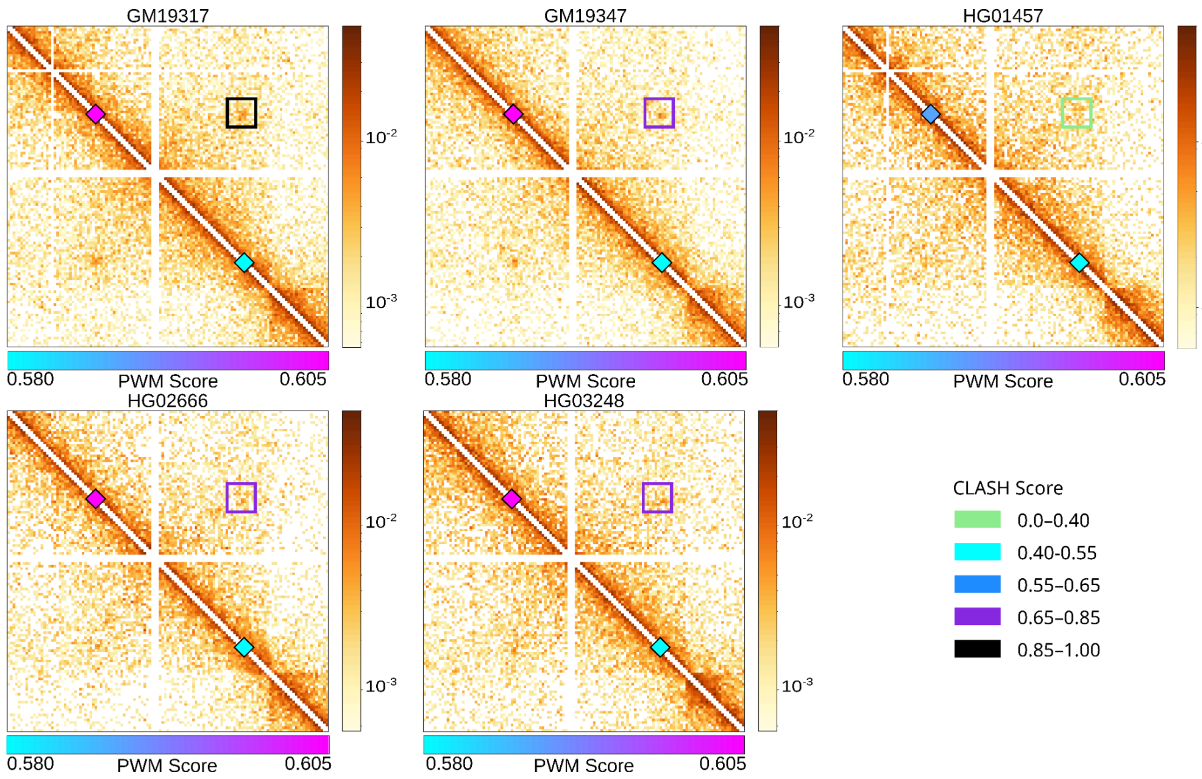


**Supplementary Figure 34. Representative locus (chr2:134,714,609–134,963,448) illustrating the relationship between CTCF PWM scores and loop strength across samples.** The same

CLASH score coloring annotation as Supplementary Figure 32 is used. In HG01457, the upstream loop-associated CTCF site contains a heterozygous SNP at the 14th base position (G → A), resulting in a reduced PWM score (darker blue). This nucleotide change corresponds to both the lowest CTCF occupancy at that site and the weakest loop formation across samples. For clarity, only relevant CTCF sites are shown.

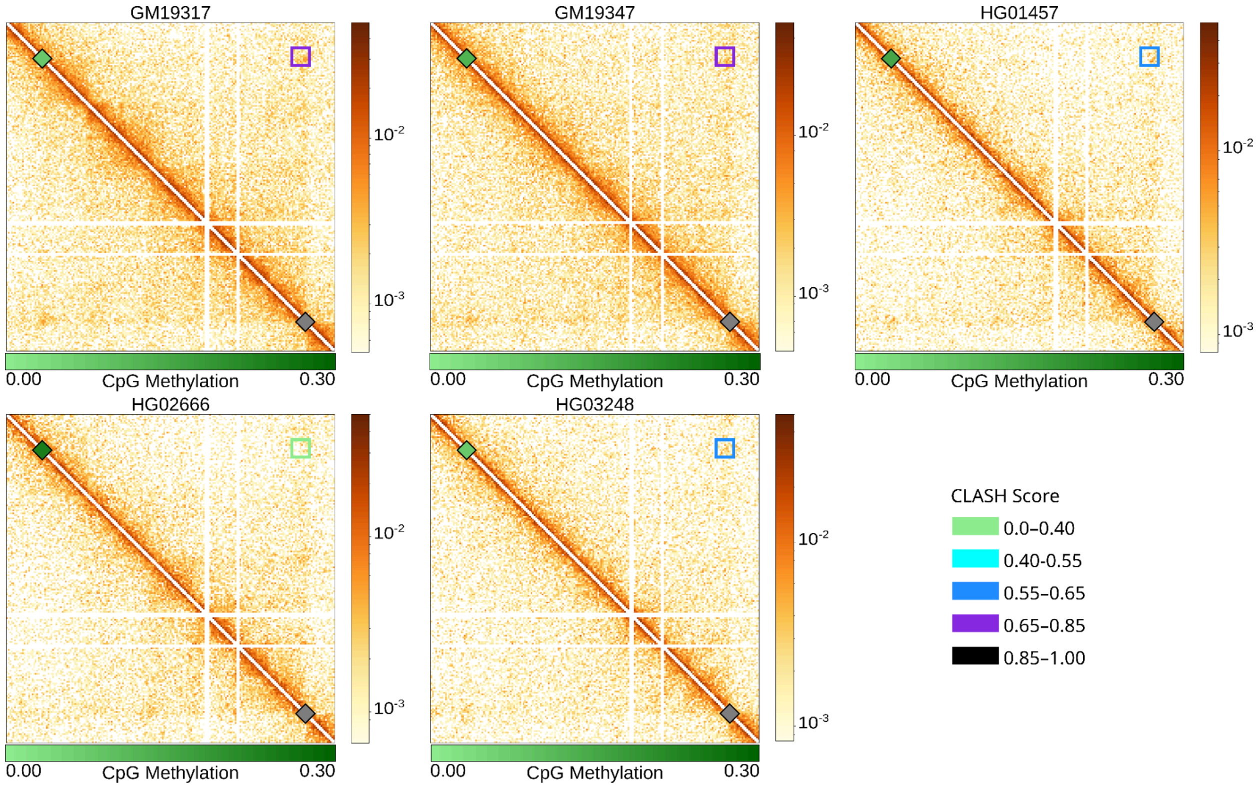


**Supplementary Figure 35. Representative locus (chr4:85347667–85810593) illustrating the expected relationship between m$^5$C methylation and loop strength across samples.** The same CLASH score coloring annotation as Supplementary Figure 32 is used. In HG02666, the upstream loop-associated CTCF site is significantly more methylated (darker green) than the other samples. This corresponds to both the lowest CTCF occupancy at that site and the weakest loop formation across samples. For clarity, only relevant CTCF sites are shown.

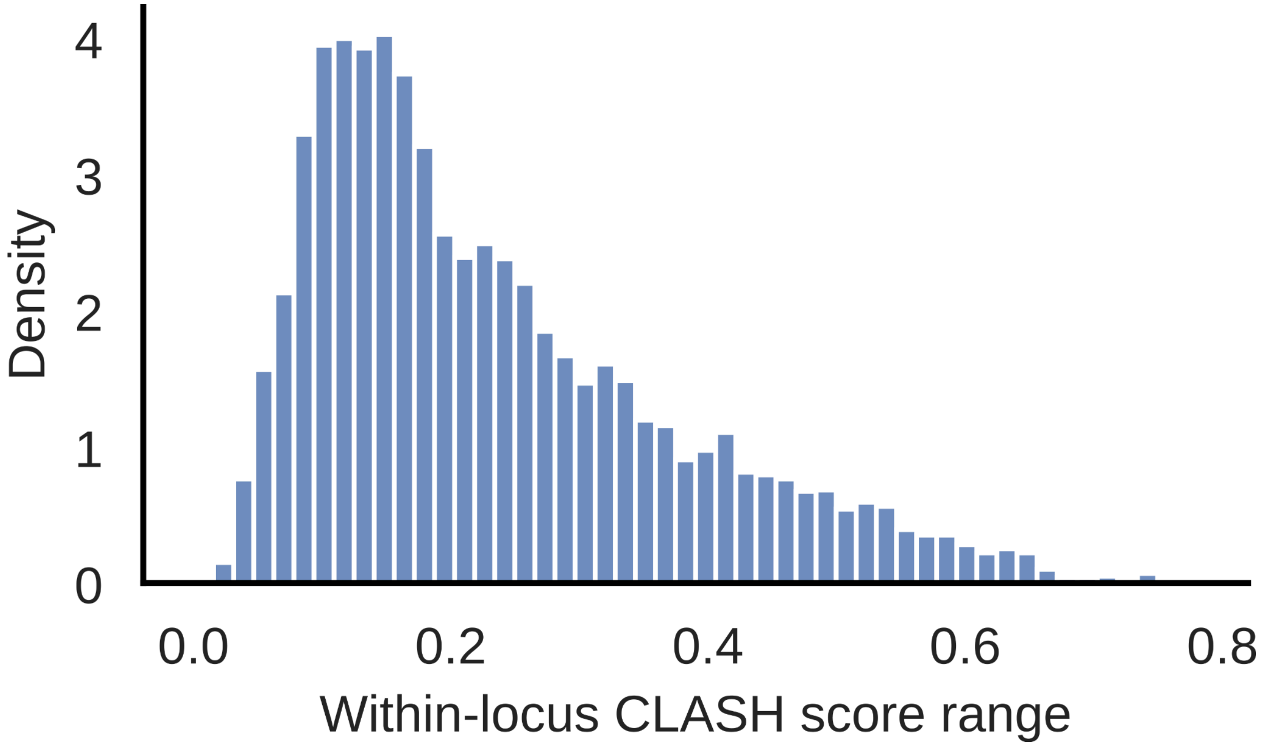


**Supplementary Figure 36. Distribution of within-locus CLASH loop score ranges (n = 6,377).**

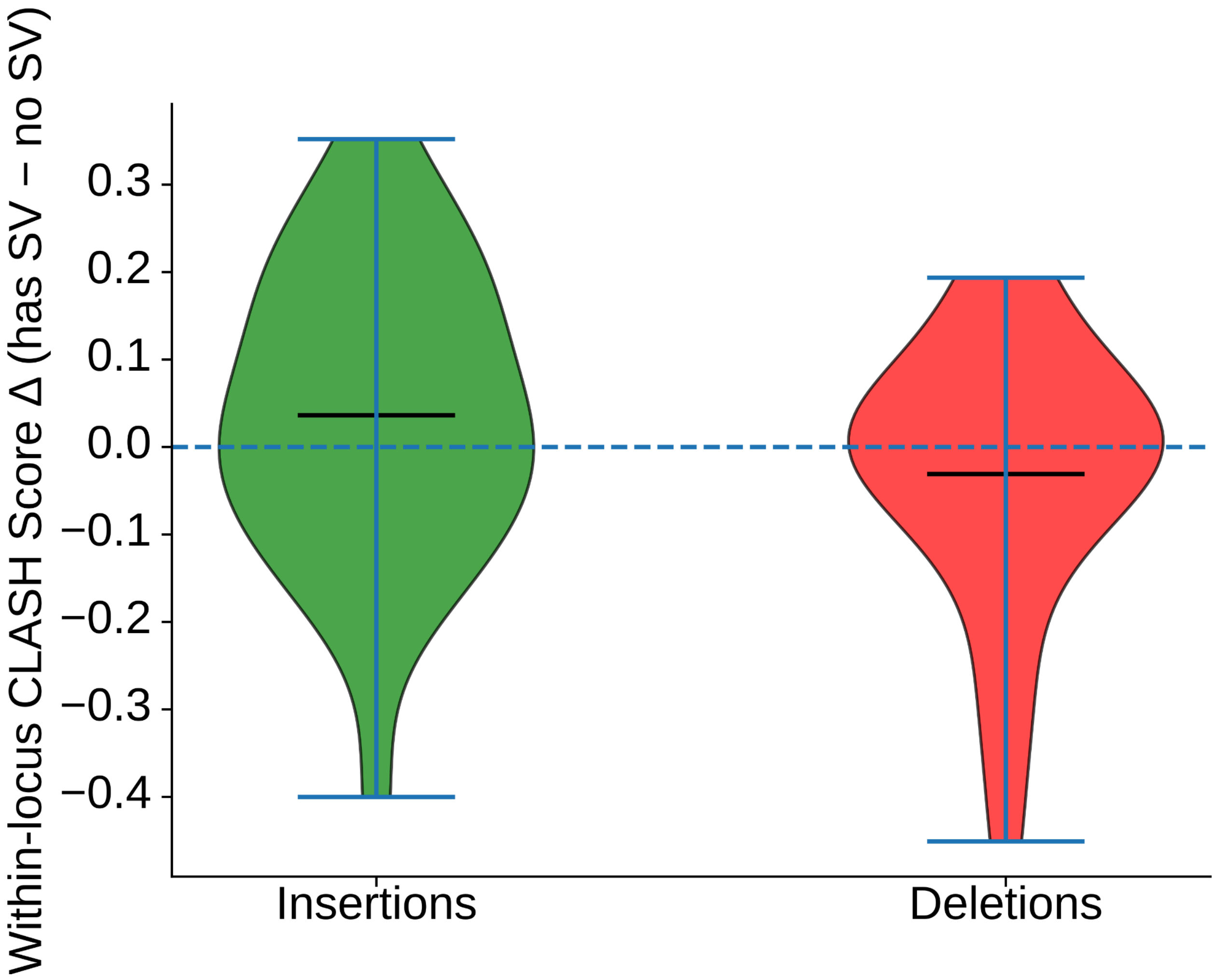


**Supplementary Figure 37. Distribution of CLASH score differences for loci containing structural-variant insertions (mean Δ=0.0363, n = 27, left) or deletions (mean Δ =-0.0308, n = 23, right), compared with samples lacking the variant at the same locus, indicating a minimal global effect of structural variation on loop strength.**

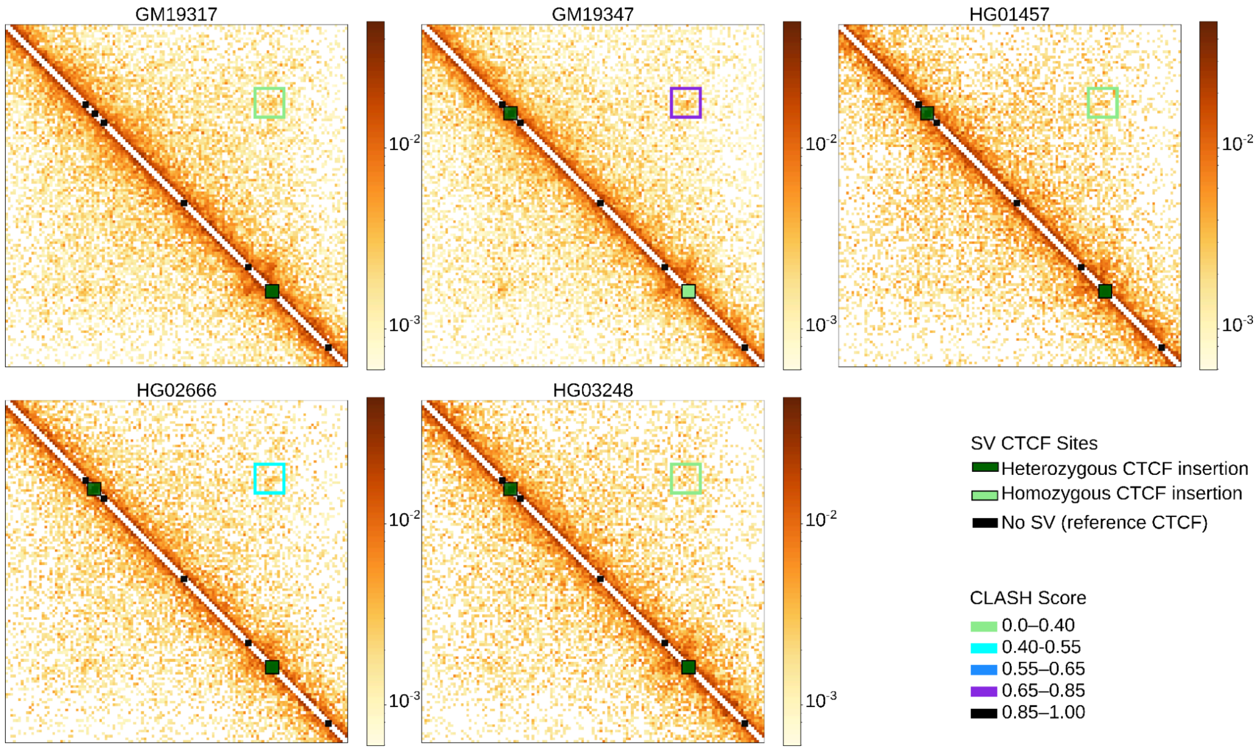


**Supplementary Figure 38. Example locus (chr2:119456242-119717829) showing a structural-variant insertion that introduces a homozygous CTCF site (light green) in GM19347, in contrast to the other samples with heterozygous insertions (dark green), leading to increased loop strength in GM19347.** Reference CTCF sites are shown in black, and the same CLASH score coloring annotation as Supplementary Figure 32 is used.

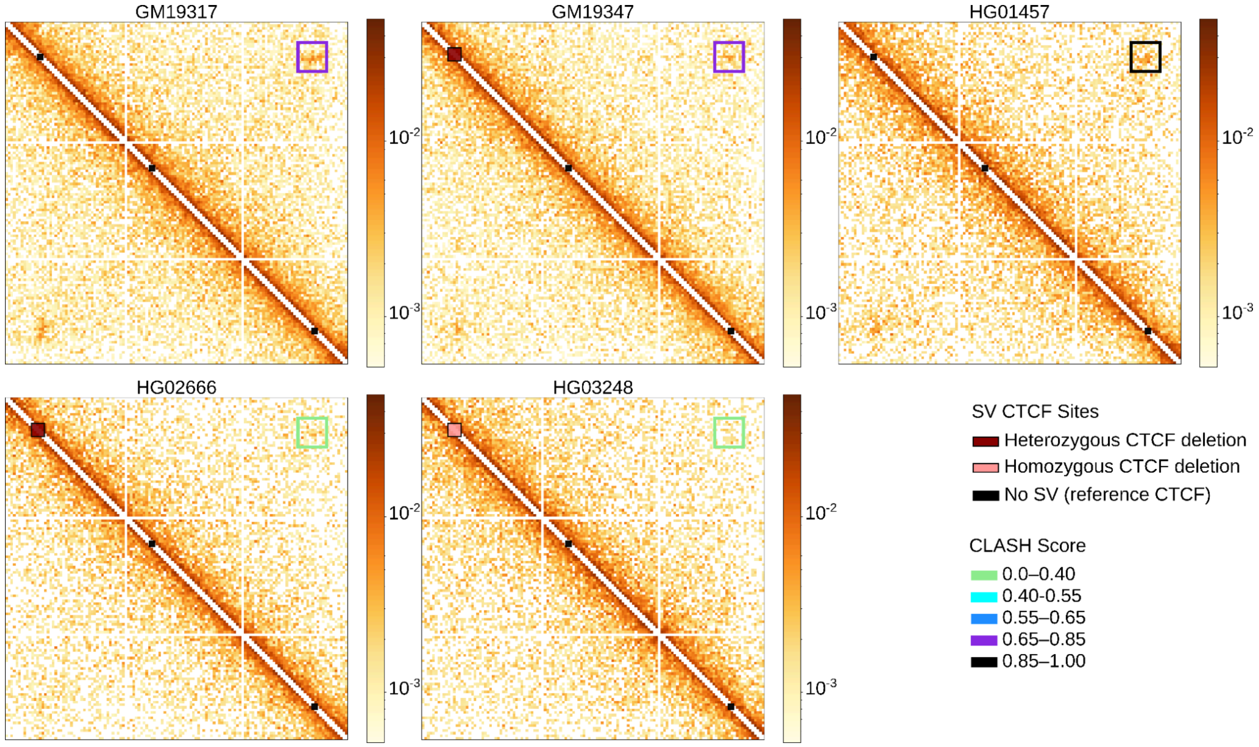


**Supplementary Figure 39. Example locus (chr18:53883045-54145059) showing a structural-variant deletion that removes a CTCF site, present as heterozygous deletions (dark red) in GM19347 and HG02666 and as a homozygous deletion (light red) in HG03248.** Reference CTCF sites are shown in black. Samples without any deletions exhibit strong loop formation, samples with heterozygous deletions show variable loop strength (as expected given non-haplotype-resolved Hi-C maps), and homozygous deletions exhibit minimal loop formation. The same CLASH score coloring annotation as Supplementary Figure 32 is used.

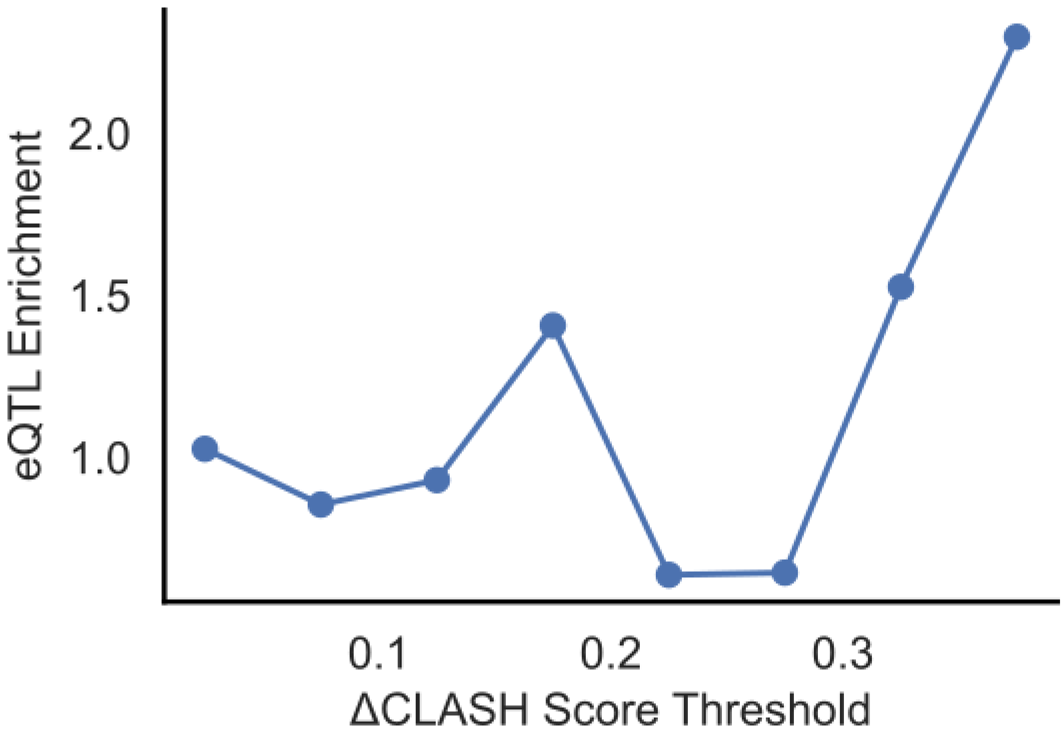


**Supplementary Figure 40. Enrichment of eQTLs among loop-altering SNPs (putative iQTLs; n = 555 total SNP-containing CTCF loci) at ΔCLASH score thresholds from 0.025 to 0.375.** Although this analysis lacks statistical power due to small sample size, the observed trend is consistent with expectations.

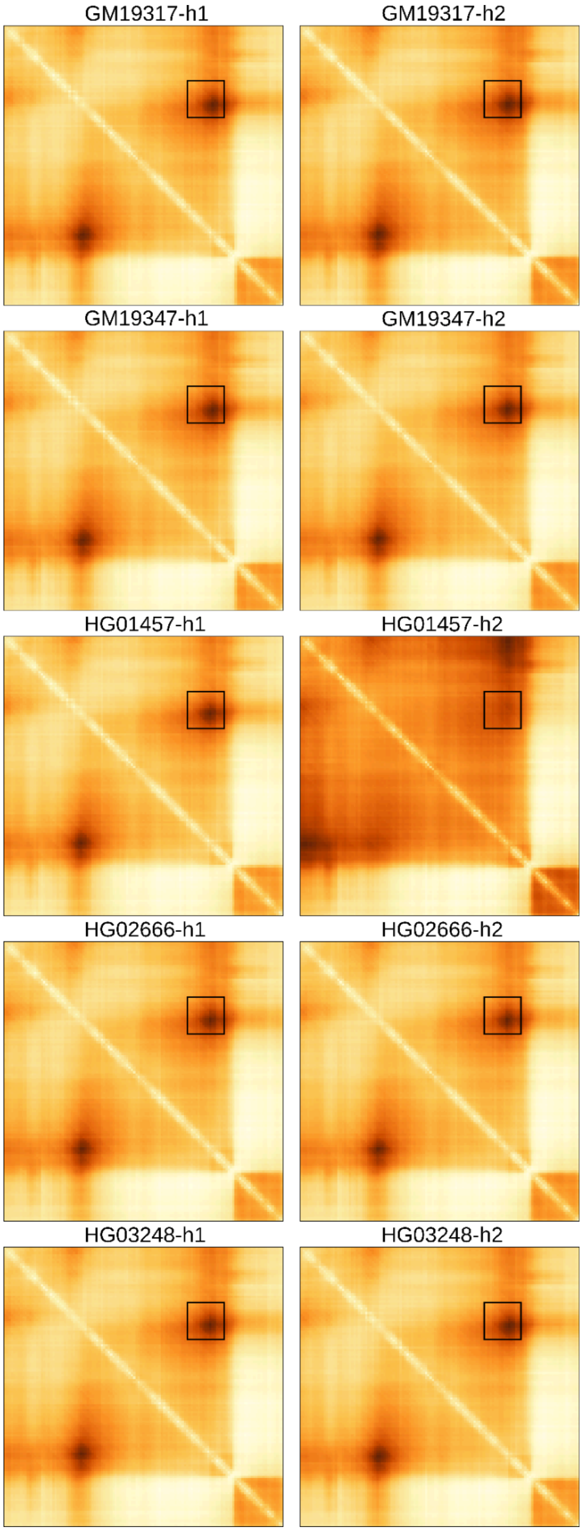


**Supplementary Figure 41. Accurate haplotype-specific loop prediction by AlphaGenome, matching our observation.** The haplotype-specific loop prediction for chr2:134,714,609-134,963,448 with observed contacts shown in Supplementary Figure 34**.** The predictions match the observed loop formation across the same samples, accurately predicting homozygous loops in all of the samples except HG01457 which contains a SNP in the corresponding CTCF binding site. The area of interest is highlighted with a black rectangle.

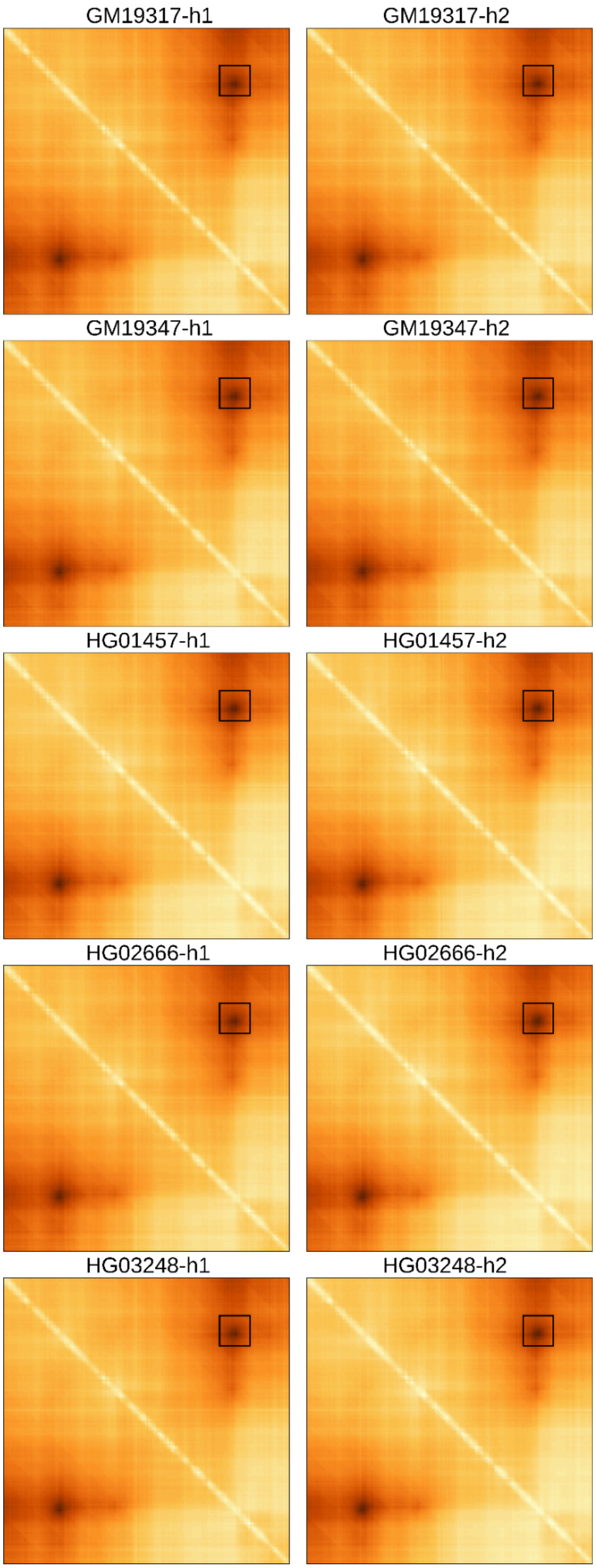
GM19317-h1
GM19317-h2
GM19347-h1
GM19347-h2
HG01457-h1
HG01457-h2
HG02666-h1
HG02666-h2
HG03248-h1
HG03248-h2

**Supplementary Figure 42. Misprediction of a haplotype-specific loop by AlphaGenome, chr1:115216363-115527735.** The region is the same locus as shown in Supplementary Figure 32. A homozygous loop (black rectangle) is predicted to form across all 5 samples, but these predictions do not match the observed loop formation variation, indicating that the model may benefit from incorporating CTCF occupancy signal into its predictions.

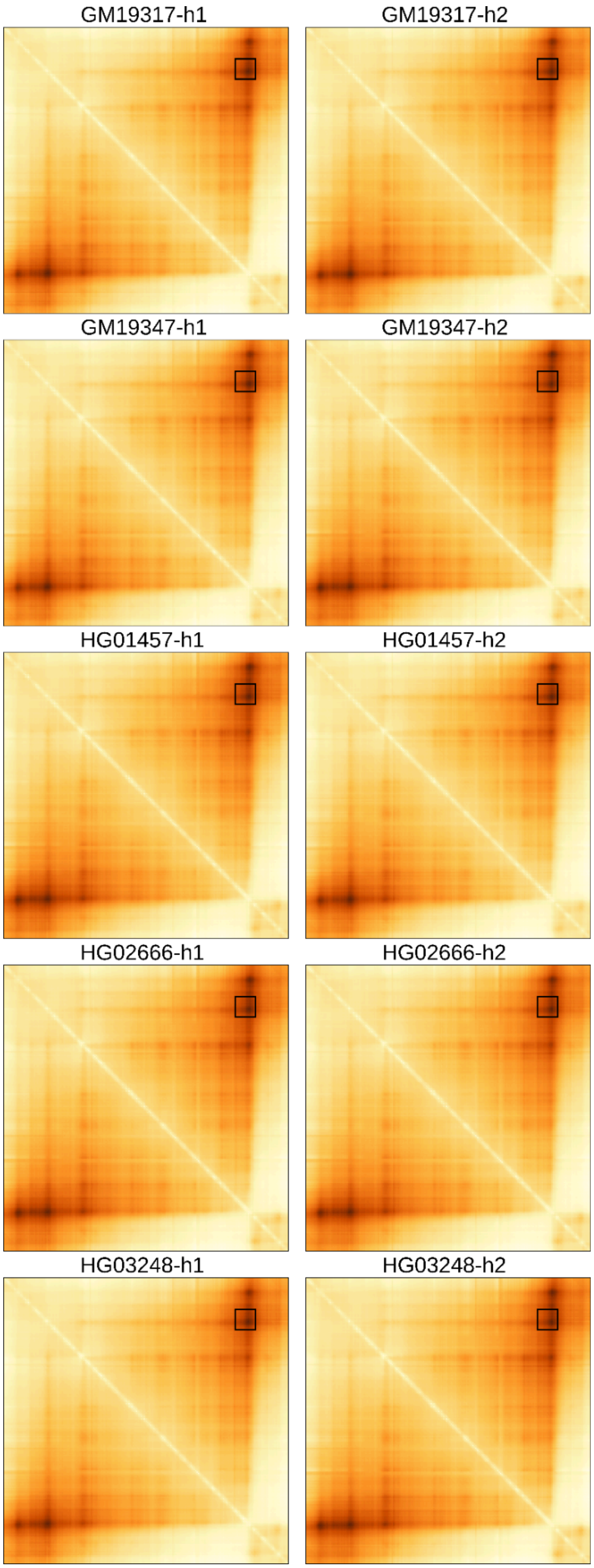
GM19317-h1
GM19317-h2
GM19347-h1
GM19347-h2
HG01457-h1
HG01457-h2
HG02666-h1
HG02666-h2
HG03248-h1
HG03248-h2

**Supplementary Figure 43. Misprediction of a haplotype-specific loop by AlphaGenome due to methylation, chr4:85347667–85810593.** This is the same locus as shown in Supplementary Figure 35. A homozygous loop (black rectangle) is predicted to form across all 5 samples, but these predictions do not match the observed loop formation variation, indicating that the model may benefit from incorporating methylation signals into its predictions.
